# The diploid reference genome of a human embryonic stem cell line

**DOI:** 10.64898/2026.03.26.714432

**Authors:** Ivana Pačar, Matteo Tommaso Ungaro, Ying Chen, Hamza Dallali, Jack A. Medico, Prajna Hebbar, Mark Diekhans, Elena Di Tommaso, Margarita Geleta, Patricia P. Chan, Todd M. Lowe, Jennifer Balacco, Nivesh Jain, Francine Ackerman, Michela Mochi, Alexander G. Ioannidis, Niranjan Sawarkar, Kasandra Diaz, Kavitha Krishna Sudhakar, Joseph E. Powell, Miten Jain, Alessandro Rosa, Gist F. Croft, Andrea Tanzer, Erich D. Jarvis, Giulio Formenti, Sofie R. Salama, Simona Giunta

**Author notes:** These authors contributed equally (Ivana Pačar, Matteo Tommaso Ungaro, Ying Chen). Senior authors (Gist F. Croft, Andrea Tanzer, Erich D. Jarvis, Giulio Formenti, Sofie R. Salama, Simona Giunta).

## Abstract

Advances in DNA sequencing and assembly technologies are spurring a shift from haploid reference genomes to sample-specific diploid assemblies. Here, we generated the first telomere-to-telomere (T2T) diploid reference for the widely used human embryonic stem cell (hESC) line, H9 (WAe009-A). This haplotype-resolved assembly is highly accurate with comprehensive annotation of genes, segmental duplications, methylation, and chromatin conformation. Pangenomic and phased-locus inference point to H9’s mixed ancestry with a predominant European component. H9-specific genomic features include near-perfect telomeres ∼1.65-fold longer than other T2T assemblies, consistent with telomerase activity during pluripotency; chromosome 17 inversions that can predispose offspring to neurological syndromes; and expansions of ncRNA clusters, with overall genomic stability maintained despite extensive culturing. Mapping multi-omic datasets to the genome, we demonstrate the power of this resource for allele-specific, high-precision transcriptomic, genetic, and epigenetic analyses, with far-reaching implications for human development and disease.

## INTRODUCTION

Technological and methodological advancements over the past several years have enabled us to generate Telomere-to-Telomere (T2T) genomes. The first complete human genome was published in 2022.^1^ It was derived from a haploid cell line of a molar pregnancy containing no maternal and two copies of the paternal genome to simplify the assembly process, as there were essentially no haplotypes to phase. More recently, diploid near-T2T assemblies have been generated for human genomes.^2,3,4,5,6,7^ For most experimental biologists, a linear reference genome, no matter how complete, fails to encapsulate the specific variation of the model system under study. Hence, there is a need for diploid reference genomes that match widely used cell line models.^8^ Furthermore, tissue representation in chromosome-level assemblies currently available is largely lacking. Indeed, most high-quality reference human genomes are derived from immortalized peripheral blood lymphocytes from donors, with one exception for the assembly of a cell line derived from the epithelium of the human retina.^7^

Here, we present the first T2T diploid genome reference genome assembly of an (hESC) line, H9 (WAe009-A). H9, one of the first five hESC lines derived by Thomson, *et al.*,^9^ is a female line that is highly stable and easy to maintain in a pluripotent state. This resulted in a uniquely rich historical data ecosystem and ongoing utility for modeling human development and disease, particularly in neuroscience and cell therapy trials.^10^ As of 2016, it has been used in 46% of all peer-reviewed publications on hESCs^11^ and is the most commonly requested stem cell line, as reported by the National Stem Cell Bank^12^ and WiCell. Importantly, H9 is registered in the European Human Pluripotent Stem Cell Registry (hPSCreg^®^), with verified ethical provenance and compliance with European Commission guidelines, ensuring transparent and responsible use of human pluripotent stem cells in research, and is currently the only hESC line approved for clinical use in Europe. The advent of human-induced pluripotent stem cells (iPSCs)^13,14^ almost two decades ago opened the door to studying individual human variation in stem cell models. However, because H9 is a true hESC, it lacks the reprogramming genetic changes and epigenetic memory that can be confounding variables in iPSC studies.^15^ Thus, widely used and well-understood hESC lines like H9 serve as workhorse experimental models, and continue to be the gold standard for pluripotent studies and the development of differentiation protocols.^10,16,17^

Our T2T H9 genome assembly used state-of-the-art approaches, including high-coverage Pacific Biosciences, Oxford Nanopore Technologies long-read sequencing, and high-throughput chromosome conformation capture (Hi-C) data, to generate a chromosome-level diploid assembly spanning all centromeres, with high base-level and structural accuracy.^2^ We also provide haplotype-resolved annotations of coding and non-coding elements, as well as centromeres, telomeres, and other repeats using multiple orthogonal tools.^18,19^ We also performed chromosomal segment-level ancestry inference to estimate continuous ancestry along H9’s two haplotypes. As an additional resource, we provide publicly available methylation information, transcriptome, and chromatin-accessibility data aligned to the new reference, which showcase the power of combining a haplotype-resolved genome with functional genomic data. These data are publicly available as a UCSC Genome Browser track hub,^20,21^ providing the human stem cell research community with the genome as a ready-to-use resource. Ultimately, this H9 diploid reference constitutes a shift from a generalized reference model towards a high-precision tool required to disentangle allele-specific gene regulation. By providing a cell-line matched reference that eliminates reference bias, this work establishes a new foundation for high-precision genomic and epigenomic studies in one of the most widely used models of human development and disease.

## RESULTS

### A near-complete diploid reference genome of a human embryonic stem cell

The reference genome from the H9 hESC line was generated using a combination of Pacific Biosciences (PacBio) HiFi reads (coverage 75×, **Figure S1**), Oxford Nanopore Technology (ONT) R10 ligation reads (coverage 123×, including 47× >100 Kbps), and Arima high-throughput chromosome conformation capture (Hi-C) long-range information (coverage 87×). Two genome assembly strategies were attempted using Verkko v2.2.1. The first assembly (asm1) used HiFi reads for graph construction with ONT reads for graph resolution; the second (asm2) incorporated HiFiasm-corrected ONT reads into the graph construction (**Figure 1A**). Each method yielded 27 and 31 T2T chromosomes, respectively (**Figure S1**). Asm2 contigs complemented asm1 in regions where it is known that sequencing dropout of PacBio HiFi in GA-rich regions generates assembly gaps.^1^ But, asm1 is more accurate at the base pair level than asm2 owing to the significantly higher accuracy of PacBio HiFi reads over ONT-corrected reads.^1^ Therefore, the final diploid assembly (H9v1.0) was constructed by preferentially selecting T2T chromosomes from asm1 (27 T2T chromosomes) whenever present, and for the remaining 9 cases, asm2 chromosomes were used as T2T assemblies. A representation of the remaining 10 chromosomes was generated by manual graph curation and resolution in Verkko (see **Methods**, and **Table S1**).

**Figure 1.**
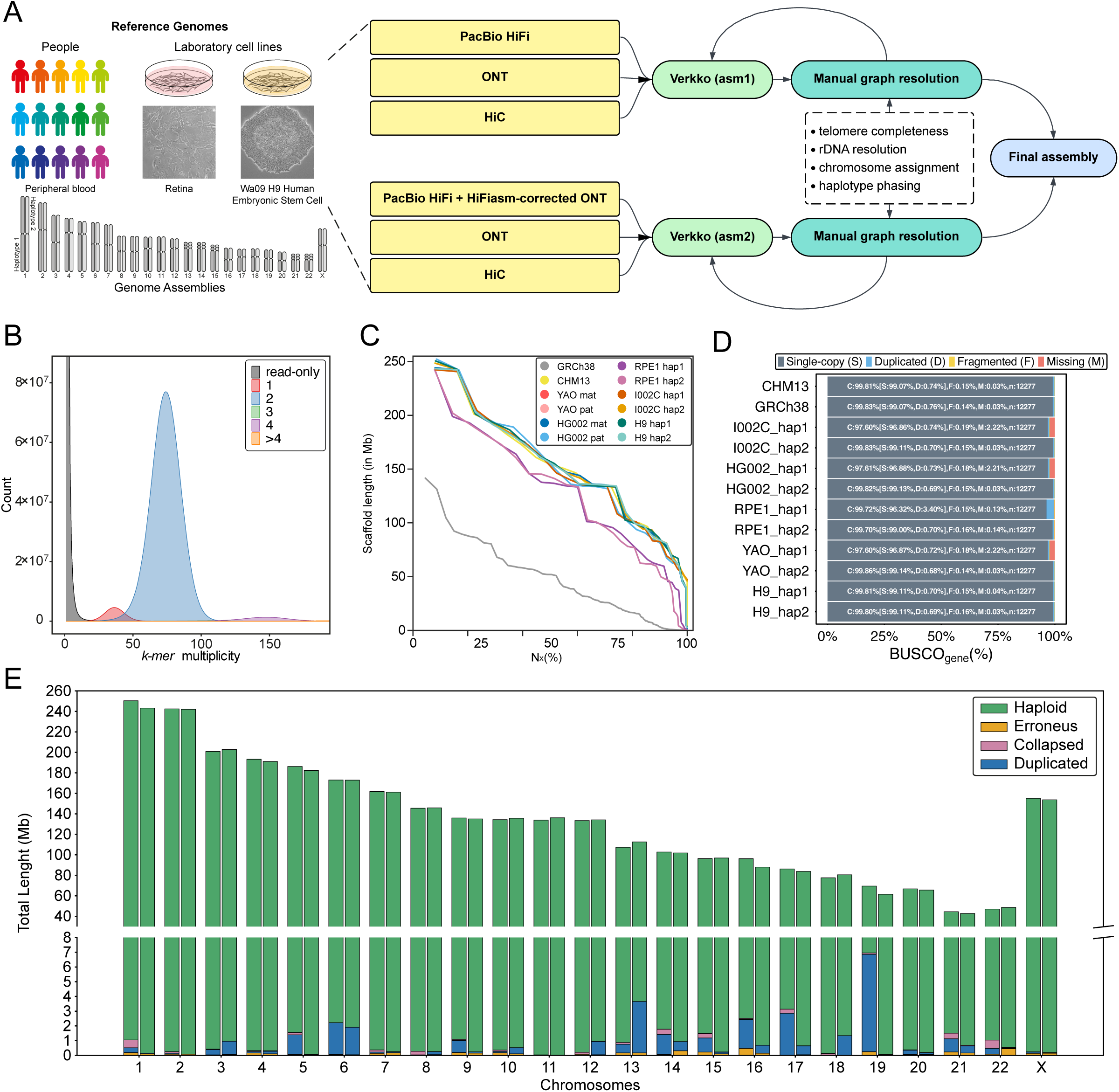
Assembly Workflow with relevant metrics and QVs. (**A**) Bioinformatic workflow of the T2T genome assembly for the human embryonic cell line H9. (**B**) Merqury assembly spectra plot showing *k-*mers found in hap1 and hap2. (**C**) Assembly contig Nx of H9 compared to other human reference genomes in GRCh38 and CHM13, respectively. (**D**) BUSCO gene completeness for 12 T2T haplotype genomes. (**E**) Bar plot showing the amount of sequence (in Kbps) in H9 assembly as erroneous (orange), collapsed (red), duplicated (blue) or haploid (structurally correct, green) classified by Flagger.

### Quality control of the H9 genome

The H9v1.0 assembly was evaluated using various quality control (QC) tools and strategies. We first assessed the alignment of HiFi and ONT reads across the entire genome using NuqFreq and NucFlag v1.0.0a2. Coverage plots showed an overall homogeneous distribution across chromosomes in both haplotypes (**Figure S2**). Some regions exhibited coverage anomalies, such as dropout in chromosome 9 and an increase in chromosome 16 due to the large satellite arrays near the centromeres (**Figure S3** and **S4**), as previously reported.^1^ Some short arms of acrocentric chromosomes (SAACs) in chromosomes 13, 14, 21, and 22 (hap2 only) showed an increase in coverage (**Figure S2-4**) due to the collapsed repetitive rDNA arrays. Merqury *k*-mer based QC yielded a quality value (QV) of 63.6 for haplotype 1 and 66.1 for haplotype 2, well within the range observed for other high-quality human assemblies, and 99.87% completeness for both haplotypes. *K*-mer spectra profiles revealed a multiplicity count consistent with a near-complete assembly, with no detectable duplications (**Figure 1B** and **Figure S1C-D**). Contig Nx curves show that, similar to near-complete human assemblies such as CHM13v2.0^22^ and HG002v1.1,^6^ H9v1.0 is essentially gapless, except for the rDNA arrays on the acrocentric chromosomes (contig N50=155.2 Mbps for hap1; 153.7 Mbps for hap2; **Figure 1C** and **Table S2**). The contiguity of H9v1.0 also exceeds previously released cell-line assemblies, such as RPE1v1.1^7^ (contig N50=136.6 Mbps for hap1; 141.0 Mbps for hap2), placing H9v1.0 among the most contiguous cell-line assemblies to date.

Next, we evaluated the presence of conserved single-copy ortholog genes using compleasm.^18^ The analysis identified 99.11% single-copy complete genes in haplotypes 1 and 2, with a small difference in duplicated genes (0.7% haplotype 1; 0.69% haplotype 2). The missing genes are 0.04% and 0.03%, while the fragmented ones are 0.15% and 0.16% for haplotype 1 and haplotype 2, respectively (**Figure 1D** and **Table S3**). HiFi read alignments analyzed using HMM-Flagger^23^ (see **Methods**) show that of the total 6.06 Gbps diploid assembly, 6.02 Gbps (99.29%) was classified as a reliable haploid sequence, with haplotype 2 displaying a slightly higher proportion of reliable sequence (3.00 Gbps, 99.51%) compared to haplotype 1 (3.01 Gbps, 99.08%) (**Figure 1E**). Regions flagged as assembly errors were rare, totaling 5.45 Mbps (0.09%) and distributed evenly across haplotypes (2.87 Mbps in hap1; 2.58 Mbps in hap2). We observed haplotype-specific differences in structural anomalies: potentially falsely duplicated regions accounted for 33.43 Mbps (0.55%) of the total assembly, with a somewhat higher burden in haplotype 1 (21.38 Mbps, 0.70%) compared to haplotype 2 (12.05 Mbps, 0.40%). Potentially collapsed regions totaled 4.10 Mbps (0.07%), driven primarily by 3.86 Mbps (0.13%) in haplotype 1, compared with only 0.24 Mbps (0.01%) in haplotype 2. We used the coordinates of these regions to define a low-confidence annotation track in the final assembly. Altogether, the H9v1.0 represents a reference-quality, near-complete assembly to support genome-wide functional analysis with precision.

### Pangenome and ancestry analysis confirm genome quality and reveal H9 origins

As further validation of the H9v1.0 assembly, we evaluated its clustering relative to other high-quality genomes from the Human Pangenome Reference Consortium (HPRC)^24^ release. We selected the 44 individuals from the initial release,^24^ for which recent *de novo* assemblies were made publicly available by the HPRC using methods comparable to those employed for the H9v1.0 assembly, and determined its position in principal component (PC) space computed from the pangenome graph structure (see **Methods**); we included in this analysis the two human references GRCh38 (the main linear incomplete reference) and CHM13v2.0 (the T2T haploid reference). As expected, the second principal component discriminates between more contiguous and complete genomes (on the left in **Figure 2A**) and GRCh38, the latter largely missing centromeric HOR arrays and fragmented in several contigs (gray square on the right in **Figure 2A**). The H9 genome groups with the HPRC samples and CHM13 (**Figure 2A**), indicating better alignment in graph space across these high-quality assemblies.

**Figure 2.**
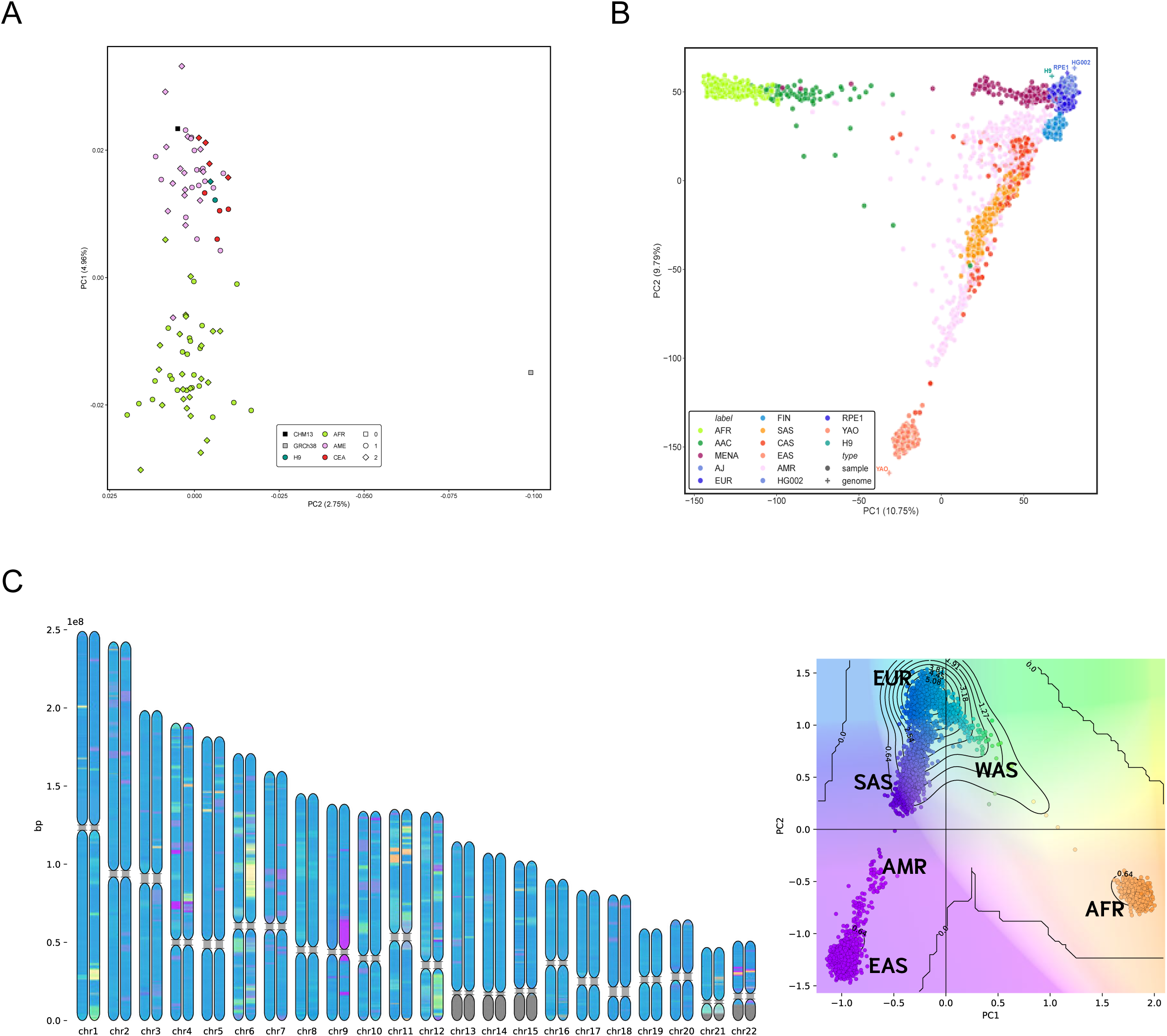
Pangenome PCA and ancestry analyses of the H9 genome. (**A**) Pangenome-based PCA scatterplot of the first release of the HPRC samples (44 total) with the addition of the two human references in GRCh38 (gray) and CHM13 (black) as well as the H9 genome (teal). Shapes indicate whether the sample is an haploid reference (0), or a diploid genome (1: haplotype one, 2: haplotype two); colors distinguish samples from different population groups as reported in the 1KG dataset. (**B**) Genomic ancestry of the H9 cell line inferred based on the GenoTools panel. (**C**) Point cloud local ancestry inference (PCLAI) for the phased H9 genome using the combined 1KG and HGDP reference panel; local ancestry assignments along each autosome for both haplotypes (left), and the corresponding PCA contour plot of the point cloud placement of H9 chromosomal segments relative to the major continental reference clusters (right), indicating a mosaic of predominantly European-like segments with substantial West Asian-like segments.

H9 ancestry had been previously inferred from a limited number of single nucleotide polymorphisms (SNPs) using short-read technology.^25^ We leveraged our whole genome data to define and further refine the estimation of the major ancestral components for this genome across phased haplotypes. We initially used GenoTools, a method applied to various large disease genetics initiatives, including NIH’s Center for Alzheimer’s and Related Dementias (CARD) and the Global Parkinson’s Genetics Program (GP2).^26^ GenoTools assigns a population label to a query according to a pre-computed set of genetic distances integrating samples from the 1000 Genomes Project (1KG),^27^ the Human Genome Diversity Panel (HGDP),^28^ and an Ashkenazi Jewish reference panel (AJ).^29^ In line with the previous SNP chips analysis, H9 clusters in proximity to European populations. We found it to be particularly close to Ashkenazi Jewish (AJ) samples present in the dataset, supporting the evidence for this cell line being derived from an IVF oocyte donated from a medical center in Israel.^30^ We also used GenoTools to place three T2T genome samples, HG002, I002C, and RPE1 – for which the population background is known – and found they cluster in their appropriate groups (**Figure 2B**), confirming the robustness of this analysis.

Next, we increased resolution using a complementary set of ancestry analyses. We first applied point cloud local ancestry inference (PCLAI)^31^ using a combined 1KG^27^ and HGDP^32^ reference, which assigns continuous PCA coordinates along phased H9 haplotypes (see **Methods**). PCLAI indicated that H9 has predominantly European-like ancestry (EUR), albeit with substantial West Asian-like (WAS) chromosomal segments and little evidence for African-like ancestry (AFR), consistent with point cloud placement of haplotypes along the EUR-WAS continuum (**Figure 2C**). To contextualize this result at a finer scale, we performed PCA projection using 300000 max-heterozygosity sites selected with PLINK2.0^33^ onto a Mediterranean, Levantine, and North African subset of 432 samples from the 1KG and HGDP combined panel. In this analysis (**Figure S5**), H9 projects between Bedouin, Palestinian, and Druze references on the one side and Italian and Spanish on the other, reinforcing the PCLAI analysis of an individual having ancestry from both Europe and West Asia, in particular the Levant. We then performed a 3-population test using snputils^34^ F_3_ (YRI; H9, reference) using subset references; the estimates agree with the above conclusions, indicating the greatest shared genetic drift with Southern European and Levantine references, with lower affinity to North African references (**Table S4**).

### Whole-genome annotations highlight haplotype-specificity

We next sought to annotate the genes present genome-wide and identify differences in each haplotype. Gene annotation based on RNA-Seq and comparative genomics^35^ identified 20098 protein-coding genes, 15445 pseudogenes, and 44273 non-coding RNAs, of which 35354 are lncRNAs in haplotype 1, and 20148 protein-coding genes, 15480 pseudogenes, and 44356 non-coding RNAs of which 35382 are lncRNAs in haplotype 2 (**Figure 3B**). This includes 3043 and 3052 genes detected on the X chromosomes in haplotypes 1 and 2, with 891 and 899 of them being protein-coding, respectively. One-to-one orthologs of 19925 human genes were identified in haplotype 1 and 19943 in haplotype 2, which represent 99.07% and 99.16% of all previously identified protein-coding genes. After computing gene copy numbers across both haplotypes, we identified genes present on only one haplotype. After filtering to reduce false positives from closely related genes and excluding sex chromosomes, we identified 7 haplotype 1-only and 8 haplotype 2-only genes (**Table S5**.) Excluding genes absent from one haplotype, we identified 20 genes with higher copy numbers on haplotype 1 and 31 on haplotype 2. These numbers are in line with observations from various projects, including HPRC^24^ and HG002^6^, and consistent with haplotype-specific regions of duplication and variation (see **Identification of segmental duplications specific to H9**).

**Figure 3.**
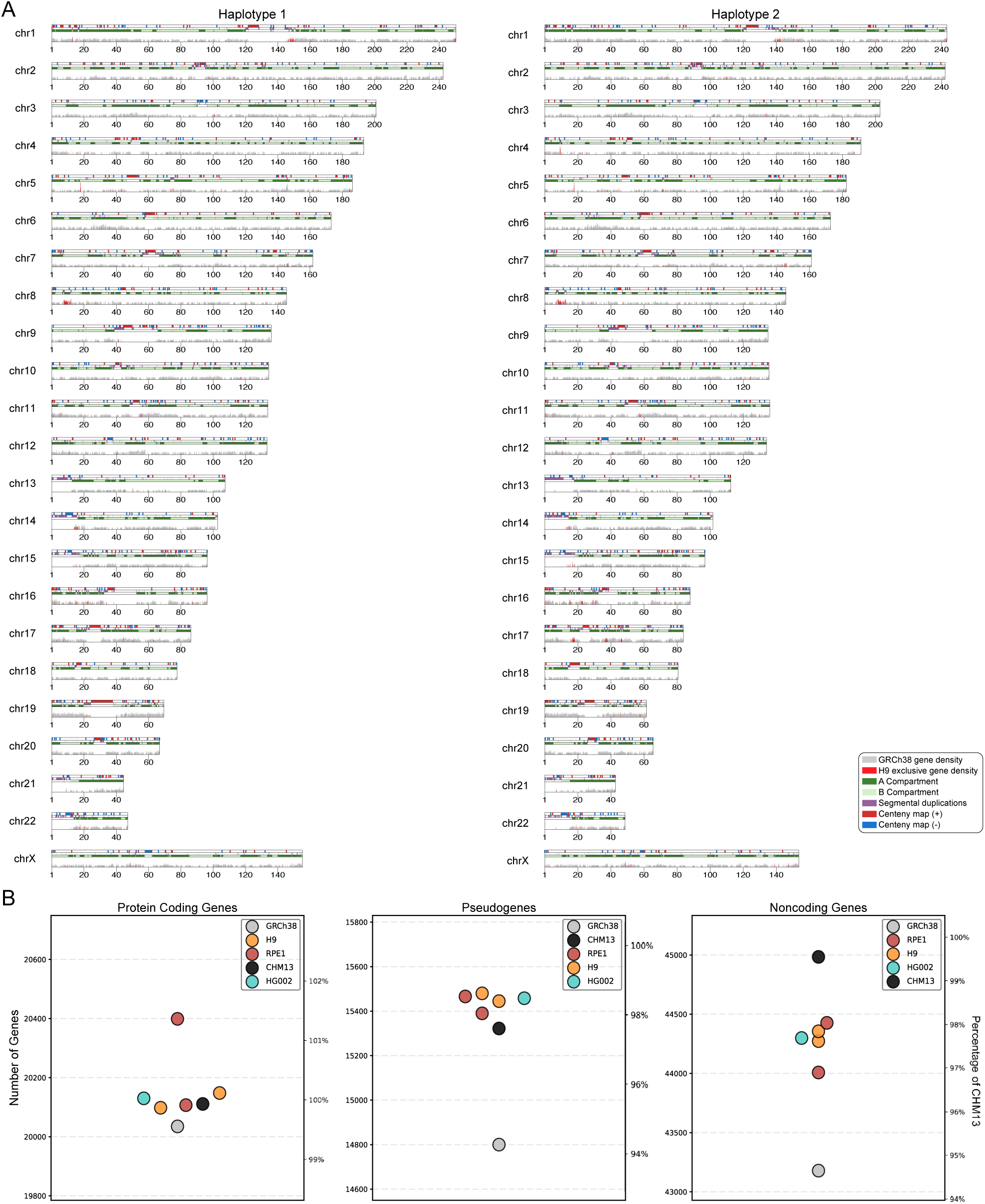
Whole-genome annotation and gene content comparison of the H9 assembly. (**A**) Karyotype plots illustrating the genomic landscape of chromosomes 1–22 and X for H9 haplotype 1 (left) and haplotype 2 (right). Tracks visualize gene density profiles, specifically highlighting GRCh38-annotated gene density (grey) and H9-exclusive gene density (red). Major structural features are annotated, including centromeres (red and blue), segmental duplications (purple), and the A/B compartments (green shades). (**B**) Comparative analysis of gene annotation completeness across various high-quality human genomes, including GRCh38, CHM13, H9, HG002, and RPE1. Plots display the total counts for protein-coding genes, pseudogenes, and non-coding genes, with the right y-axis representing the percentage relative to the CHM13 reference.

To assess differences in gene content and density between GRCh38 and the H9v1.0 assembly, the GRCh38 GENCODE V48^36^ protein-coding genes were projected to the H9 haplotypes via the minimap2^37^ genomic alignment. The analysis shows the intermittent presence of H9-exclusive genes across both haplotypes, overlaid on GRCh38 gene density (**Figure 3A**). Regions with a noticeably higher proportion of H9-exclusive gene density (such as on the p-arm of chromosome 8 in both haplotypes) have been verified to have consistent Hi-Fi read coverage (**Figure S3**), implying a lack of assembly errors and reflecting a gene content increase.

Next, we sought to establish the chromatin state of the H9 genome by determining the A (euchromatin) and B (heterochromatin) compartments. Using Hi-C reads, the A/B compartment analysis of the H9 genome was displayed as a characteristic checkered pattern on top of the gene density block (**Figure 3A**). In addition to the chromatin analysis, the karyotype plots show segmental duplications, determined with SEDEF,^38^ and the Centeny map obtained marks the placement and orientation (blue for forward and red for reverse complement sequence within the same strand) of CENP-B boxes obtained using the Genomic Centromere Profiling (GCP) pipeline.^39^ For both tracks, we observe largely conserved patterns between the two haplotypes.

Furthermore, we conducted an assessment of the small RNA gene repertoire in H9v1.0. Using BLAST, we aligned microRNA precursors from GRCh38 to CHM13v2.0 and to the H9 haplotypes, and assessed copy numbers for each microRNA family. For the majority of families (1367, 95.1%), the number of family members was consistent between GRCh38 and both H9 haplotypes, as well as CHM13. For 67 families, we detected copy number variations (CNVs) in either haplotype 1 or haplotype 2 of the H9v1.0 assembly (**Table S6**). The most striking case is the microRNA family MIR506, which forms a cluster of 20 individual microRNAs on chromosome X. This microRNA cluster has been associated with various types of cancer^40^ and spermatogenesis.^41^ The members of this family reside in two different lncRNA host genes (ENSG0000029678, ENSG00000287017), suggesting independent transcriptional regulation. The three copies of the MIR506 family member MIR514A are associated with lncRNA ENST00000658633.1 (ENSG00000287017.2). MIR514A1 is downstream of the last exon, MIR514A2 resides in intron 4, and MIR514A3 resides in intron 2. The locus is rich in segmental duplications (**Figure 5D**), suggesting the cluster has arisen through local tandem duplications.^42^ In H9 haplotype 2, we observe two additional copies of MIR514A3 in exon 2, and in haplotype 1 the cluster further increases in size by another tandem duplication, consisting of 6 microRNAs in total with 4 copies of MIR514A3 in intron 2 (**Figure 5D** and **Figure S6**). Local duplications are fully covered by sequencing reads indicating these are unlikely to pertain to assembly errors (**Figures S7C-D**).

We identified tRNA genes in H9 using tRNAscan-SE^43^ and cross-mapped them onto CHM13v2.0 using whole-genome sequence alignments. Similar to CHM13, we find additional tRNA genes in the H9 haplotypes that were not originally identified in GRCh38, resulting in an overall increase of 11% more genes compared with the most commonly used reference genome for tRNA studies (**Table S7A**). More than 90% of these are high-scoring and exhibit canonical tRNA gene features, thus suggesting only a limited number of pseudogenes or misassigned repetitive elements in our H9 tRNA annotations. Almost all the gene gains are located in a highly repetitive region on chromosome 1 supported by full and homogenous read coverage (**Figure S7A-B** and **Table S7B**). They are arranged as multiple sets of five tRNA genes (tRNA^Glu(CUC)^, tRNA^Gly(UCC)^, tRNA^Asp(GUC)^, tRNA^Leu(CAG)^, and tRNA^Gly(GCC)^), with each set spanning over 7 Kbps to form a tandem repeat unit. Previous studies have demonstrated that the number of this tRNA tandem repeat varies across individuals in the 1KG cohort.^44^ Consistent findings were observed when comparing CHM13 and the H9 haplotypes. While CHM13 has 22 repeat units, H9 has 20 and 16 repeat units in haplotype 1 and haplotype 2, respectively, highlighting the copy number variation inherited from each parent (**Table S7C**). Interestingly, H9 also harbors some tRNA genes (7 in hap1; 4 in hap2) that are absent from CHM13 (**Table S7C**). While these extra genes that reside in segmental duplication regions in both haplotype 1 and haplotype 2 are mostly low-scoring, tRNA^Gly(CCC)^ on chromosome 1 of haplotype 1 represents an extra copy of three identical tRNA^Gly(CCC)^ genes found on chromosome 1 of CHM13 and H9. When taking a closer look on the genes that are found in both CHM13 and H9, we noticed that the sequences of 7.5% (53 out of 705) and 8.5% (57 out of 671) tRNA genes in H9 haplotype 1 and haplotype 2, respectively, are different from those in CHM13 due to single nucleotide variants (SNVs). Single nucleotide variations (SNVs) also produce sequence differences in 44 tRNA genes when comparing the two H9 haplotypes. The functional consequences of these inter-haplotype differences will depend on the position of such SNVs within tRNA genes and the epigenetic state of those genes, yet this unexpected variability may drive allele-specific expression at translational level.

### Centromere annotation points to longer satellite arrays in hESC

The availability of fully resolved centromeres in the H9 genome enables, for the first time, a detailed investigation into their nature and organization in a human embryonic stem cell line. We next compared these centromeres to those of Retinal Pigmented Epithelial 1 (RPE1), whose genome has been recently assembled^7^ and characterized in detail,^8^ and which has a different tissue of origin (retina pigmented epithelium). We annotated the H9 centromeres using multiple orthogonal methods, each providing a different layer of information; specifically, we employed Hum-AS-HMMER for AnVIL (https://github.com/fedorrik/HumAS-HMMER_for_AnVIL), which outputs complete higher order repeats (HORs) and annexed superfamily annotation, dna-nn^45^ that represents AATTC repeats and alpha satellites, and the Genomic Centromere Profiling (GCP) pipeline,^39^ based on CENP-B boxes orientation and outputting distance values between motifs which can provide architectural information pertaining to centromere organization in T2T genomes and chromosome-level assemblies. We found the three to be highly concordant for annotation of the active region of the centromere, within which the kinetochore binding occurs, with Hum-AS-HMMER for AnVIL providing comprehensive alpha satellite families and superfamilies assignment. Therefore, we used this annotation to compare centromere lengths between the two H9 haplotypes, and with the RPE1 cell line. On average, we observed that in both the RPE1 and H9 genomes, haplotype 1 has slightly longer centromeres than haplotype 2. Interestingly, the majority of H9 chromosomes have longer centromeres compared to RPE1, with ten H9 centromeres being on average 18.8% longer than the corresponding RPE1 counterpart; this was determined by considering those chromosomes with longer centromeres in both haplotypes. To provide an estimate, this analysis excluded chromosome 19, where we found a centromere in H9 haplotype 1 that is larger in size than currently reported centromere length distributions across humans^46^, with a total length of about 13.4 Mbps. (**Figure S8**).

To investigate whether the long chromosome 19 haplotype 1’s centromere is an assembly artifact, we used several orthogonal approaches. With dna-nn alpha satellite annotations, we calculated the SNV density between the two H9 haplotypes, showing variation to a higher extent (0.53%) than what was expected for centromeres (**Figure 5A**), where haplotype-to-haplotype alignment may be affected by the centromeres’ size difference. We then combined evidence from the GCP pipeline with NuqFlag^46^ v1.0.0a2 used as a means of assembly quality validation, with particular attention to centromeric regions. According to one of the GCP pipeline outputs, the Centeny map (**Figure S9**), chromosome 19 haplotype 2 has an unexpected gap in the CENP-B high-density domain of the active centromere, indicating a possible assembly error. However, GCP pipeline models 1 and 2, which present visual renderings of the centromere structure (**Figures S10** and **S11**), show homogeneous patterns between the two haplotypes, with color profiles aligned to the pattern expected for chromosome 19 in humans, in spite of minor regions of previously undocumented organization present on both haplotypes (**Figure S12**). Finally, we assessed the centromere regions with NucFlag (**Figure S13**), which showed no evidence of assembly errors, pointing to this being the longest assembled centromere at over 13 megabases of DNA.

### Telomere annotation in H9 and other human references

Teloscope (Medico *et al.*, in preparation) identified 692.6 Kbps of telomeric DNA in the diploid H9v1.0 assembly, mainly composed of canonical CCCTAA/TTAGGG repeats (638.5 Kbps, 94.8%). We annotated 91/92 telomeres on all chromosome ends of H9v1.0 (**Figure 4A**), except for chromosome 22 p-arm in haplotype 2, which only contains a short tract of telomeric variant repeats. Assembled telomere length averaged 8.01 ± 4.49 Kbps in haplotype 1 (median 6.93 Kbps) and 7.20 ± 4.39 Kbps in haplotype 2 (median 5.94 Kbps), with no significant difference between haplotypes (Wilcoxon V = 439, p = 0.382). These assembly-derived telomere lengths were consistent with prior Telomere Restriction Fragment (TRF) estimates for H9 cells (7.8 to 13.2 Kbps).,^47,48^

**Figure 4.**
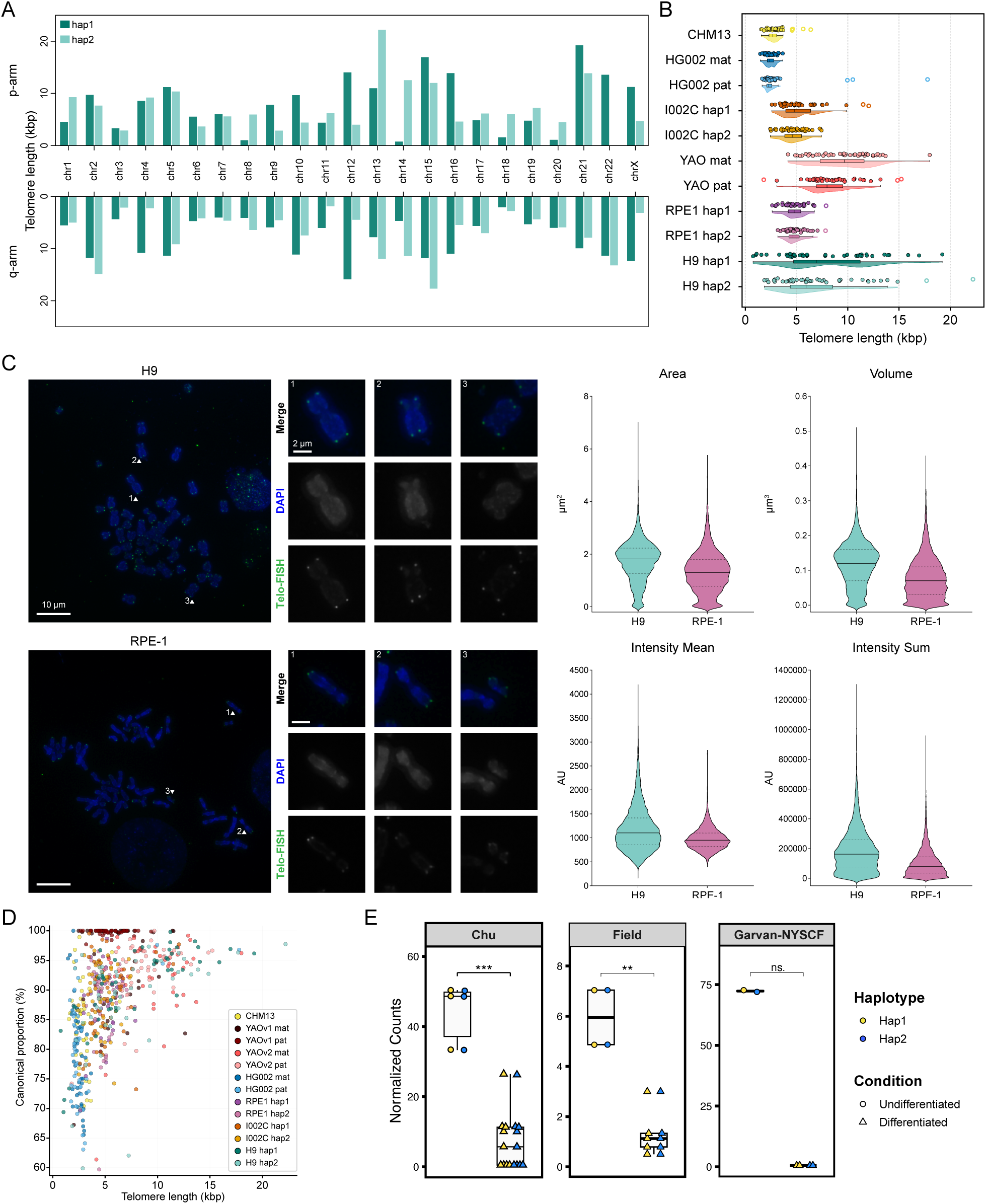
Telomere annotation and validation of the H9 assembly. **(A)** Assembly-based telomere lengths by haplotype and chromosome arm annotated with Teloscope. **(B)** Comparison of telomere lengths of high-quality human (CHM13, HG002, YAO, and I002C) and cell line genomes (RPE1 and H9). **(C)** Telomere FISH (left) and quantification of telomere length (right) in each cell line. We analyzed 35 metaphase spreads containing 120-200 measurable telomere foci each. **(D)** Proportion of canonical telomeric repeats (CCCTAA/TTAGGG) against telomere length. **(E)** TERT expression across undifferentiated and differentiated hESC cells from both unpublished and published datasets.

Telomeric repeats remain challenging to fully and correctly assemble even with long reads,^49^ therefore caution should be exercised in drawing biological conclusions when evaluating them. We aimed to validate chromosome ends by combining evidence from NucFlag read support using HiFi and ONT data. Using an overlap-based classification (**Supplementary Information**), 90.7% of telomeric DNA was supported by at least one sequencing technology, and only 0.21% showed errors in both (**Figures S14-17**). Flagged bases (9.3%) consisted predominantly of repeat-context (6.4%) and mismatch-signal (2.7%) (**Figure S18**). Among acrocentric chromosomes, the percentage of telomeric bases deemed correct by HiFi was markedly lower than at metacentric or submetacentric ends (Kruskal-Wallis H=26.201, p=2.0442e-06) while ONT maintained near-complete support (**Figure S18**). We manually verified the presence of ONT and HiFi reads anchoring in unique subtelomeric sequences and extending into the telomere. As expected for chromosome ends, we observed a high coverage at the subtelomere boundary followed by a gradual drop toward the distal end. Overall, ONT and HiFi reads complemented each other since the former resolved bases at distal tips, while the latter improved mismapping calls.

Compared to other near-complete human genomes, telomeres in both H9 haplotypes were longer than those reported in current assemblies, including CHM13, HG002, and RPE1 (**Figure 4B**). To orthogonally validate the difference in telomere length, we used fluorescence in situ hybridization (FISH)^50,51^ and quantitative microscopy to compare telomere length in H9 versus RPE1 cells (**Figure 4C**), which have been reported to have short telomeres.^7^ Three-dimensional (3D) images acquired with wide-field deconvolution microscopy were analyzed to calculate telomere fluorescence intensity mean, sum, area, and volume (**Methods**). Despite the expected intra-chromosomal and metaphase variability (**Figure S19**), we observed that the telomeric probe signal was higher on average in H9 compared to RPE1 for all metrics analyzed (**Figure 4C**), validating the presence of longer telomeres in this cell line and confirming the results found using the assemblies.

### TERT expression is associated with hESC-specific lengthening of near-perfect telomeres

We also compared the telomeric hexamer composition of the H9v1.0 assembly with those of recently generated human assemblies (CHM13v2.0, HG002v1.1, I002Cv0.7, YAOv1.0 and v2.0, and RPE-1v1.1). YAOv1^3^ telomeres visibly cluster as short, highly canonical (TTAGGG-rich) points from decoy telomeric-like sequences that were later resolved in YAOv2^5^ (**Figure 4D**), indicating that our analysis successfully intercepted differences in telomere length and base-pair composition. However, H9 telomeres, and partly YAOv2, showed broad dispersion in telomere length, which was positively associated with the canonical TTAGGG proportion (**Figure 4D**). This pattern suggests that longer, near-perfect telomeres are enriched for recently added canonical repeats driven by telomerase activity in embryonic stem cells.^44,51^ To validate this, we analyzed TERT expression in three publicly available RNA-seq datasets from H9 differentiation to dorsal forebrain organoids,^52^ definitive endoderm^53^ as well as an unpublished dataset from ESC, midbrain progenitors, and astrocytes (this study). Due to differing time points across studies, we compared the cumulative expression between undifferentiated and differentiated samples. In all cases, TERT expression was consistently higher in the undifferentiated H9 cells, which was the cell state used to generate the samples for this assembly (**Figure 4E**). These results support H9 as a model of developmental TERT regulation and confirm the pluripotent state of the line, with actual evidence of near-perfect telomere elongation due to active telomerase activity in this cell type.

### Genome-wide sequence and structural variation

We took advantage of the H9 diploid genome with phased haplotypes to explore inter-haplotype variation. To investigate regional variation in greater detail, we quantified SNP density in nonoverlapping 10 Kbps windows across the genome (see **Methods**). While the average SNPs density across chromosome arms between haplotypes was 0.11% (one heterozygous marker every 1000 bps), it increased to 0.53% within centromeric regions (active alpha satellite HORs), with the highest peaks in the centromeres of chromosome 19 (5.51%), possibly due to also having the widest difference in length of the centromeres as described above, and the lowest in the chromosome X centromeres (0.01%) (**Figure 5A**). The highly divergent regions (HDRs) are predominantly associated with centromeres (**Figure 5A**), which are also enriched in structural rearrangements. This pattern has been observed across other haplotype-resolved genome assemblies^6,3,7^, largely due to limited meiotic recombination that preserves centromere blocks into intact haplogroups with high genetic diversity between the two parental haplotypes. In total, the H9 haplotypes show 3010848 SNVs, 69 inversions, 1141 translocations, about 1.1 Mbps of insertions and deletions (30–500 bps), and ∼95.6 Mbps of other complex and/or large structural variants (>100 Kbps).

**Figure 5.**
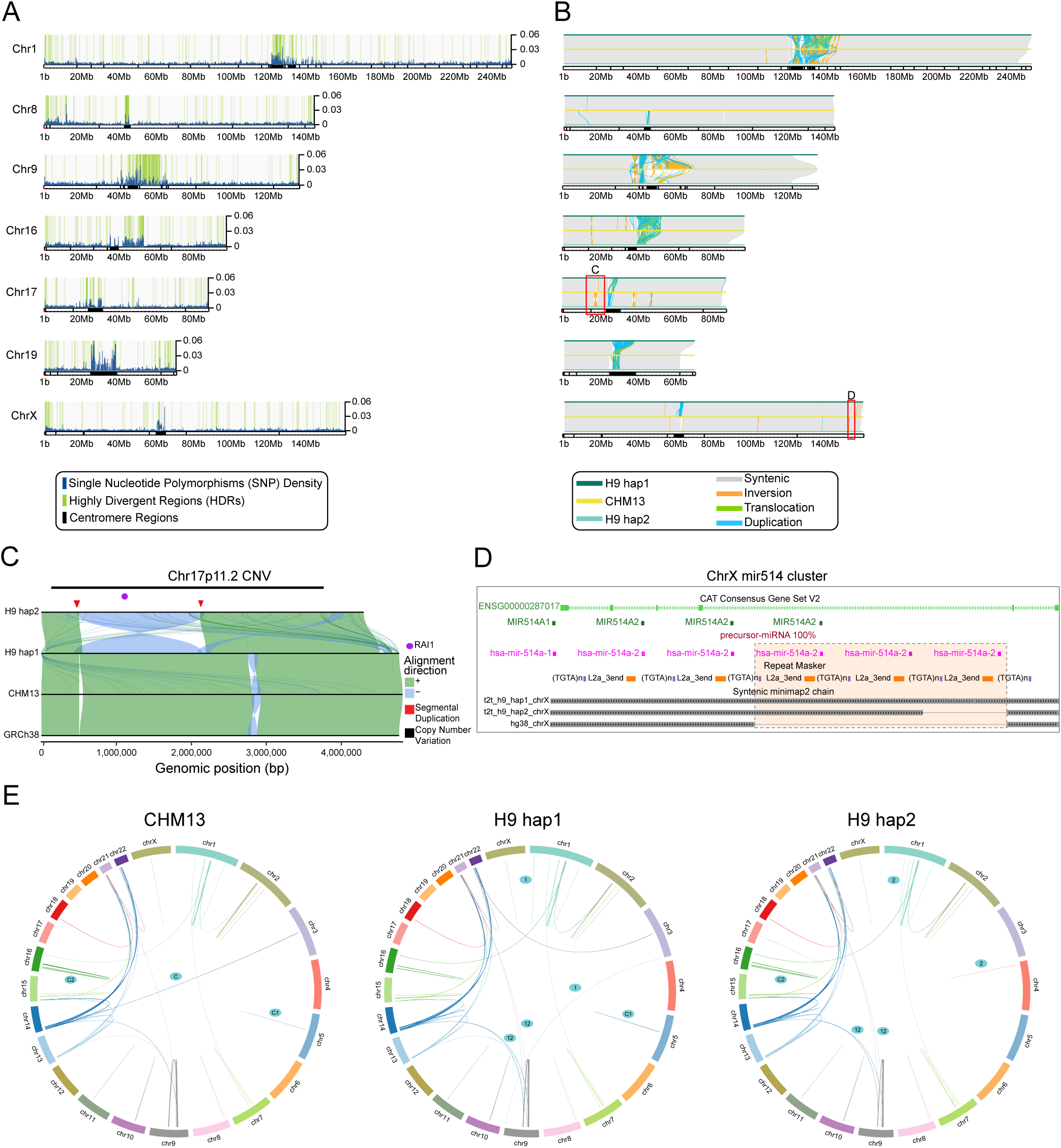
Characterization of H9-specific variants and structural rearrangements. (**A**) Genome-wide distribution of cell-line specific variants across selected chromosomes. Single-nucleotide polymorphism (SNP) densities (blue traces) and highly divergent regions (HDRs, green shading) are shown alongside centromeres (black bars). (**B**) SyRI synteny and alignment visualization highlighting H9-specific genomic features. The top panel illustrates structural relationships between H9 haplotype 1 (hap1), CHM13, and H9 haplotype 2 (hap2), including syntenic (grey), inverted (orange), translocated (green), and duplicated (blue) regions. (**C**) SVbyEye^75^ of Chr17 displaying the p11.2 neurodevelopmental CNV locus with a distinct inversion on haplotype 2 relative to haplotype 1 and the human references GRCh38 and CHM13. (**D**) microRNA cluster mir-514 shows copy number variations and cluster expansion in H9 relative to GRCh38 assembly and other T2T genomes (**E**) Circos plot providing a global summary of inter-chromosomal relationships and structural variation within the H9 assembly, as further validation of assembly consistency and structural specificity.

To better define haplotype differences in H9v1.0 and to compare with CHM13v2.0, we identified their nature and length with SyRI.^54^ Syntenic regions compared to CHM13 show that H9 haplotype 1 is 99.90% syntenic with the CHM13 reference, and haplotype 2 a slightly lower 99.88%. As a consequence, both display a comparable number of variants relative to it: haplotype 1 shows 3124343 SNPs, 66 inversions, and 1847 translocations, while haplotype 2 has 3089128 SNPs, 62 inversions, and 1697 translocations. Only a minimal fraction of either H9 haplotype does not align to CHM13, at roughly 139 Mbps for haplotype 1 and 134 Mbps for haplotype 2, with an average unaligned sequence block of 343 Kbps (**Figure 5B**). Altogether, both between H9 haplotypes and when aligned to CHM13, the chromosomes show long stretches of syntenic regions (**Figure 5A-B**, see **Figure S20** for all chromosomes), validating H9 as a stable model system deprived of major rearrangements or other genomic changes due to culture conditions.

### Identification of segmental duplications specific to H9

Segmental duplications (SDs) are regions of DNA greater than 1000 bps that are repeated on the same or different chromosomes with >90% sequence identity.^55^ They are a source of new gene evolution and risk factors for numerous diseases.^56,57,58^ In cell lines, there is the potential that structural variation arising from cell line establishment and prolonged culture passaging could influence their phenotypes.

Segmental duplications in the H9v1.0 assembly were annotated using SEDEF.^38^ The two haplotypes comprised 5.6% and 5.7% segmental duplications, respectively, with substantial variability in SD frequency across chromosomes (**Table S8**). This is consistent with the 5.6% SD coverage of the CHM13 assembly, excluding chromosome Y, the mitochondrial genome, and the rDNA arrays absent from the H9v1.0 assembly.

To characterize large-scale, recent segmental variation between CHM13 and H9, duplicated regions of at least 250 Kbps with ≥98% sequence similarity were compared. These large, recent duplications covered 1.2% of the genome in both H9 haplotypes, compared to 1.0% in CHM13, and most of these recent SDs are shared between the H9 haplotypes and CHM13 (**Figure 5E** and **Table S9)**. This suggests that the generation and propagation of the H9 cell line have not led to significant expansion or deletion of sequences in segmental duplication regions. Comparing segmental duplication events between assemblies in finer detail is challenging due to overlapping duplication annotations and variability in computational annotations, even across nearly identical DNA regions. Cross-assembly comparison showed that 85.7% of CHM13 segmental duplication bases were shared with the H9 haplotypes, while 78.4% and 79.0% of H9 haplotype 1 and haplotype 2 SD bases, respectively, were shared with CHM13 (**Table S10**). Acrocentric chromosome arms were excluded from this analysis due to the difficulty of generating reliable genomic alignments in these regions. To identify SD bases unique to each assembly, SD annotations were intersected with regions lacking genomic alignments to the other assemblies of at least 500 bp. In a three-way comparison, the H9 haplotypes had 3.5% and 3.4% of their SD bases unique to the assembly, compared to 2.1% for CHM13. When comparing the H9 haplotypes with each other, 7.2% and 6.2% unique SD bases were observed (**Table S11).**

To further evaluate the genomic integrity of H9 as a model for human biology, we compared our assembly against two established datasets of clinically and biologically relevant structural variants. First, we investigated a set of 20 genomic regions previously identified as common CNVs in prolonged human iPSC culture.^59^ Our H9v1.0 assembly showed no evidence of insertions or deletions at any of these 20 loci, which were mapped from the original GRCh37 reference to H9 (**Table S12**).

Next, we cross-referenced the H9 and CHM13 genomes against 83 loci from a systematic review of recurrent CNVs associated with neurodevelopmental disorders.^60^ While both H9 haplotypes lacked CNVs in these loci, we identified four structural variants involving interhaplotype inversions among these loci (**Table S13**). Three out of four of these inversions have been observed in HPRC samples: 17p11.2 in 5 haplotypes, 17q21.31 in 2 haplotypes, and 2q27 in 31 haplotypes.^24^ Notably, we observed a ∼1.6 Mbps inversion at 17p11.2 in haplotype 2 (**Figure 5C**), flanked by segmental duplications. While this specific paracentric inversion is generally considered non-pathogenic, in carriers it may represent a “pre-state” for genomic instability. This risk is illustrated by a documented familial case where a phenotypically normal father carrying this inversion sired a child with a complex rearrangement, resulting in a deletion associated with Smith-Magenis syndrome, which is thought to be driven primarily by mutations in the *RAI1* gene (**Figure 5C**).^61,62^

To investigate the functional impact of these observed structural variations, RNA-Seq reads were analyzed for known causal genes in each of the four CNVs in the RNA-Seq data set from H9-derived dorsal forebrain organoids.^52^ While causal genes for three of four CNVs showed no differential expression between H9 haplotypes, one of five genes implicated in 17q21.31 CNV pathogenesis showed haplotype-specific expression. We observed higher *KANSL1* expression in the neurally differentiated cells in haplotype 1 compared to haplotype 2 (66:34). In addition to *KANSL1*, neighboring genes *CRHR1*, *SPPL2C*, *MAPT*, and *STH* are all implicated in the pathogenesis of 17q21.31 microdeletions,^63,64^ but they did not show evidence of allele-specific expression. However, we cannot rule out the possibility that this is due to a combination of lower expression and/or a lack of haplotype-specific SNPs within the transcripts of these genes (**Figure S21**). This caveat also applies to the other loci where we did not detect allele-specific expression when examining the pathogenic CNV-associated genes affected by the inversions.

The 17q21.31 locus exhibits two distinct haplotypes, which differ by a ∼1 Mb inversion.^65^ The direct haplotype (H1), observed in H9 haplotype 1, is common with an approximate frequency of 0.8 in European ancestry populations. In contrast, the inverted haplotype (H2), observed in H9 haplotype 2, is less frequent, occurring in ∼0.2 of European ancestry populations and <0.09 in East and South Asian populations.^66^ While the major haplotype H1 is associated with increased risk for multiple neurodegenerative diseases, including Alzheimer’s Disease (AD) and Parkinson’s Disease (PD), the less common H2 haplotype predisposes offspring of carriers to the 17q21.31 microdeletion syndrome.^65^ This dual genetic landscape underscores the value of H9 as a robust system for modeling both early-stage neurodevelopmental and late-onset neurodegenerative pathologies.

### Improved haplotype-resolved transcriptomics analysis using the H9 reference

To discern the effects of the reference genome used for read alignment on transcriptomic quantifications, we conducted differential transcript mapping analysis of RNA-Seq data, generated on H9 cells, by comparing the same biological replicates aligned to different reference genomes (H9 hap1 vs. H9 hap2, H9 hap1 vs. GRCh38, and H9 hap2 vs. GRCh38). Indeed, our analysis revealed divergence in gene mapping profiles, mainly due to the reference genome used for read alignment (**Figure S22**, **Tables S15** and **S16**). Observed differences can be attributed to mapping discrepancies rather than expression profile variations, which points out differentially mapped genes (DMGs) instead of differentially expressed genes (DEGs) as previously described.^8^ We found 265 and 274 DMGs when comparing H9 haplotype 1 and haplotype 2 to GRCh38, respectively. Most importantly, we identified 232 DMGs shared between the two H9 haplotypes when using GRCh38 as a reference, including 59 genes not mapped at all to GRCh38 (**Tables S17** and **S18**). These results emphasize the importance of using our diploid H9 reference when analyzing omics data generated from experiments on H9 cells, where certain genes would appear as blind spots to assess differential expression between samples. In addition, 81 genes were differentially mapped between H9 haplotypes, which shows the added value of using the H9 diploid reference in uncovering haplotype-specific differential gene expression (**Table S19**, **Figure S22C**).

Next, we assessed the impact of haplotype-specific SNVs on H9 allele-specific expression by assigning them a genomic feature annotation. We found that the majority of the 3010848 SNVs reside within intronic (53.1%) and intergenic regions (40.2%), while promoter and UTR SNVs comprise only 1.9% and 1.1% of their total number, respectively (**Figure S23**). Among the 81 DMGs between H9 haplotypes, 13 and 21 genes harbor promoter and UTR SNVs, respectively. Notably, 7 DMGs contain both promoter and UTR SNVs (**Tables S20-22**). Some of these genes, such as RPL9, RPS2, and RRP7A, are central to ribosome biogenesis and translational capacity. Notably, the latter encoding for a ribosome subunit, displays a pronounced reduction in mapped reads for haplotype 1 (logFC=−5.04) and contains 12 UTR variants. Similarly, PDPR, encoding a metabolic regulator controlling pyruvate dehydrogenase activity during aerobic glycolysis, shows a similar trend of reduced mapping to haplotype 1 (logFC=−3.03). Promoter SNVs may alter transcription initiation through differential transcription factor binding, whereas UTR ones may influence mRNA stability. Consequently, these results suggest potential allele-specific control of protein synthesis and glycolytic metabolism genes in H9.

Finally, we selected a small panel of genes whose protein dosage and activity, in specific cell types, are directly linked to neurodegenerative diseases: SNCA, LRRK2, and GBA, associated with Parkinson’s; HTT with Huntington’s Disease; and SMN1 with Spinal Muscular Atrophy. When probed for cell-type and allele specific regulation, the majority of RNAseq reads for each gene mapped equally to both haplotypes, which was unsurprising given relatively low divergence, including near homozygosity for GBA1 (**Figure S24A**). Allele-specific expression of LRRK2 was balanced across haplotypes in all cell types, while HTT showed a slight shift to Hap1 in ESC. Notably, while SNCA was expressed evenly in most cell types, astrocytes showed a dramatic switch to Hap2-dominant expression, while SMN1 was dominantly expressed from Hap1 in all cell types (**Figure S24B**). The significance (**Figure S24C**), and magnitude of ASE for SNCA and SMN1 are substantial enough to potentially affect disease risk and phenotypes, especially in the context of natural or genome-engineered heterozygous coding variant risk alleles, or for testing gene therapy approaches. H9v1.0 therefore enables the discovery of these specific regulatory mechanisms, and is an important new tool to more precisely understand risk gene regulation and disease models in future studies.

### Haplotype-specific chromatin accessibility

We leveraged the improved mapping quality using matched reads-reference to investigate chromatin accessibility using an ATAC-Seq dataset from H9 cells during early neuronal differentiation (GSE291907).^67^ We found 90584 open chromatin peaks in H9 haplotypes (**Figure 6A**). As expected, ATAC peaks are predominantly located in intronic, intergenic, and promoter regions (**Figure 6B**).

**Figure 6.**
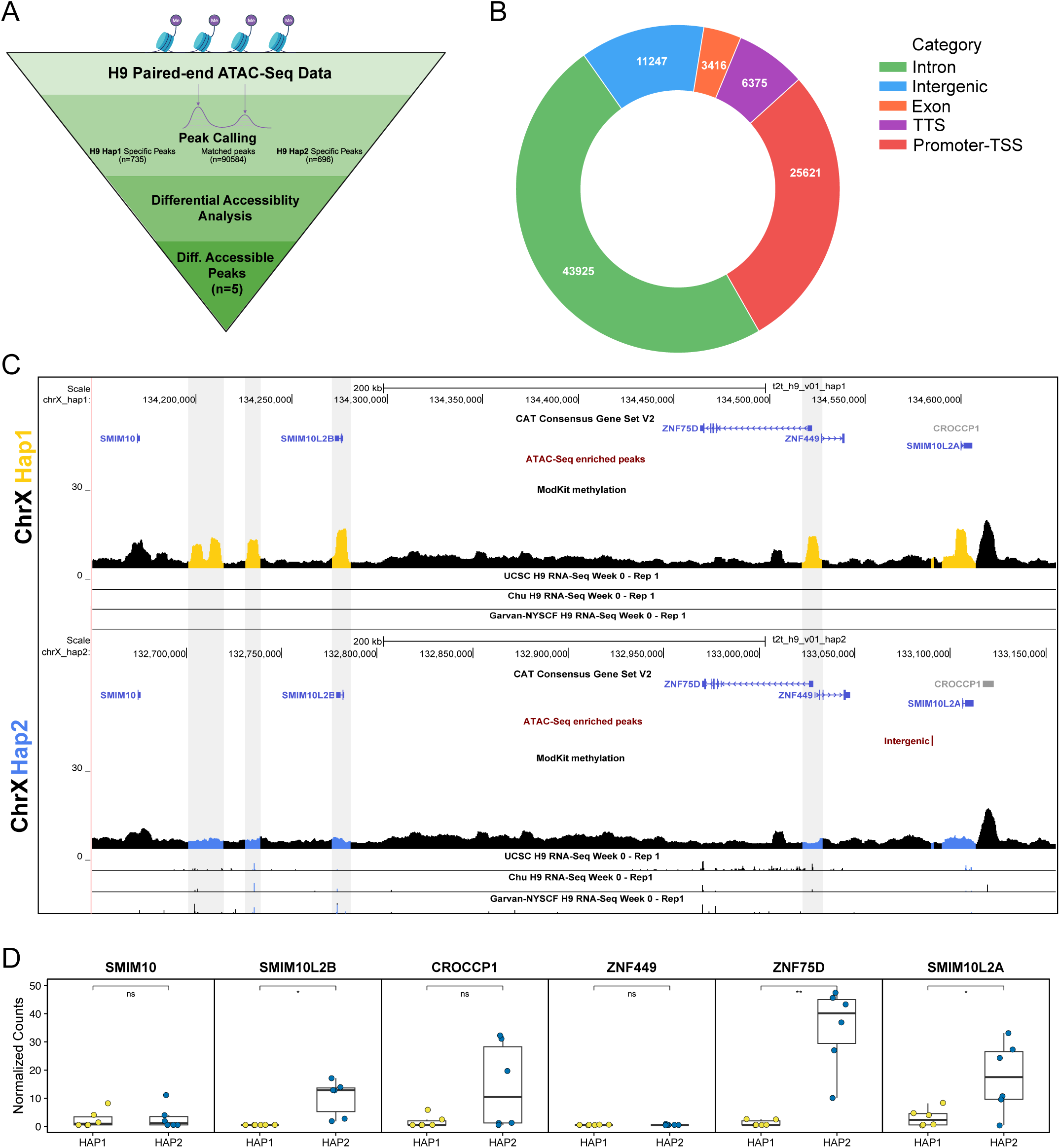
Haplotype-specific chromatin accessibility and transcriptional correlation in H9 cells. (**A**) Schematic workflow for H9 paired-end ATAC-Seq processing, showing the identification of haplotype-specific and matched peaks leading to five differentially accessible regions. (**B**) Donut chart illustrating the genomic distribution of the shared ATAC-Seq peaks between H9 haplotypes, with the majority located in promoter-TSS and intronic regions. (**C**) Genome browser tracks of the chromosome X locus showing haplotype-specific accessibility. Yellow (hap1) and blue (hap2) peaks highlight differential methylation enrichment and chromatin accessibility near genes including *ZNF75D*, *ZNF449*, and *SMIM10L2A*. (**D**) Box plots of normalized RNA-Seq counts for candidate genes, demonstrating higher expression levels on haplotype 2, correlating with the observed chromatin accessibility enrichment in panel **C**.

Differential chromatin accessibility analysis indicated that only 5 regions exhibited peaks in both haplotypes, but with statistically significant differences in enrichment between the two haplotypes, with |log2FC| > 1 (**Figure S25A**, **Table S23**), of which 3 out of 5 peaks show higher ATAC-seq enrichment in haplotype 1. The strongest signal is located in the promoter region of the noncoding RNA ENSG00000290832 gene on chromosome 7, with a >8-fold increase in accessibility, compared to the peak on the same region on haplotype 2. This result may indicate promoter silencing of the RNA gene in a haplotype-specific manner. A moderate enrichment of signal in haplotype 1 compared to haplotype 2 is located in the promoter region of the TPPP gene, which plays a key role in microtubule organization. Recent studies have indicated that the microtubule network is crucial for guiding cell division orientation in pluripotent stem cells, thereby maintaining proper self-renewal or differentiation potential.^68^ The third differential chromatin accessible region higher in haplotype 1 falls within the exonic region of the NT5CP1 pseudogene on chromosome 14. This exonic ATAC peak, with moderate effect size (fold change ∼ 2.4), may represent a cryptic cis-regulatory region for other genes in the region. For haplotype 2, there is one enriched peak within the intronic region of the GTF2IRD2B gene on chromosome 7, with a strong effect size (fold change >4). This gene encodes for a protein that is involved in transcription regulation. It is lowly expressed in H9 hESCs, but increases upon differentiation in the two differentiation time course data sets we examined. However, the level of expression and low density of SNVs within the transcripts precluded assessing haplotype-specific expression in this case, highlighting a limitation of short-read sequencing data. It is also possible that this intronic-enriched peak in haplotype 2 is part of a distal enhancer for other genes in the region.

Interestingly, haplotype 2 shows a differentially accessible chromatin region on chromosome X near the promoter region of the ZNF75D gene (**Figure 6C**) and 9 Kbps downstream of ENSG00000229015, a novel pseudogene ortholog to a mouse gene involved in testis differentiation (**Table S23**) reported to be expressed in sperm and other tissues but never described in hESC. Three different publicly available RNA-seq datasets from H9 differentiation to cortical neurons,^52^ definitive endoderm^53^ as well as an unpublished dataset from ESC, midbrain progenitors, and astrocytes (this study), were used to assess differential gene expression, focusing only on reads that map specifically in one of the two haplotypes (see **Methods**). We found strong evidence of haplotype-specific RNA expression for ZNF75D, as well as association with multiple differential peaks of DNA methylation. Because of higher methylation consistently associated with haplotype 1 this may point to evidence of X inactivation as a possible reason for differential accessibility and gene expression. Accordingly, this differential ATAC-Seq peak may contribute to an alteration of the chromatin landscape favoring gene expression in haplotype 2 (**Figure 6D**). ZNF75D encodes a KRAB zinc finger protein, likely involved in recruiting repressive chromatin to target sites. It also appears to bind to HERV-H elements,^69^ which play a characterized role in maintaining pluripotency.^70^

Allele-specific chromatin accessibility analysis in H9 cells undergoing early neural differentiation identified 735 haplotype 1–specific and 696 haplotype 2–specific ATAC-Seq peaks (**Figure S25B-C**). Gene ontology enrichment analysis of these haplotype-specific accessible regions (p-adj < 0.1) revealed distinct functional biases between haplotypes. Haplotype 1-specific peaks were slightly enriched in genes involved in the regulation of cell morphogenesis (GO:0022604, fold enrichment=4.00, p-adj=0.07), such as the DBN1 gene encoding the drebrin protein, thought to play a role in neuronal growth (**Table S24**). In contrast, the haplotype 2-specific peaks showed a strong enrichment in genes involved in the regulation of small GTPase-mediated signal transduction (GO:0051056, fold enrichment=4.69, p-adj=0.0017) (**Table S25**). GTPase-mediated signal transduction in hESC is an essential part of the cellular machinery that regulates self-renewal, differentiation, and cytoskeletal dynamics.^71^ In addition, haplotype 2 peaks were enriched for neurotransmission processes, such as positive regulation of synaptic vesicle fusion to presynaptic active zone membrane (GO:0031632, fold enrichment=26.19, p-adj=0.0068) and calcium-dependent activation of synaptic vesicle fusion (GO:0099502, fold enrichment=26.19, p-adj=0.0068). These terms were driven by presynaptic machinery genes such as DOC2B, SYT1, ERC2 and RPH3AL, encoding proteins involved in vesicle trafficking and neurotransmitter release. This suggests that allele-specific expression may modulate the H9 gene expression landscape both in pluripotency and upon differentiation.

## DISCUSSION

The last decade has seen a revolution in long-read sequencing technologies and genome assembly methods,^72^ enabling the generation of high-quality near-T2T diploid genome assemblies for any individual, population, and cell line of interest. To date, most of these genomes have been generated from lymphoblastoid cell lines generated by Epstein Barr virus transformation of donor peripheral blood mononucleocyte cells (PBMCs), and the focus of these assemblies has been on representing human diversity. The genetic variation revealed by these efforts highlighted the advantages of using a reference genome specific for the experimental model system from which functional genomic and multi-omic datasets are being generated. The recent near-T2T genome of the RPE1 cell line further showcased how a matched reference genome can provide important context for understanding the phenotypic features of the experimental model.^8,7^ Here, we present H9v1.0, a fully phased, T2T diploid genome assembly of the H9 hESC line. By integrating long-read sequencing by complementary technologies, Hi-C phasing, and orthogonal cytogenetic validation, we establish a structurally complete and functionally annotated genomic framework for the most widely used human pluripotent stem cell line for modeling human development and disease.

### hESC genomic architecture includes extended satellite repeats and freshly-added telomeres

H9v1.0 achieves chromosome-scale phasing with minimal gaps and high consensus accuracy across both haplotypes. Compared to other T2T assemblies, H9 appears to have longer telomeres that are enriched for canonical repeats. These features are supported by long-read coverage and FISH validation, and are in line with the detection of TERT expression in multiple RNA-Seq data sets from H9 cultured exclusively under pluripotency conditions but shut off upon differentiation. We also find extended centromeric arrays across multiple chromosomes compared to those found in previous near-T2T genome assemblies. Because centromere size and higher-order repeat organization can influence kinetochore assembly and chromosome segregation, it will be important to determine whether extended arrays represent a characteristic of embryonic-stage genomes or reflect selection during pluripotent stem cell culture. Together, these findings position H9 as a developmental model in which repetitive genome compartments remain in a structurally “youthful” state, and support a hypothesis that pluripotency is accompanied by distinctive repetitive DNA architecture, potentially linked to genome stability, proliferative capacity, and/or epigenetic plasticity.

### A genetically stable reference for clinical studies

H9 is one of the most extensively used hESC lines and is the only such line approved for clinical and experimental applications in Europe. The lack of structural variants previously associated with prolonged pluripotent stem cell culture^59,60^ or neurodevelopmental disorders^60^ reinforces its genomic integrity. While we identify interhaplotype structural variation in four of these regions, these appear to represent inherited polymorphisms rather than culture-acquired lesions. Three of the detected inversion-containing haplotypes have been previously identified.^61,673,763^ Importantly, no major pathogenic CNVs were detected. These results underscore that H9 retains a stable genomic configuration consistent with its use as the gold standard model system for developmental and translational studies.

### Ancestry analysis provides a population-genetic context for H9

Our comparisons with recent HPRC genome assemblies and SNP-based ancestry analyses place H9 near European and West Asian (Levantine) samples. Additional locus-specific analyses using ancestry inference along the genome find both European-like and West Asian-like segments. A 2010 study using short reads and a limited SNP chip array had pointed to the sample’s European genetic ancestry,^6^ consistent with the geographical source of the sample for cell line generation being a clinic in Israel.^30^ Our in-depth analyses show additional representation of segments from specific West Asian populations. We note that genomes of individuals of European Jewish heritage have often been reported to have both European-like and West Asian-like (Middle Eastern) ancestral components, although we cannot exclude another possible source for such genetic ancestries in H9. Embedding a widely used stem cell line within a population-genetic framework provides important context for haplotype-specific variation, copy number polymorphisms, and regulatory diversity. As our knowledge of the effect of extant human variation on genome regulation and function expands, such contextualization will become increasingly important for the interpretation of experimental and clinical outcomes.

### Benefits of using the correct reference

The most immediate translational impact of this resource lies in improved mapping of functional genomic data. Aligning identical RNA-Seq datasets to H9 versus GRCh38 results in hundreds of differentially mapped genes, including genes entirely absent from GRCh38 alignments. This demonstrates that reference choice alone can alter biological conclusions. We found that haplotype-aware analysis can reveal allele-specific differences in gene expression, promoter and UTR structure, miRNA and tRNA cluster organization, and variation in tandem repeat copy number and sequence, including those in centromeric and telomeric regions. A limitation is that most of the existing RNA-Seq data is short read, which poses challenges for identifying allele-specific expression above the threshold of heterozygosity, unless the gene has multiple variants within the transcript and is robustly expressed. However, with this isogenomic resource, future long-read RNA-Seq data sets could enable identification of haplotype-specific gene expression across the transcriptome.

To help experimental biologists take advantage of this new resource, we have built a UCSC Genome Browser Track Hub with browsers for each haplotype, including tracks allowing comparisons between the H9 haplotypes as well as the GRCh38 and CHM13 references, tracks for the gene and genomic feature annotations described and generated in this study, and additional coverage tracks for the published datasets we analyzed. As an example, integration of phased RNA-Seq, ATAC-Seq, and methylation profiling data and visualization in the browsers (**Figure 6C**) enabled us to identify regulatory asymmetries between the two haplotypes that are obscured when using a haploid and non-isogenomic assembly.

### A foundational resource for stem cell genomics

For more than two decades, H9 has served as a benchmark model for studies of human pluripotency, early development, and disease. As such, a wealth of functional genomic data has been generated using H9 for both undifferentiated and differentiated cells in a wide variety of lineages. H9v1.0 provides a matched, high-resolution genomic resource that can be used to extract a more precise picture of previously generated data and will inform new functional genomic studies to advance our knowledge of embryonic development, pluripotency, differentiation, and associated processes. As the field moves toward precision genome engineering^8^ and allele-aware regulatory modeling, matched diploid references for experimental model systems will be indispensable. H9v1.0 establishes such a framework for a foundational human embryonic stem cell line and complements ongoing T2T efforts that will add additional cell line models for a paradigm shift in the use of reference genome assemblies in scientific research for decades to come.

## ACKNOWLEDGEMENTS

We thank the High Performance Computing (HPC) Core Terastat2 and Terastat3 from Umberto Ferraro Petrillo of the Statistics Department, University of Rome Sapienza. We acknowledge all the members of the Giunta lab, especially Valentina Liguori for administrative support, as well as Luca Corda and Alessio Colantoni for helpful insights and discussion, and Isabel Kurth and Mauricio Orantes Bonilla for critical reading and help with editing. We acknowledge Regine S. Tipon for hESC culture and midbrain progenitor differentiation at NSYCF. Bonhwang Koo for helping submit the genome and data to NCBI. We acknowledge Dr. Ambra Tarquini and the Stem Cells and Organoids Facility, Dept. of Biology and Biotechnologies ‘Charles Darwin’, Sapienza University of Rome, for support. We thank Kristof Tigyi from the Salama Lab for preparing H9 samples for the various sequencing assays, the UCSCs Institute for the Biology of Stem Cells Tissue Culture Facility, and Max Haeussler and the UCSC Genome Browser Group for enabling our data analysis and for providing a platform to create a stable, publicly accessible resource for the H9 research community.

The study was performed in accordance with institutional guidelines and was approved by the Ethics Committee CET Lazio Area 1 (Ethical Approval Verbale 3_2024, 08.01.2024; Prot. 0662/2025, Ref. 8076, 17.07.2025, AIRC Project) and by the Ethics Committee of Sapienza University of Rome (Prot. 513/2023, 07.06.2023, CentroFun ERC StG project), where applicable.

## AUTHOR CONTRIBUTIONS

S.G. conceived the project; S.G., S.R.S., M.T.U., I.P., and M.D. designed the study and wrote the manuscript; S.R.S. and G.F.C. provided samples for the study; M.J., S.G., E.D.J, G.F., J.B., N.J., and F.A. provided the sequencing data; I.P., M.T.U., Y.C. and M.D. performed the main computational analyses; M.D. and I.P. generated the UCSC Genome Browser Track Hub and wrangled the data tracks; Y.C. and G.F. assembled the genome; Y.C., M.T.U., and I.P. performed assembly QC; J.A.M. performed the annotation and analyses of telomeres and three-dimensional genome features; M.T.U. performed centromeres annotation and analysis; P.H., M.D., A.T., P.C., and T.M.L. performed gene annotations; M.D., I.P., and S.R.S. conducted the segmental duplication and structural variation analysis; A.T. conducted the miRNA analysis; E.D.T. conducted the telomere FISH assay; S.G., M.T.U, G.F.C., G.F., M.G., and A.G.I. designed and performed the ancestry analyses; G.F.C. designed and supervised experiments for the new RNA-Seq data reported; K.D. differentiated astrocytes, and N.S. conducted ESC and midbrain experiments. I.P., H.D., S.G., S.R.S., M.D., G.F.C., J.E.P., and K.K.S. contributed to the computational functional genomics data analyses. I.P. extensively contributed to figure design with help from E.D.T., and M.T.U. All authors have reviewed and approved the manuscript.

## Declaration of generative AI and AI-assisted technologies in the manuscript preparation process

During the editing, some authors used AI services (i.e. Claude, Gemini, etc.) to check for spelling and grammar, and for suggestions on condensing specific sentences. After this, the authors reviewed the text in its entirety and thus take full responsibility for the content of the published article.

## DATA AVAILABILITY

A UCSC Browser hub with data associated with this project is available at https://public.gi.ucsc.edu/hausslerlab/t2t-h9-hub/. Annotations and datasets discussed in this manuscript are available as tracks in the browsers for each H9 haplotype and CHM13.

GenBank accessions of assembly: GCA_054883195.1 (haplotype 1), GCA_054883265.1 (haplotype 2)

SRA accessions/bioproject of reads used to construct the assembly: BioProject PRJNA1431686, SRA BioProject SRP680790

Locations and identifiers for other data:

- GenomeArk: https://genomeark.s3.amazonaws.com/index.html?prefix=species/Homo_sapiens/H9_T2T/genomic_data/

## CODE AVAILABILITY

Custom scripts used in this study and processed data are available at the GitHub repository https://github.com/GiuntaLab/H9.

## METHODS

### Sample collection and sequencing

Human embryonic stem cells (hESCs) H9 (Wisconsin International Stem Cell (WISC) Bank, WiCell Research Institute, WA09 cells) were cultured according to WiCell protocols^74^ on MEFs for three passages and banked at passage 27 (P27). One of these vials was then thawed onto two wells of a MEF containing six-well dish in StemFlex media (ThermoFisher). When 60-80% confluent, cells were passaged using 0.5 mM EDTA onto vitronectin (Thermofisher) coated 6 cm plates according to the StemFlex culturing recommendations. The culture was expanded by passaging every 4-5 days at 1:6-1:10 and banks were made at P31 and P34 yielding 5 vials/10 cm dish. A P34 vial was thawed, onto a 6 cm dish, expanded twice and sent for array-based karyotype analysis using the ThermoFisher KaryoStat assay and confirmed to have no significant chromosomal aberrations. To prepare cell pellets for sequencing assays, P34 vials from our bank were thawed onto a 6 cm VTN coated plate. Cells were expanded into 2, 10cm plates 4-5 days after thaw, keeping 1x 6cm as a passage plate, which was expanded to 3, 10 cm plates 4-5 days later. Plates were grown to 70-80% confluence to prevent overgrowth, differentiation, and cell deterioration. Counting of cells for pellets was done by dissociating cells with Stempro Accutase for 6 min at room temp. Cells were collected and pelleted at 300xg for 5 min, and resuspended with 1mL PBS with a P1000 initially, and then cell solution volume was brought up to 10mLs with PBS. Cell counts were done using trypan blue in a Countess 4 cell counter (ThermoFisher). Desired cell numbers (5, 10 or 20 million cells/tube) plus 10% for error were aliquoted into 15mL conical tubes for final pelleting at 300xG for 5 min. After aspirating off the PBS, samples were flash frozen with liquid nitrogen, stored at -80C and shipped to the sequencing centers on dry ice.

Approximately two million flash-frozen cells were processed for high molecular weight (HMW) DNA isolation using the Nanobind PanDNA Kit (Pacific Biosciences, PN: 103-260-000). The resulting DNA was quantified via the Qubit 4 Fluorometer (Thermo Fisher Scientific, Waltham, MA, USA) and assessed for quality with the Agilent FEMTO Pulse system (Agilent Technologies, Santa Clara, CA, USA).

For sequencing, the HMW DNA was mechanically sheared to a target size of 23 kb using the Megaruptor 3 (Diagenode, Denville, NJ, USA) and purified with Ampure PB beads. Using 5 μg of sheared and cleaned DNA as input, PacBio HiFi sequencing libraries were then constructed using the SMRTbell prep kit 3.0 (Pacific Biosciences, PN 102-182-700). Fragments shorter than 10 Kbps were eliminated via size selection on the Pippin HT instrument (Sage Science, Beverly, MA, USA). Finally, the size-selected libraries underwent annealing, binding, and cleanup with the PacBio Binding kit 3.2 (Pacific Biosciences, PN: 102-333-300) before being loaded onto two PacBio SMRT Cells and sequenced on the PacBio Revio Instrument, yielding a total of 227 Gbps of data.

For Nanopore sequencing, DNA extractions were performed using 6 million cells and the NEB Monarch HMW DNA extraction kit for tissue (New England Biolabs T3060). Nanopore libraries were prepared using the ONT ultralong DNA sequencing kit (SQK-ULK114). Sequencing was performed using R10.4.1 PromethION flow cells for 72 h. Flow cells were washed using the flow cell wash kit (EXP-WSH004) every 24 h. Fresh libraries were loaded after each wash.

Separately, the Arima Genomics v2 kit (Arima Genomics, PN: A410110) was utilized to generate 264 Gbps of Hi-C data via the Illumina Novaseq 6000 platform; approximately one million flash-frozen cells were crosslinked for HiC using the Arima High Coverage HiC Kit (PN: A410110) following the Mammalian Cell Lines with low input procedure. DNA was then ultrasonically fragmented to approximately 550 bps using a Covaris S220 ultrasonicator (Covaris, Woburn, MA, USA) and purified with Ampure XP Beads (Beckman Coulter, PN: A63880). Purified DNA then underwent library preparation using the Arima Library Prep Module following the manufacturer’s procedure. Library quantity and quality were assessed using the Qubit 4 Fluorometer (Thermo Fisher Scientific, Waltham, MA, USA) and the Agilent Fragment Analyzer system (Agilent Technologies, Santa Clara, CA, USA). Sequencing was done on the Illumina Novaseq 6000 platform yielding 264 Gbps.

### Genome assembly and curation

Two assembly strategies were attempted in Verkko v2.2.1. The first assembly (asm1) used HiFi reads to build the graph, ONT reads for graph resolution, and Hi-C data for haplotype phasing. This resulted in 27 complete T2T chromosomes with no gaps. The second assembly strategy (asm2) used both HiFi reads and hifiasm-corrected ONT reads to generate the graph, raw ONT reads for graph resolution, and Hi-C data for haplotype phasing. The ONT read correction was done in HiFiasm v0.25.0-r726. This asm2 generated 31 complete gapless T2T chromosomes. Since neither approach achieved T2T level for all chromosomes, we selected the best chromosomes from each to construct a combined final assembly (v1.0). Specifically, we included all 27 T2T chromosomes from asm1, along with 9 additional T2T chromosomes from asm2. The remaining 10 chromosomes that did not reach T2T standards, were selected from the more complete assembly – favoring fewer gaps and presence of telomeres – and manually curated in BandageNG v2024.07-dev.

To facilitate manual curation, we assigned the chromosome names by aligning each assembly to CHM13v2.0 (NCBI accession GCF_009914755.1). Graphs were annotated for telomeres identified by Teloscope v0.1.3, and rDNA using the script https://github.com/marbl/training/blob/main/part2-assemble/docker/marbl_utils/verkko_helpers/remove_nodes_add_telomere.py with the human rDNA sequence (NCBI accession KY962518.1) as a bait. Graphs were colored using phasing information from Hi-C data. Gfalign tool was used to evaluate all possible paths for tangled gaps. For paths that are equally well-supported by ONT reads, we randomly selected one path to represent the region (chromosome 5 hap1 and chromosome 16 hap1, **Table S1**). Disconnected scaffolds caused by rDNA tangles in three chromosomes were joined based on Hi-C evidence, with rDNA gaps inserted at the junctions. The rDNA tangles on eight chromosomes could not be resolved and were retained as gaps in the final assembly. The details of chromosome selection from two assembly strategies and the manual curation are provided in **Table S1**. The consensus sequences of the curated paths were computed in Verkko, resulting in the final combined assembly.

### Assembly QCs

Assembly metrics and quality controls (QCs) have been performed in line with standard approaches for evaluating genome contiguity, completeness and correctness, as well as with other recently published T2T genomes.^6,75,7^ Specifically, contiguity was measured as an Nx plot by looking at individual contigs’ length against the entirety of the genome for both haplotypes; assembly basic statistics were computed using Gfastats v1.3.11.^76^ Gene completeness was determined by compleasm^18^ v0.2.7 using *mammalia_odb12* as the gene dataset, whereas overall assembly completeness and QV were assessed with Merqury^19^ v1.3 using *21*-mers. Detection of anomalies in read coverage along the assembly was performed using HMM-Flagger^23^ v0.4.0, classifying the assembly into the following categories: erroneous, falsely duplicated, haploid (structurally correct), and collapsed. HiFi reads were mapped to each haplotype using the minimap2^37^ v2.30 with the *map-hifi* option and genome-wide coverage plots were generated using NucFreq^77^ v0.1. We also mapped HiFi and ONT reads to the diploid genome using minimap2^37^ v2.30 and assessed the coverage in NucFlag^46^ v1.0.0a2 with a window size of 50 Kbps after removing unmapped, secondary, and supplementary reads using SAMtools^78^ v1.22 (-F 2308). Local regions were assessed at a default window size of 5 Kbps in NucFlag. The other parameters were kept as default in NucFlag.

### Pangenome-based PCA and Ancestry inference analysis

The pangenome PCA was computed based on a PGGB^79^ v0.7.4 graph built from the initial year one HPRC samples^24^ plus the two human references (GRCh38 and CHM13), with the inclusion of the H9 genome hereby assembled; this dataset totals 47 genomes, adding up to 92 haplotypes. The exact options and commands are those in Liao *et al.* 2023, in concordance with high-quality, contig-level human genomes. To determine contig placement in graph space, each haplotype was aligned against CHM13 with minimap2^37^ v2.30 using options for intraspecies genome-to-genome alignment, and subsequently, pangenomes were independently built in parallel for each chromosome; this strategy was envisaged due to the high computational demand of the construction step. Later, these chromosome pangenomes with all individuals were parsed back together employing ODGI^80^ squeeze v0.9.2 to generate the final data structure for the analysis. To output a similarity matrix from pangenome alignments, ODGI similarity (v0.9.2) was employed for processing the graph into a tractable score-based tabular file opportunely rendered for PCA plotting.

Ancestry inference was performed with GenoTools^26^ v1.3.6 which uses predetermined ancestry components and labels to assign to the queried sample(s) the most parsimonious population category according to a panel — in this case primarily built upon the 1KG+HGDP integrated datasets and provided as an available resource for the analysis. Three human T2T genome assemblies with known ancestry (HG002,^6^ YAO^5^, and RPE1^7^) were processed along with H9 through GenoTools to ascertain the inferential power and correctness of this approach. VCF files were generated from high coverage HiFi reads for all samples aligned to the HPRC minigraph-CACTUS^24,81^ pangenome graph. Alignments were run in personalized mode with VG^82,83^ v1.68.0 to capture more accurately individual variation, and then projected to GRCh38 coordinate space since this is the reference for the GenoTools panel. Variants were called with DeepVariant^84^ v1.9.0 and a joint VCF (jVCF) was produced with GLnexus^85^ v1.4.1. The jVCF was postprocessed with bcftools^86^ v1.22 to remove missing sites and restrict the analysis to only SNVs. Repetitive and low mappability regions, centromeres as well as segmental duplications from the UCSC browser coordinate files for GRCh38 were filtered out from the final file. PLINK2.0^33^ was employed to output the PGEN and BIM files needed for GenoTools, which was then executed with the commands --ancestry and --pca required to infer a sample’s ancestry and plot it in PC space based on the provided panel. In both cases, the PCA analysis was computed only on the 22 autosomes, considering phased positions and only reliable alignments and calls.

Local ancestry inference was performed with PCLAI^31^ using default settings. The employed PCLAI model was trained on an autosomal subset of 26062361 biallelic SNPs from a 1KG and HGDP merge with 2356 training samples. The PCLAI principal component embedding space was spanned using the same reference individuals, pruning the variants to 300000 max-heterozygosity SNPs using PLINK2.0. Forward-in-time simulation has been used to augment the training dataset and introduce coordinate-annotated admixed samples. For inference, we intersected H9 sites with the reference panel; phasing and imputation of missing sites was done with BEAGLE 5.5.^87^ We computed the three population statistic F_3_ using snputils^34^ in the form of F_3_(O; H9, X) where the fixed outgroup O is the Yoruba from Ibadan (YRI) and X is a reference from the Mediterranean, Levantine, and North African subset from the 1KG and HGDP combined panel containing samples from the following subpopulations: Basque in France, Bedouin in Israel, Bergamo Italian in Italy, Druze in Israel, French in France, Mozabite in Algeria, Palestinian in Israel, Sardinian in Italy, Spanish in Spain, and Tuscan in Italy. Supplemental PCA-based ancestry analysis was computed on the same subset as F_3_.

### SNVs density, HDRs, and centromeres

To preliminarily quantify differences at the haplotypic level, we determined differential peaks in single nucleotide variants between haplotype 1 and haplotype 2. This was done by aligning haplotype 2 against haplotype 1 (as reference) with minimap2^37^ v2.30 with the option -x asm5 --eqx and, otherwise, default parameters. Subsequently, the SyRI^54^ v1.6.3 output containing synteny and structural rearrangement types and counts was post-processed to extract chromosome-wide SNVs along with highly divergent regions (HDRs). Finally, centromeres were added after their identification with dna-brnn^45^ v0.1 run in default mode and restricting the BED output to only alpha satellites, and annotation by Hum-AS-HMMER for AnVIL (https://github.com/fedorrik/HumAS-HMMER_for_AnVIL).

Similarly, the SyRI analysis conducted to ascertain structural rearrangements in both haplotypes against the same T2T reference genome in CHM13^1^ was computed post genome-to-genome alignment with minimap2 v2.30 and the options above, followed by a SyRI run in default mode. The plotting output has been done with the in-built Plotsr feature of SyRI.

### Centeny maps and centromere annotations using GCP pipeline models

The whole-genome and centromere-specific characterizations were performed using our previously described Genomic Centromere Profiling (GCP) pipeline, using CENP-B boxes based annotation of the motif position and orientation (blue = forward; red = reverse complement sequence found within the same DNA strand) to define the *centeny* maps. Additional annotations of centromere structure were added using GCP pipeline Models 1 and Models 2 showing the inter-motif distances within centromeres (Model 1) and the organization of the motif within the alpha-satellite monomers (Model 2).^39^

### Gene annotation

Genome annotation was performed using the Comparative Annotation Toolkit (CAT) (improving upon ^35^). First, whole-genome alignments between the two H9 haplotypes, the two RPE1 haplotypes, the two HG002 Q100 haplotypes, GRCh38, and CHM13 genomes were generated using Cactus ^81^. CAT then used the whole-genome alignments to project the GENCODE v44 annotation set from GRCh38 to all the other genome assemblies. CAT was run with transMap^88^, transMap-pairwise, AUGUSTUS,^89^ Liftoff,^90^ and AUGUSTUS-PB modes. transMap lifts over gene annotations from the reference onto all the genomes in the cactus alignment. transMap-pairwise works the same way but with pairwise minimap2 alignments between the reference and each of the target genomes. Liftoff lifts over gene annotations from a reference onto minimap2 alignments between individual reference transcripts and the target genome. CAT was given PacBio IsoSeq^91^ data to provide extrinsic hints for *ab initio* prediction of coding isoforms. CAT then combined these *ab initio* prediction sets with the various gene projection sets to produce the final gene sets used in this project.

### MicroRNA analysis

For microRNAs, 1,968 human miRNAs listed in miRBase^92^ 22.1 were grouped into 1,437 families based on MirGeneDB^93^ annotation or simply by their microRNA identifier. All microRNAs were searched using BLAST^94^ v2.17.0 against the primary assembly of GRCh38, CHM13 and both haplotypes of H9 with 100% and 95% sequence identity cutoff, respectively. Overlapping results of orthologous miRNAs were removed, and only the best match per locus was kept.

### tRNA gene annotation

tRNA genes were predicted using tRNAscan-SE^43^ 2.0 with -Hy --mt mammal --detail options. Predictions that were found to have mitochondrial origin (within nuclear-mitochondrial DNA segments) were filtered. tRNA genes were then classified into high scoring predictions and potential repetitive elements using EukHighConfidenceFilter in tRNAscan-SE with default parameters. To preserve the tRNA gene names across different human genomes, the gene sets in H9 were mapped against the tRNA genes in CHM13 based on the whole genome sequence alignments and liftover^95^ between assemblies. Genes found to have identical corresponding loci in CHM13 and/or GRCh38 were assigned identical gene names and HGNC identifiers. Genes only found in H9 were assigned gene names following the naming convention of GtRNAdb.^96^ tRNA gene classifications were adjusted accordingly to maintain consistency between the H9 haplotypes and CHM13.

### Chromosome conformation capture annotation

Adaptor sequences from Arima Hi-C paired-end reads were trimmed with cutadapt^97^ v5.1. A total of 874 million trimmed read pairs were aligned to the diploid, haplotype 1, and haplotype 2 assemblies using BWA-MEM^98^ v0.7.17-r1188 with the -SP5M flags. Alignment quality was evaluated with SAMtools^78^ stats v1.22 and rdeval.^99^ Hi-C contact maps and downstream 3D genome annotations were generated with the Open Chromosome Collective (Open2C)’s pipeline comprising pairtools,^100^ cooler,^101^ and cooltools^102^. Hi-C read pairs were extracted from name-sorted BAMs using pairtools parse using the default --walks-policy mask, followed by pairtools sort and pairtools dedup with --max-mismatch 0. After deduplication, 6.8, 244.9, and 245.0 million unique pairs (UU) remained for the diploid, haplotype 1, and haplotype 2 assemblies, corresponding to 94.4%, 58.6%, and 58.6% of valid pairs, respectively.

Contact matrices (.cool) were built with cooler cload pairs with 5 Kbps bins and normalized with cooler balance. Distance-dependent contact decay (P(s) curve) was computed with cooltools expected-cis. Multi-resolution maps (.mcool) were generated with cooler zoomify at 10, 20, 50, 100, and 200 Kbps. Gene density tracks were computed at each resolution using bedtools coverage and the gene annotation. A/B compartments were called from the first cis eigenvector computed with cooltools eigs-cis on the .mcool file. We used the coding gene density as a phasing track to orient the first eigenvector (E1) sign so that positive E1 corresponds to higher gene density. Compartmentalization strength was summarized with saddle plots generated by cooltools saddle. Topologically associated domain (TAD) boundaries were computed at 200 Kbps using cooltools insulation.

### Telomere annotation and validation

Telomere annotation was performed using Teloscope^103^ v0.1.3, developed in the Vertebrate Genome Lab. Teloscope was run with default parameters -w 1000 -s 500 -c TTAGGG -p TTAGGG -d 200 -l 500 and custom flags -x 1 -g -r -e -m -i to scan for canonical and noncanonical telomeric repeats, allowing up to one mismatch. Teloscope also generated genome-wide window metrics and a summary report with statistics per assembly. Scaffolds were assessed and classified based on telomere completeness (T2T, incomplete, miss-assembly, and discordant) and gap presence. In addition to the H9v1.0 assembly, we also annotated publicly available assemblies from NCBI and additional references of interest, including GRCh38, CHM13,^1^ HG002,^6^ I002C,^4^ YAO,^5^ and RPE1^7^. This enabled direct comparisons of telomere completeness, assembled telomere lengths, and canonical proportions. To validate our findings, we re-aligned our HiFi reads and ONT reads against the H9 haplotypes using minimap2^37^ v2.30 and flags -ax HiFi and -ax lr:hq --eqx, respectively. We filtered unmapped reads, secondary and supplementary alignments using SAMtools^78^ v1.22 with flag -F 2308 and assessed the coverage for the 25 Kbps most distal region of each chromosome end using NucFlag v1.0.0a2 with -x hifi and -x ont_r10 for HiFi and ONT reads, respectively.

### FISH analysis

H9 cells were maintained at 37°C in a 5% CO2 atmosphere. These cells were grown in StemFlex media (ThermoFisher) supplemented with penicillin–streptomycin 1x (Sigma-Aldrich) in Vitronectin (VTN-N; ThermoFisher) coated dishes and passaged every 4-5 days using 0.5 mM EDTA. RPE1 cells were maintained at 37°C in a 5% CO2 atmosphere with 21% oxygen and were grown in DMEM/F-12 media (Corning 10-092-CV) supplemented with 10% fetal bovine serum (FBS; Life Technologies A5256701) and 100 U/ml penicillin–streptomycin (P/S; Corning 30-002-CI). RPE1 hTERT cells were obtained from a parental RPE1 line infected with the pLVX-hTERT-IRES-Hygro lentiviral plasmid and selected for hygromycin resistance and grown in DMEM/F-12, 10% FBS, P/S.

FISH for telomeric DNA was performed as previously described.^104^ Briefly, H9 and RPE1 cells were incubated for 3-4 hours with 0.1 μg/ml colcemid. Cells were harvested and swollen in prewarmed 0.075 M KCl at 37°C for 30 min. They were then fixed in fresh 3:1 methanol/acetic acid and dropped onto glass slides. After aging overnight, the slides were washed in 1× PBS twice for 2 min each and treated with RNase, washed in 1× PBS and water followed by consecutive incubation with 70, 85, and 100% ethanol. The slides were allowed to air dry before applying a hybridizing solution (60% formamide, 1 mg/ml blocking reagent [Roche], 20 mM Tris-HCl, pH 7.2) containing selected PNA probes (PNABio). The spreads and the probe were denatured for 5 min and hybridized for 10 at 80°C on a heat block and then 1h at RT. The slides were washed three times 5 min each with wash solution (0.1M Tris-HCl, pH 7.2, 0.15M NaCl, 0.08% Tween20), with the second wash supplemented by DAPI (Sigma D-9542). Slides were mounted in antifade reagent (ProLong Glass; Life Technologies P36980) and imaged. The telomere probe F1004 TelC-Alexa488 (CCCTAA repeats) from PNABio has been used for the assay.

Image acquisition was done on a Thunder Leica widefield fluorescent microscope at a 100x magnification and with a 0.2 µm z-stack and processed using FIJI-ImageJ^105^ to obtain the maximum projection images visualized in **Figure 4C**; these images have been imported and processed in Photoshop to adjust brightness and contrast for visualization purposes (Adobe Systems). In both H9 and RPE1 we applied the same acquisition parameters. Following acquisition, quantification of FISH images at telomeres was performed using the Imaris software (Bitplane) surface fitting function and extracting data on each telomere volume and sphericity.

### Impact of isogenomic diploid phased reference on transcriptome analysis

We analyzed bulk single-end RNA-Seq data from three biological replicates of H9 cells (BioProject: PRJNA735229; SRA accession numbers: SRR14735598, SRR14735599, and SRR14735603), generated by Zijlmans *et al*.^106^ Adapter trimming and quality assessment were performed using Trim Galore^107^ v0.6.10, with Cutadapt^97^ v2.868, and FastQC^108^ v0.11.9. Trimmed reads were aligned to H9 haplotype 1 and H9 haplotype 2 with the corresponding CAT gene annotation files generated in this study, and to GRCh38 with the GENCODE v49 gene annotation using STAR^109^ v2.7.11b. Aligned reads were quantified using featureCounts^110^ v2.1.1, restricting the analysis to genes present in both references. Subsequently, differential gene expression analysis was performed using edgeR^111^ v4.8.0. Our design consisted of running pairwise comparisons between the biological replicates aligned to different reference genomes (H9 Hap1 vs. GRCh38 and H9 Hap2 vs. GRCh38) to assess the effect of using an isogenomic reference on transcriptomics analysis. Genes with a false discovery rate (FDR) < 0.05 were considered differentially mapped.

Furthermore, we assessed the impact of using the new H9 diploid, phased reference assembly for uncovering haplotype-specific gene expression profiles by running a differential gene expression analysis between H9 haplotypes. Then, we investigated the putative role of the SNVs on allele-specific expression. To do so, we intersected the SNVs between H9 haplotypes, previously generated by the SyRI analysis, with the gene annotations of H9 haplotype 1, using bedtools^112^ intersect v2.27.1. Subsequently, we mapped each of the SNVs to a specific genomic annotation. As a single SNV can overlap multiple genomic features depending on the transcript, we applied the following priority rule during the assignment of the genomic feature: coding DNA sequence, UTR, non coding exon, intron, promoter, intergenic. A promoter was defined as the region spanning 2000 bps upstream of each gene.

### ATAC-Seq data analysis

Publicly available ATAC-Seq raw data, generated from H9 cells undergoing early neural differentiation (Bioproject: PRJNA1235757),^67^ were downloaded to perform chromatin accessibility analysis across H9 haplotypes. Specifically, we downloaded paired-end ATAC-Seq data from 10 samples, with the following accession numbers: SRR32687946, SRR32687947, SRR32687948, SRR32687949, SRR32687950, SRR32687951, SRR32687952, SRR32687953, SRR32687954, and SRR32687955.

Quality control and adapter trimming were performed with TrimGalore^107^ v0.6.10 using default parameters. Then, we used bowtie2^113^ v2.5.4 to separately align the reads against the H9 haplotypes with the following parameters: --end-to-end --sensitive --no-mixed --no-discordant --no-unal. Afterwards, we applied MACS3^114^ v3.0.3 to call statistically significant narrow peaks from mapped paired-end reads, with quality score > 20, using the following parameters: -f BAMPE -B -g 3.00e9 -q 0.01 --nomodel. We then proceeded to perform a differential chromatin accessibility analysis between H9 haplotypes. To do so, we mapped H9 haplotype 1 fasta genome to H9 haplotype 2 using minimap2^37^ v2.30, generating a PAF file. This latter was used to liftover the coordinates of narrow peaks called in haplotype 1 to haplotype 2. Subsequently, we intersected these lifted over peaks with those of the original haplotype 2 narrow peaks coordinates using the bedtools^112^ v2.27.1 intersect module, generating a set of matched peaks across H9 haplotypes. Aligned reads to each H9 haplotype were quantified with the bedtools^112^ v2.27.1 multicov module, restricting the analysis to the alignments within the matched peaks. The resulting counts were combined into a single matrix, which was used for the differential chromatin accessibility analysis with DESeq2^115^ v1.50.0. Chromatin regions, with a p-adj value < 0.05 and |log2FC| > 1, were considered differentially accessible between H9 haplotypes. We used the HOMER v5.1 annotatePeaks.pl algorithm to annotate the called narrow peaks to nearby genes and to associate them to genomic features (promoters, introns, exons, etc.), as defined in the H9 gene annotation files.^116^ Then, we conducted a gene ontology analysis, using the clusterProfiler^117^ v4.18.1 R package, in order to uncover biological processes enriched in differential H9 chromatin accessibility regions as well as those in a haplotype-specific way.

### RNA-Seq Expression Analysis

Three publicly available H9 RNA-Seq datasets were downloaded: a definitive endoderm differentiation study from the Morgridge Institute for Research (PRJNA305280), a Garvan-NYSCF study (PRJNA1433892) and a cortical brain organoid differentiation study from UCSC (PRJNA415990). Illumina short-read RNA-Seq reads were mapped to the genome assemblies using STAR^109^ without reference gene annotations. Mapping to each H9 haplotype assembly allowed for reads that mapped to no more than 20 locations, randomly keeping the best scoring mapping. For the allele-specific expression experiments, only reads that map uniquely to one location in the combined H9 diploid assembly are used. (see **Supplementary Information**)

The featureCounts^110^ was used to determine the number of aligned reads per gene. Differential expression for target genes between haplotypes was analyzed using DESeq2^115^ v1.50.1. Genes were considered differentially expressed if they exhibited a |log2FC| > 1 and p-adj < 0.05.

### Genomic alignments

Pairwise alignments between the CHM13 and H9 haplotypes were generated using minimap2 with the asm5 preset. The resulting alignments were chained, and a syntenic subset was selected using the UCSC Browser alignment tools.^118^ See **Supplementary Information** for details.

### Identification of segmental duplications

Segmental duplications were annotated using SEDEF (ECCB 2018). Before annotation, the H9 assemblies were soft-masked using WindowMasker^119^ v2.5.0, RepeatMasker v4.2.1 with DFAM^120^ 3.9, and Tandem Repeats Finder^121^ (TRF). SEDEF was then run on the masked assemblies using default parameters.

For CHM13 v2.0, segmental duplications were re-annotated using SEDEF with rDNA arrays hard-masked, as these sequences are not present in the H9 assembly. rDNA regions were identified using Centromeric and Pericentromeric Satellite Annotation (cenSat)^122^ data obtained from the UCSC Genome Browser. WindowMasker, repeat, and TRF annotations were likewise obtained from UCSC, and SEDEF was run using the same parameters as for H9.See **Supplementary Information** for details.

### Cross-assembly segmental duplication mapping

Segmental duplication annotations were mapped between assemblies using the transMap projection mapping algorithm, implemented with UCSC Browser utilities. Unlike liftOver, this approach produces base-level alignments, which are useful for diagnosing alignment issues. Mappings were generated between CHM13 and each H9 haplotype, yielding two sets of projected SD annotations alongside a direct SD annotation for each assembly, enabling three-way comparisons.See **Supplementary Information** for details.

### ONT methylation analysis

ONT reads were aligned to the H9 diploid genome assembly using minimap2 --preset=map-ont. BEDMethyl files were produced using ONT Modkit^123^ v0.5.0. --pileup command with default parameters. To enable visualization in the UCSC Genome Browser, BEDMethyl was converted to BigWig using bedGraphToBigWig.

### Cell line specific variant visualization

CNV location and structure between the two H9 haplotypes, CHM13, and hg38 was visualized with SVByEye^124^ v0.99.0. using the plotAVA function. The haplotype 1 and haplotype 2 outputs of the SyRI^54^ synteny plot were combined into the final visual using custom code. Whole genome annotation karyotype plots were generated using karyoploteR^125^ v1.36.0.

