## Supplementary Information File for "The diploid reference genome of a human embryonic stem cell line"

### Supplementary information and figures for: The diploid reference genome of a human embryonic stem cell line


SUPPLEMENTARY INFORMATION
- RNA-Seq Expression Analysis
- Midbrain Floor Plate Progenitor and Astrocyte RNA-Seq data set generation
- Telomere validation and NucFlag classification criteria

REFERENCES

SUPPLEMENTARY FIGURES


#### SUPPLEMENTARY INFORMATION

##### RNA-Seq Expression Analysis

STAR[1] 2.7.11b was used to map Illumina short-read RNA sequences to the genome assemblies for expression analysis.

The genome was indexed using the following command:

```
STAR --genomeSAindexNbases 12 \
     --runThreadN 32 \
     --runMode genomeGenerate \
     --genomeFastaFiles ${genome_fa} \
     --genomeDir ${genome_star}
```

Haploid mappings of short reads were performed with:

```
STAR --runThreadN 16 \
     --genomeDir ${genome_star} \
     --readFilesType ${read_files_type} \
     --readFilesIn ${read_files_in} \
     --outFileNamePrefix ${outpre}. \
     --outTmpDir ${tmp_dir} \
     --outSAMtype SAM \
     --outStd SAM \
     --outSAMunmapped Within \
     --outSAMattributes NH HI AS nM NM MD \
     --outFilterMultimapNmax 20 \
     --limitBAMsortRAM 60000000000 \
     --twopassMode Basic --outSAMmultNmax 1 \
```

```
        --outMultimapperOrder Random
```

Diploid mappings of short-reads were performed against the combined H9 assembly with:

```
    STAR --runThreadN 16 \
        --genomeDir ${genome_star} \
        --readFilesType ${read_files_type} \
        --readFilesIn ${read_files_in} \
        --outFileNamePrefix ${outpre}. \
        --outTmpDir ${tmp_dir} \
        --outSAMtype SAM \
        --outStd SAM \
        --outSAMunmapped Within \
        --outSAMattributes NH HI AS nM NM MD \
        --outFilterMultimapNmax 1 \
        --limitBAMsortRAM 60000000000 \
        --twopassMode Basic \
        --outSAMprimaryFlag=OneBestScore
```

##### Midbrain Floor Plate Progenitor and Astrocyte RNA-Seq data set generation

ESC were cultured on lamin521(biolamina) in StemFlex(Thermo) as previously described.[2,3] Midbrain differentiation was performed per[4,5] with the following changes: organoids were cultured day 0-33 with no dissociation, no DAPT was used, 1%Albumax was added throughout, and SHH protein was replaced with 1uM SAG.13 and 1uM Purmorphamine, as described.[6] Organoids were dissociated at day 33 with dispase (Worthington, following the manufacture's protocol[7]), and cryopreserved.[8] Midbrain astrocytes were differentiated from day33 midbrain floorplate progenitors generated as described above with minor adaptations to [9], and harvested at day 125. For 10x Chromium captures, ESC was captured from freshly dissociated colonies, while differentiated cryopreserved cells were thawed, depleted for dying cells with AnnexinV microbeads (Miltenyii, Dead cell removal kit), and debris was floated with a density cushion (4% BSA in L15, adjusted to pH7). Cells were hashtagged, pooled, and captured on 10x Chromium 3' Gene Expression, ~20K paired-end reads/cell following standard methods. Reads were demultiplexed using a combination of hashtagging (Biolegend) and isogenomic SNP data reference, quality control for poor quality cells and doublets was performed, and reads for all cell types were pooled for each cell type (>2000 cells each) for pseudobulk analysis used in this study.

##### Telomere validation and NucFlag classification criteria

We validated Teloscope terminal telomeres using independent HiFi and ONT read evidence summarized by NucFlag. Telomere BED intervals were intersected with technology-specific NucFlag BED tracks on the same contig using strict overlap. Raw NucFlag types were grouped into five status classes: PERFECT (correct), HARD_ERROR (misjoin, collapse, false_dup, deletion, insertion, mismatch, softclip), MISMATCH_SIGNAL (het_or_mismap, low_quality), REPEAT_CONTEXT (homopolymer, dinucleotide, simple_repeat, other_repeat), and GAPS (scaffold). Pairwise labels were assigned per overlapped segment with rescue-first precedence: ONT_rescued if ONT=PERFECT and HiFi=HARD_ERROR; HiFi_rescued if HiFi=PERFECT and ONT=HARD_ERROR; concordant if at least one platform was PERFECT and neither was HARD_ERROR; error only if both platforms were HARD_ERROR; otherwise repeat_context if either platform was REPEAT_CONTEXT; otherwise mismatch_signal. This criterion is conservative because error requires replicated HARD_ERROR in both technologies, while mismatch peaks and repeat-context calls are retained as ambiguity rather than misassembly.

Across 692,615 bps of Teloscope-annotated telomeric DNA within the NucFlag-evaluable windows, 628,307 bps (90.7%) were supported by at least one technology (concordant + ONT_rescued + HiFi_rescued), leaving 64,308 bps (9.3%) in non-supported categories. Replicated HARD_ERROR in both technologies was rare, at 1,431 bps (0.21% of telomeric DNA), and provided an upper bound on the high-confidence telomeric error signal. Non-supported bases were dominated by ambiguity classes, with repeat_context accounting for 44,411 bps (6.41%) and mismatch_signal for 18,466 bps (2.67%). ONT contributed broader PERFECT support (532,347 bp; 76.9%) but a higher mismatch-signal burden (113,687 bps, 16.4%), whereas HiFi contributed fewer PERFECT-labeled telomeric bases (335,484 bps, 48.4%) but a substantially lower mismatch-signal burden (25,206 bps, 3.6%). A major discordance mode was the ONT rescue of HiFi repeat-context at distal tips. HiFi simple_repeat totaled 286,879 bp and 243,770 bps (85.0%) paired with ONT correct, indicating longer ONT reads more often bridging from unique subtelomere into the terminal repeat tract and stabilizing mapping at chromosome ends.

Chromosome morphology strongly modulated HiFi correct% when repeats were included (Kruskal Wallis H=26.201, p=2.04e-6), driven by markedly lower HiFi correct% at acrocentric ends (24.8±24.9%, n=19) compared with metacentric (84.5±25.6%, n=20) and submetacentric (72.1±32.3%, n=52). In contrast, ONT correct% showed no morphology dependence when all bases were included (H=1.612, p=0.447), but after excluding repeat_context bases, ONT correct% became significantly higher on acrocentrics (94.6±21.5%) than on metacentrics (68.3±35.8%) and submetacentrics (69.1±35.2%). Error% did not differ by morphology after correction, indicating that the primary morphology signal reflects correct-classification capacity in repeat-rich ends rather than enrichment of replicated hard errors. This pattern is expected from the biology of acrocentric short arms, which contain extensive shared homology and rDNA-adjacent repeat structure that increases mapping ambiguity, and from long-read technology trade-offs where ONT ultra-long reads improve spanning and anchoring across complex repeats, while HiFi provides higher base accuracy and lower mismatch-driven ambiguity.

### SUPPLEMENTARY FIGURES

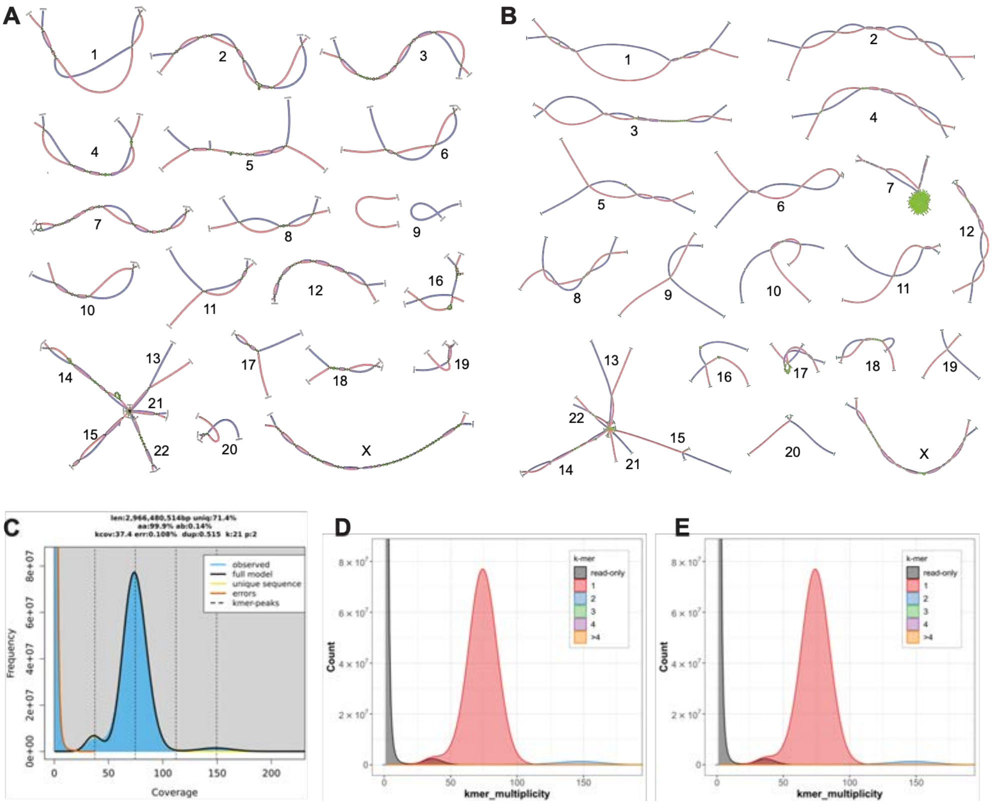


**Figure S1.** Bandage plots of 46 chromosomes of (**A**) asm1 and (**B**) asm2. The graphs were annotated using Hi-C data. The telomeres were marked at the end of chromosomes when present. The rDNA tangles were simplified to a single gap between chr13, chr14, chr15, chr21, and chr22. (**C**) GenomeScope2.0 analysis output for 21-mers computed using Meryl from HiFi reads. The diploid HiFi coverage is ~75x. (**D**-**E**) Merqury copy number spectrum plots for hap1 and hap2.

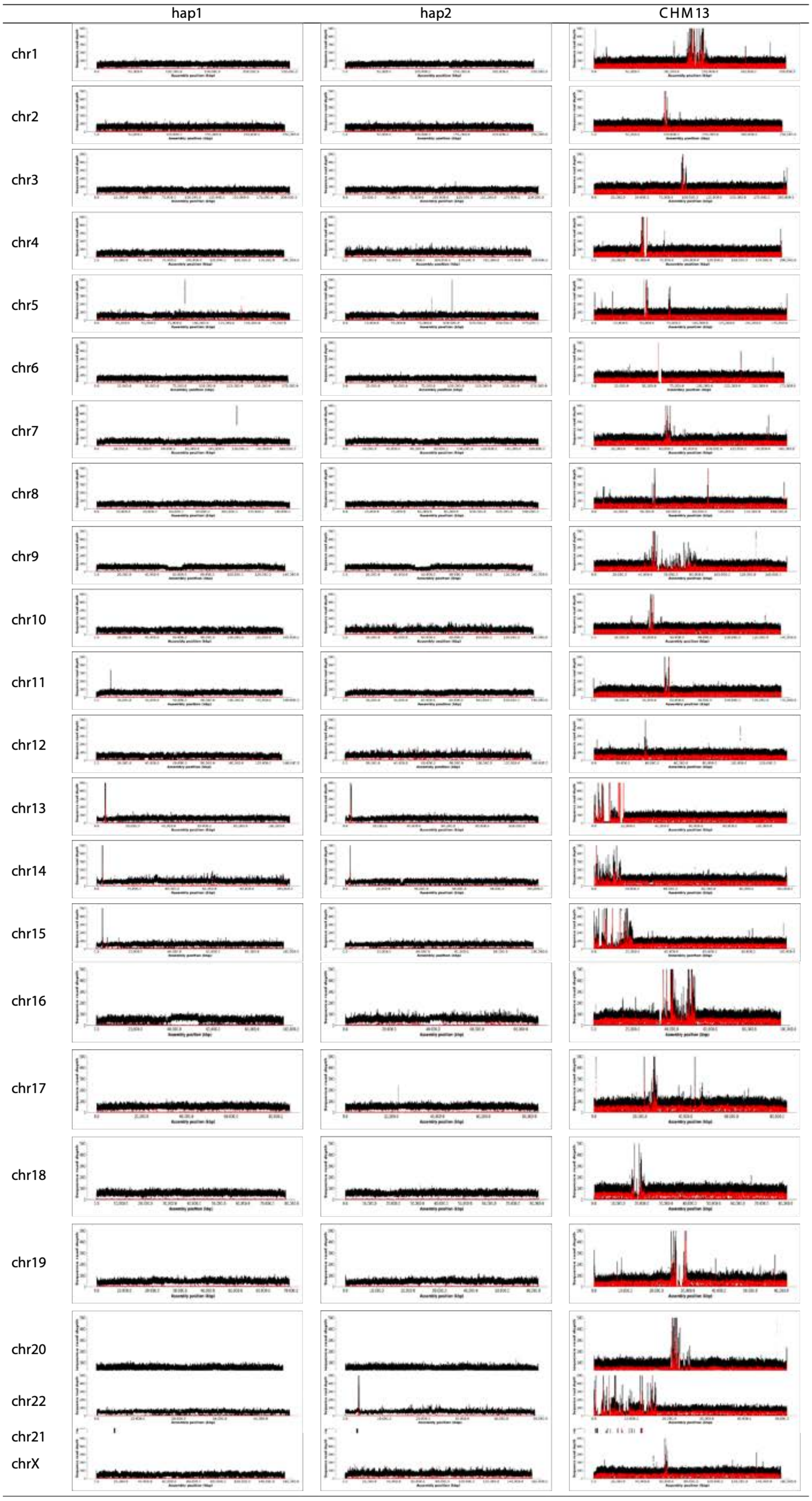


**Figure S2.** NucFreq HiFi coverage plots for all chromosomes. Coverage plots showed an overall homogeneous distribution across chromosomes in both haplotypes.

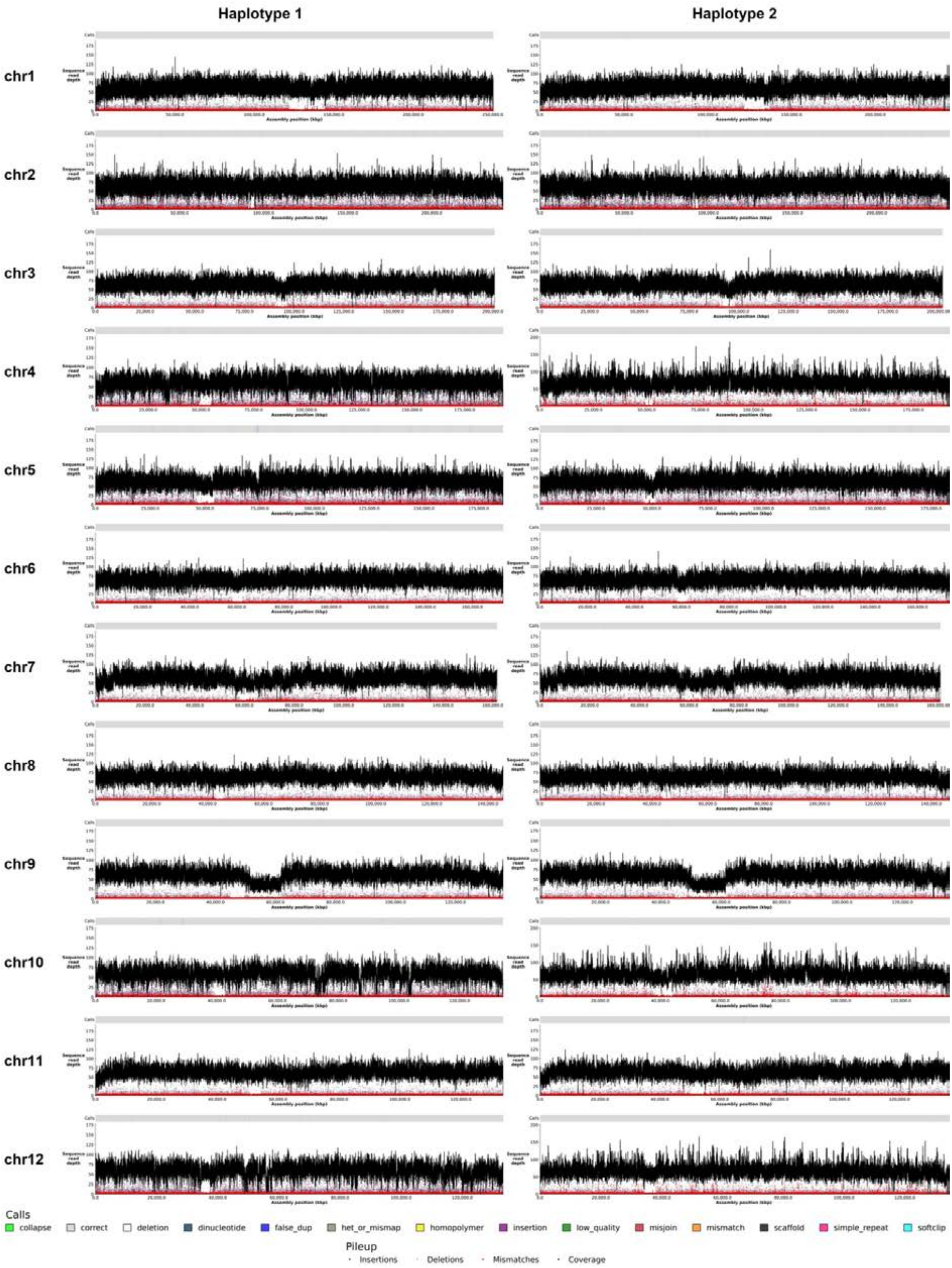


**Figure S3A**. NucFlag HiFi coverage plots on chromosome 1 to chromosome 12. Haplotype 1 is on the left and haplotype 2 is on the right.

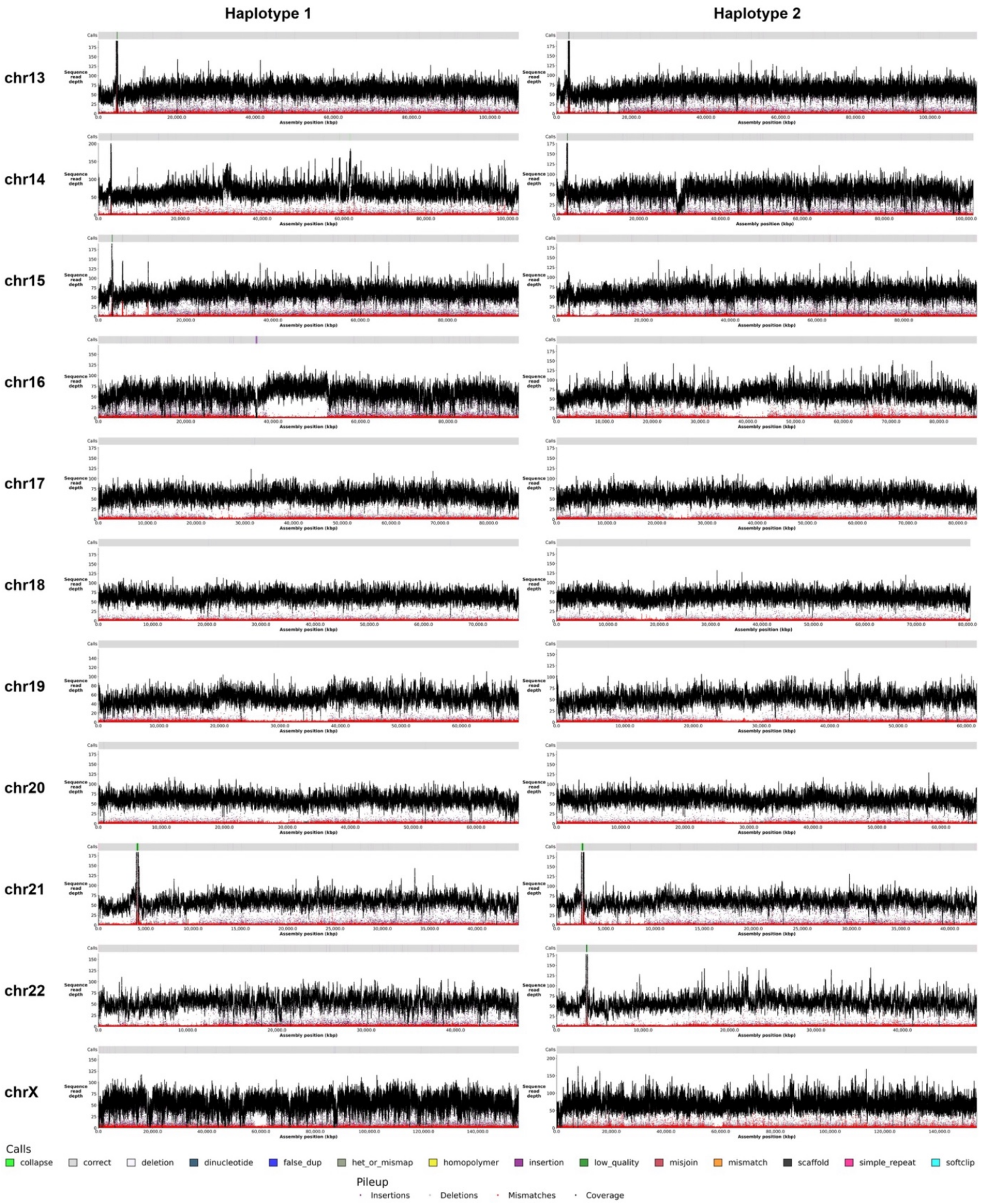


**Figure S3B** NucFlag HiFi coverage plots on chromosome 13 to chromosome X. Haplotype 1 is on the left and haplotype 2 is on the right.

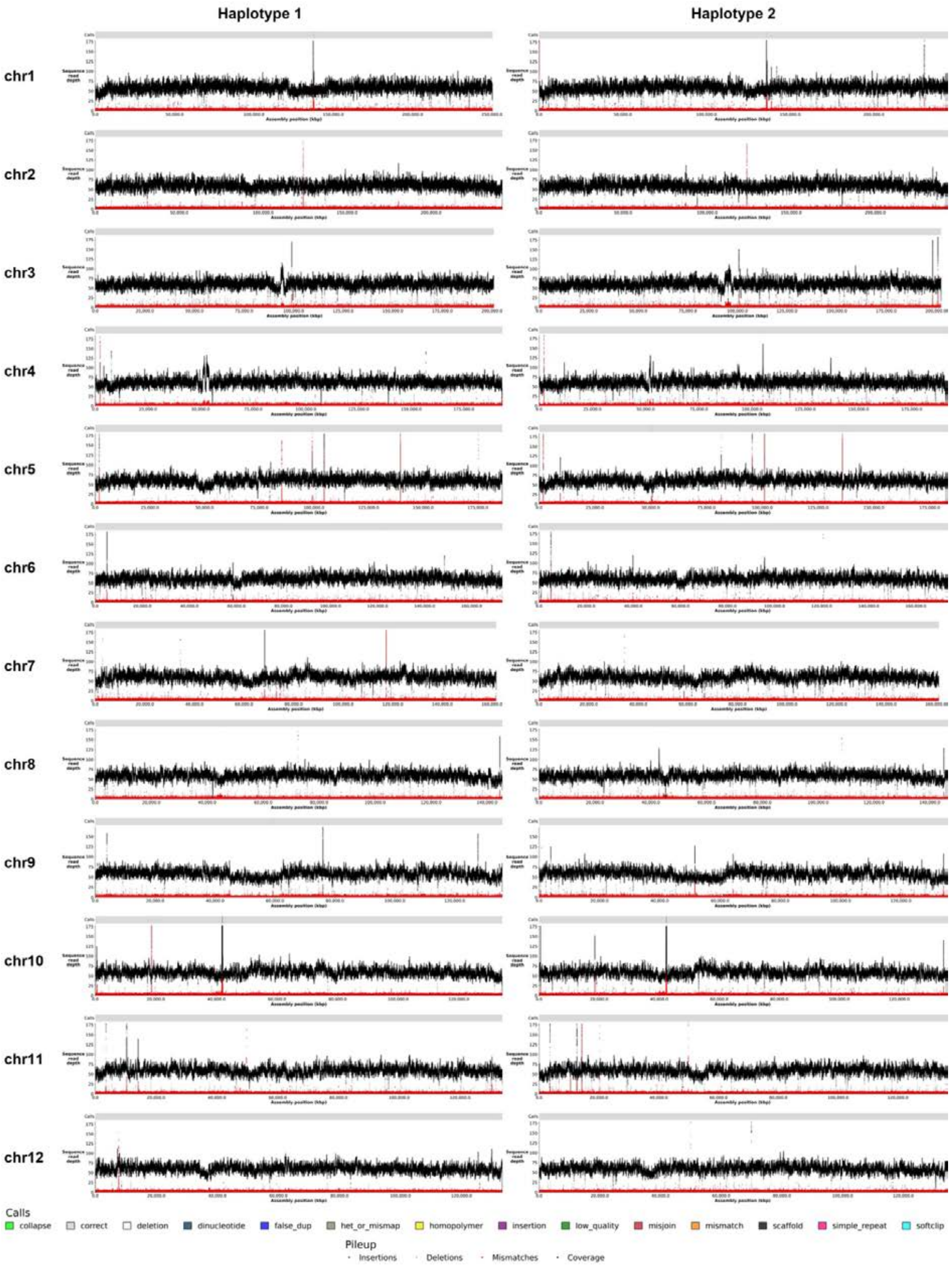


**Figure S4A**. NucFlag ONT coverage plots on chromosome 1 to chromosome 12. Haplotype 1 is on the left and haplotype 2 is on the right.

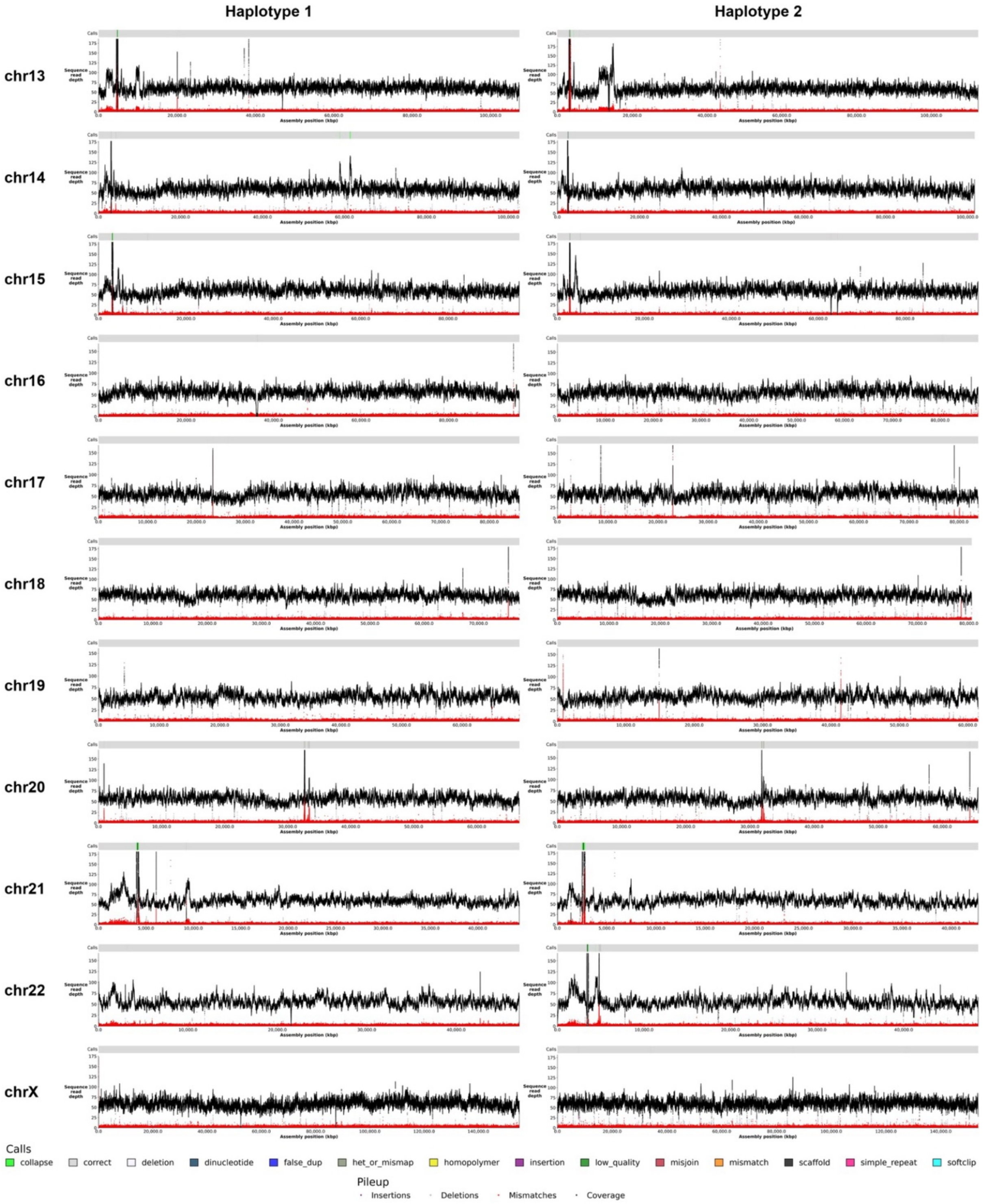


**Figure S4B**. NucFlag ONT coverage plots on chromosome 13 to chromosome X. Haplotype 1 is on the left and haplotype 2 is on the right.

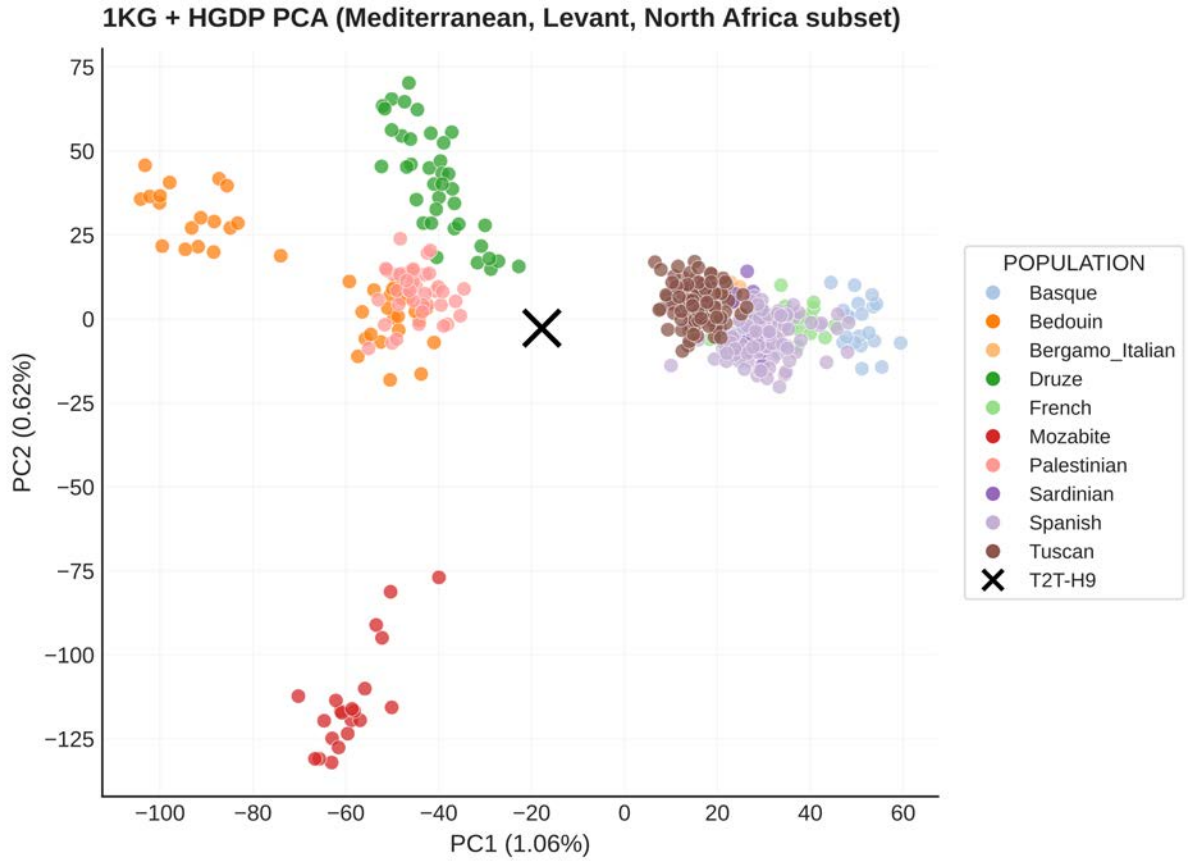


**Figure S5**. Principal component analysis of the 1KG and HGDP Mediterranean, Levantine, and North African samples with 300000 max-heterozygosity SNPs, colored by population. H9 is projected into this space and shown as a black cross. H9 falls within the Eastern Mediterranean continuum between European populations (Italian and Spanish on the right) and populations in the Levant (Bedouin, Palestinian, and Druze on the left), consistent either with admixture between the two or a source in an intermediate population along that cline. H9 has lower affinity with North African references (at bottom).

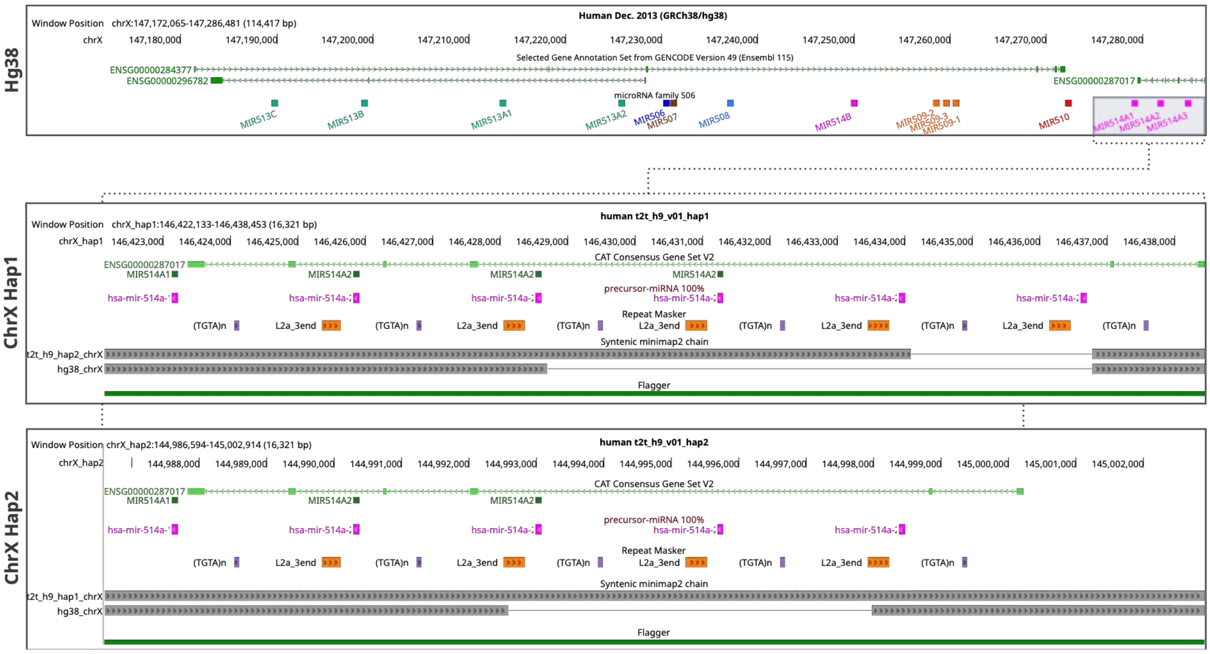


**Figure S6.** MicroRNA family 506 at chromosome X. The MIR-514 cluster at the end for this miRNA rich region shows different copy numbers of its members when comparing H9 to GRCh38 and even between haplotypes. The CNV is supported by high assembly accuracy as assessed by Flagger.

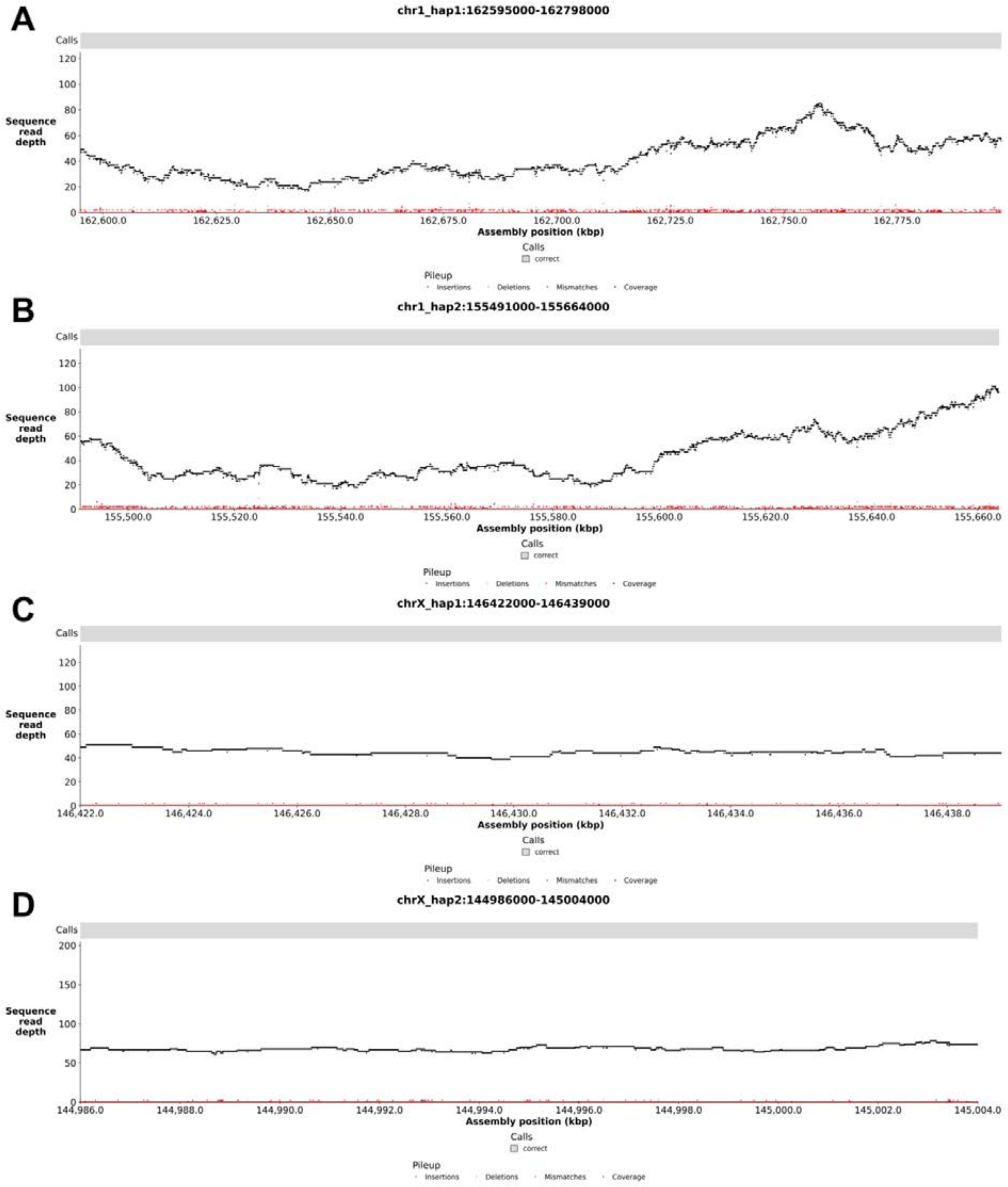


**Figure S7**. NucFlag HiFi coverage plots show well-supported regions on chromosome 1 in (**A**) haplotype 1 and (**B**) haplotype 2 with new tRNA genes compared to the GRCh38 genome, as well as the region in chromosome X of (**C**) haplotype 1 and (**D**) haplotype 2 with multi-copy MIR514A microRNA genes.

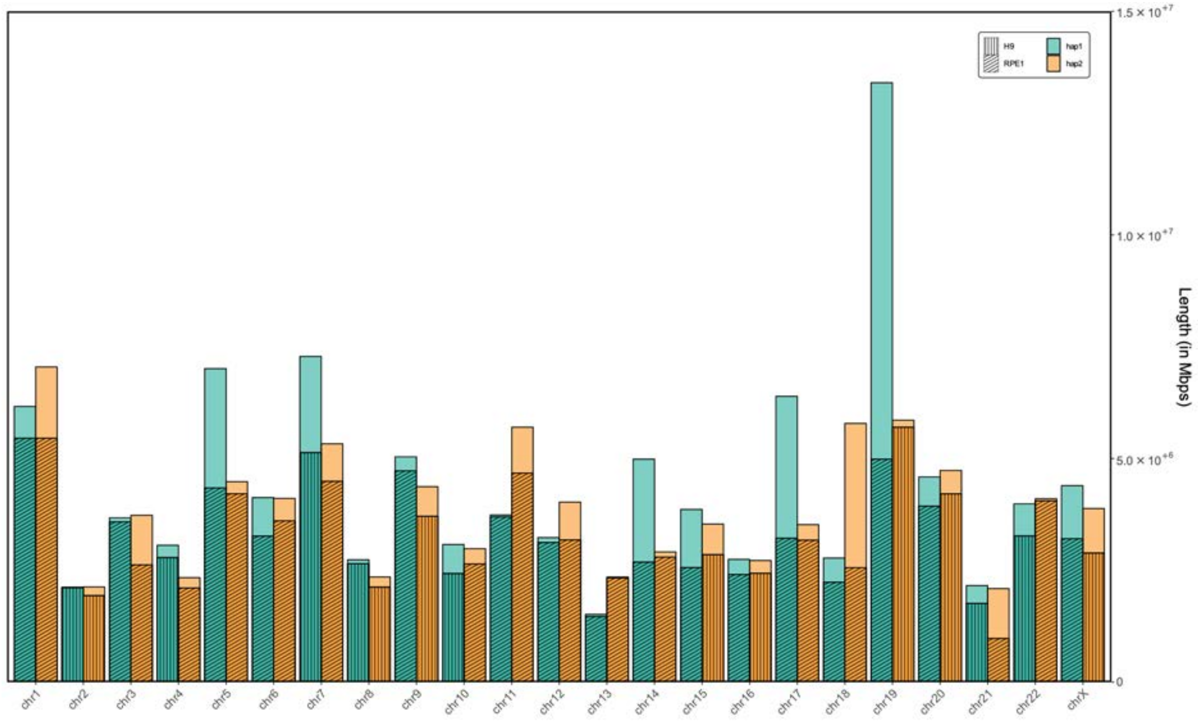


**Figure S8**. Centromeres' length comparison between the RPE1 and the H9 genome. In colors are presented both haplotypes for each assembly; the different patterns (which are assembly specific) are representative of the shorter centromere in the comparison. For instance, for chromosome 1, centromeres of RPE1 are both shorter than those in H9.

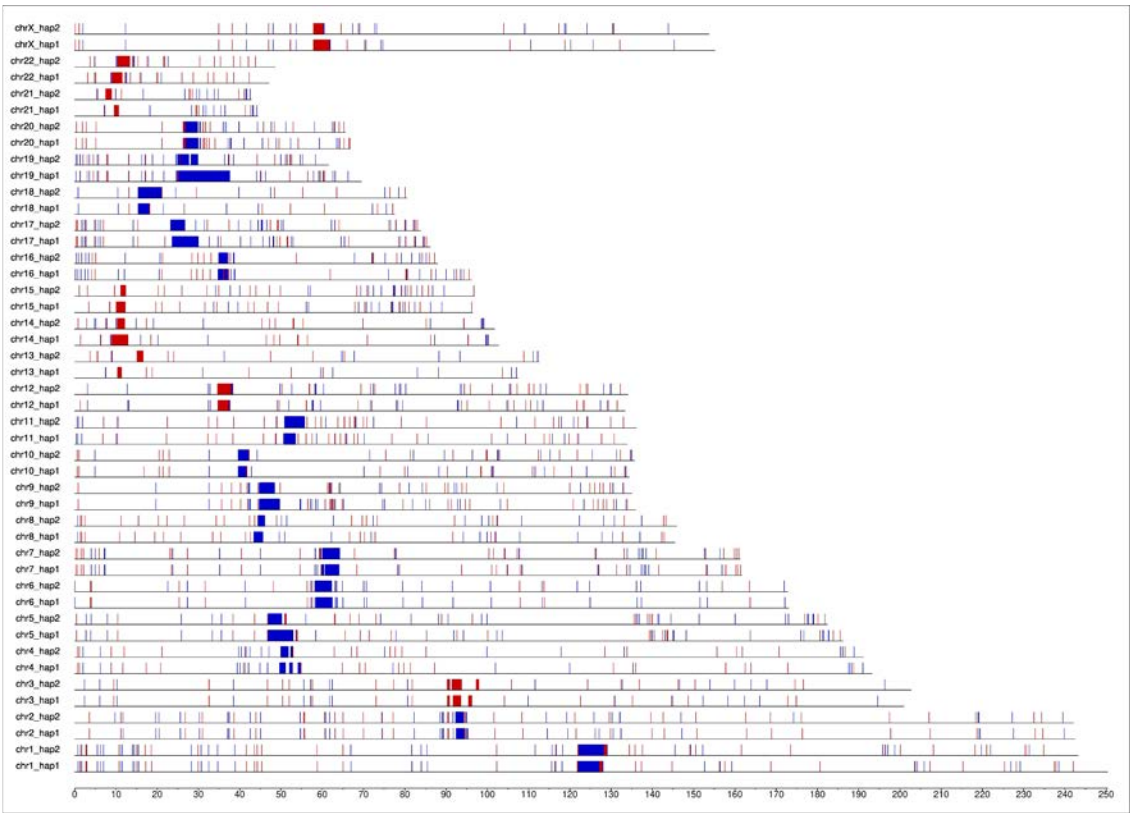


**Figure S9**. Centeny map of the H9 genome showcasing CENP-B box hits genome-wide for both haplotypes; the centromere of each chromosome is visible as a denser block of vertical bars indicating an enrichment of CENP-B box ligation sites. Strandeness is displayed with red (forward) and blue (reverse) colors.

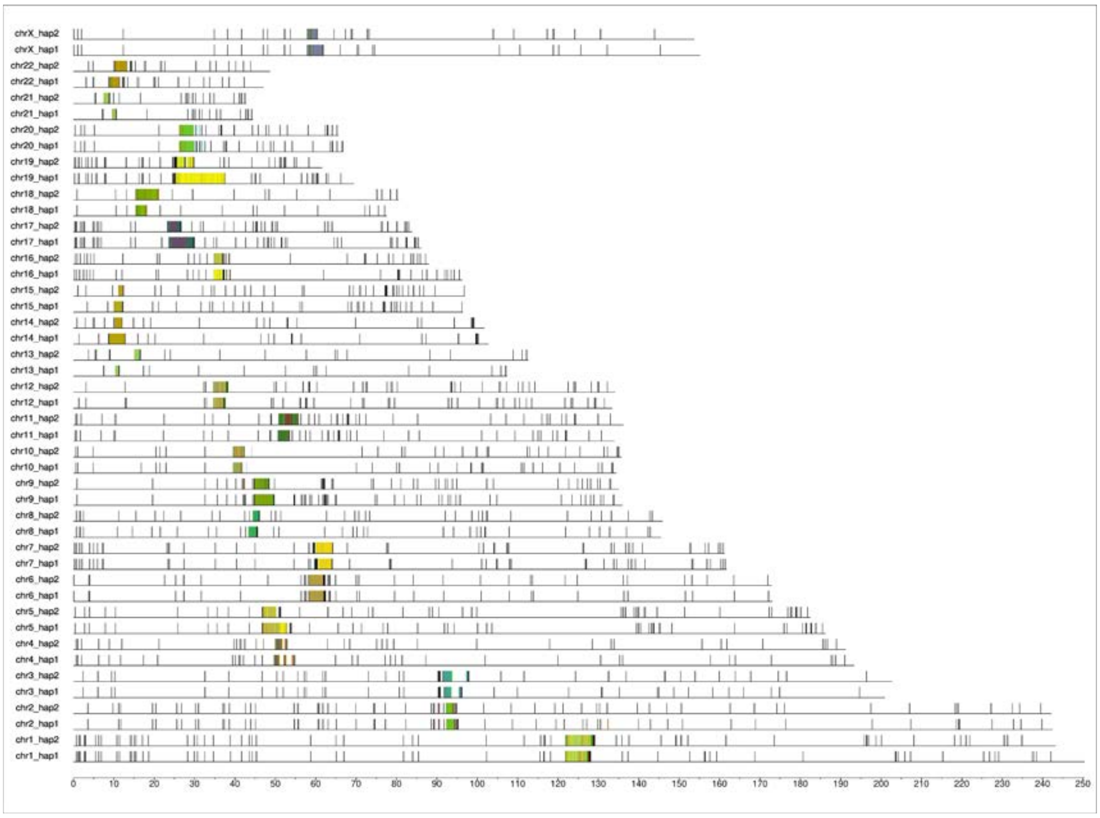


**Figure S10**. Model 1 of the GCP pipeline expressing CENP-B box distances genome-wide which is the distance in base pairs between two consecutive boxes. Color shades from lighter to darker indicate greater distances.

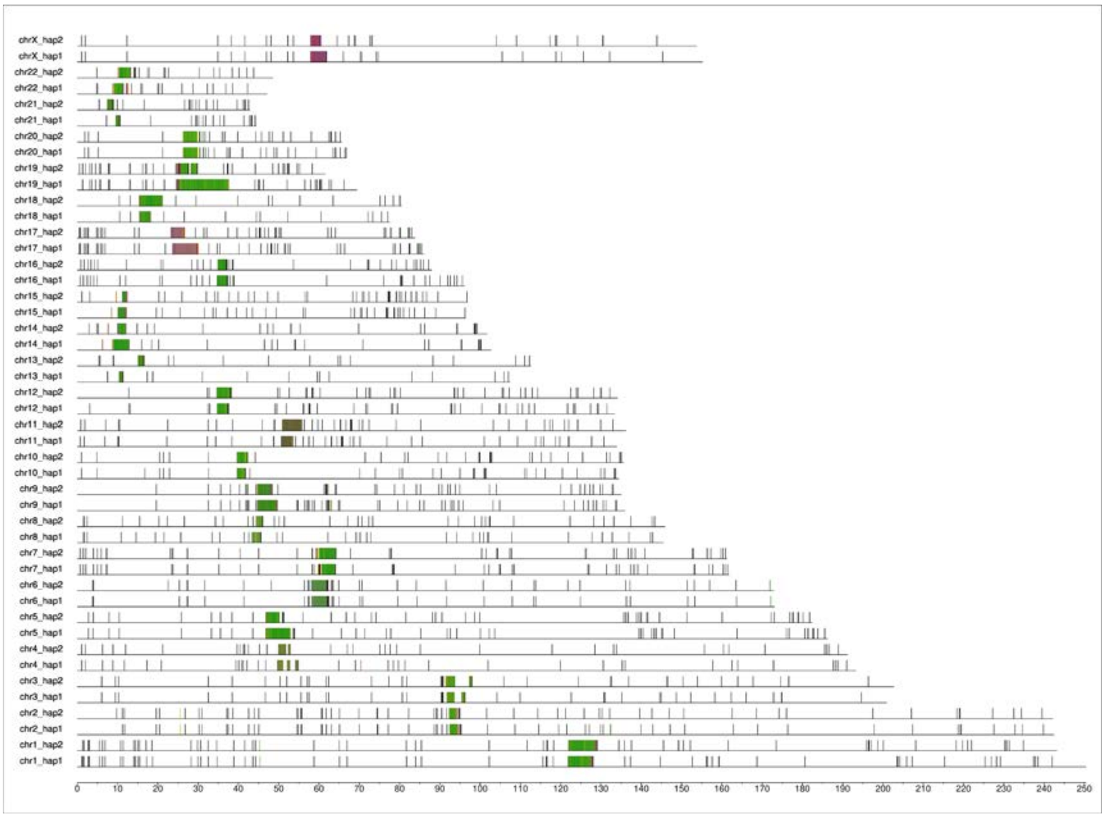


**Figure S11**. Model 2 of the GCP pipeline expressing CENP-B box motifs occurrence genome-wide, these represent for the periodicity at which a specific box is located according to the next one and whether or not they place on consecutive monomers. Color shades from lighter to darker indicate more interspersed occurrences.

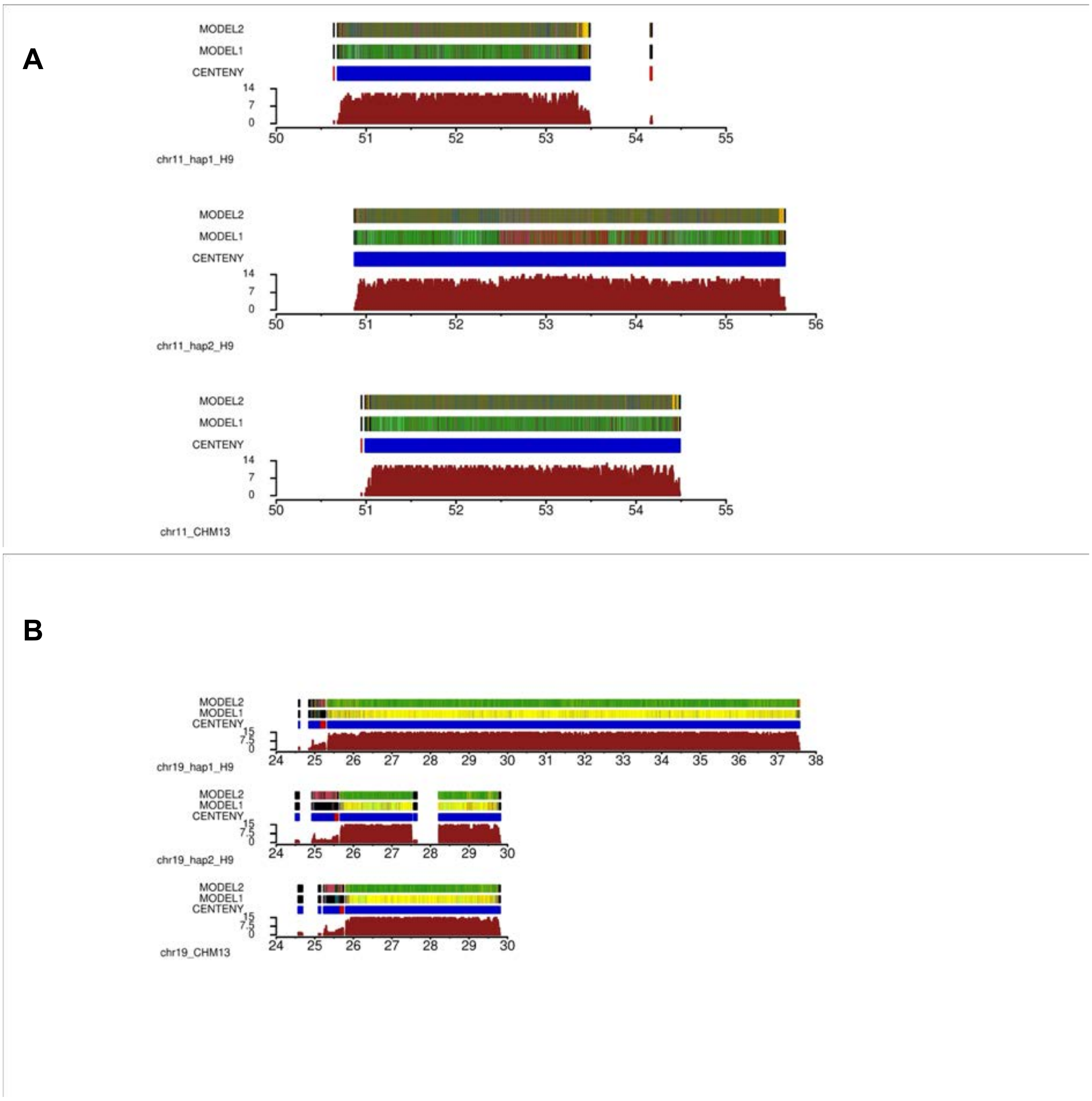


**Figure S12**. Centromere features for select chromosomes. Zoom-in of the Centeny map (bottom), Model 1 (middle), and Model 2 (top) for the centromeres of chromosome 11 (**A**) and 19 (**B**) of the H9 genome (both haplotypes). The CHM13 reference is added below for comparison. Note the change in motif composition of chromosome 11 haplotype 2 and the exceedingly long centromere of chromosome 19 haplotype 1 compared to both the complementary haplotype and CHM13.

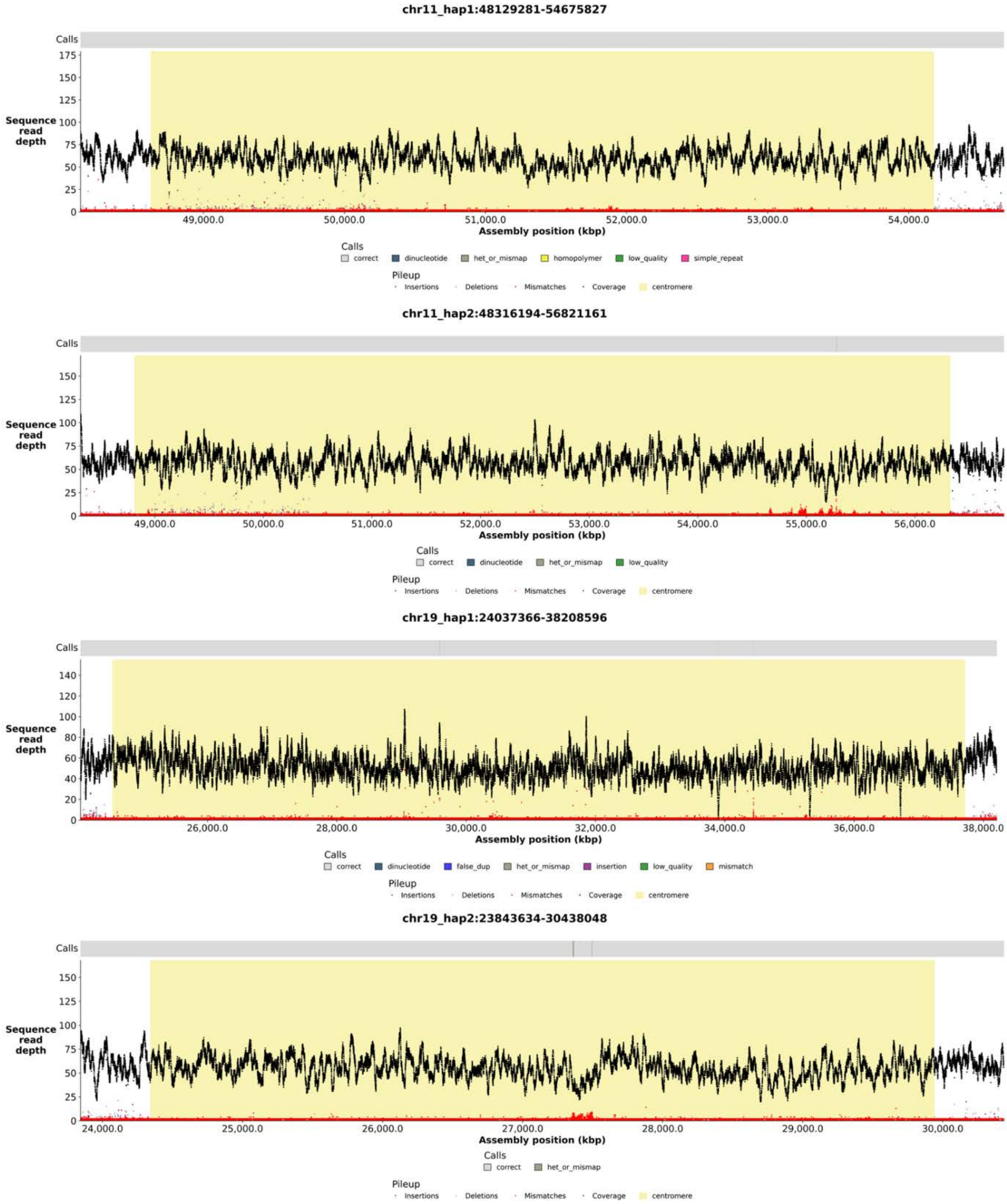


**Figure S13.** NucFlag HiFi coverage plots for centromeric regions (highlighted in yellow) of chr11 and chr19 in both haplotypes.

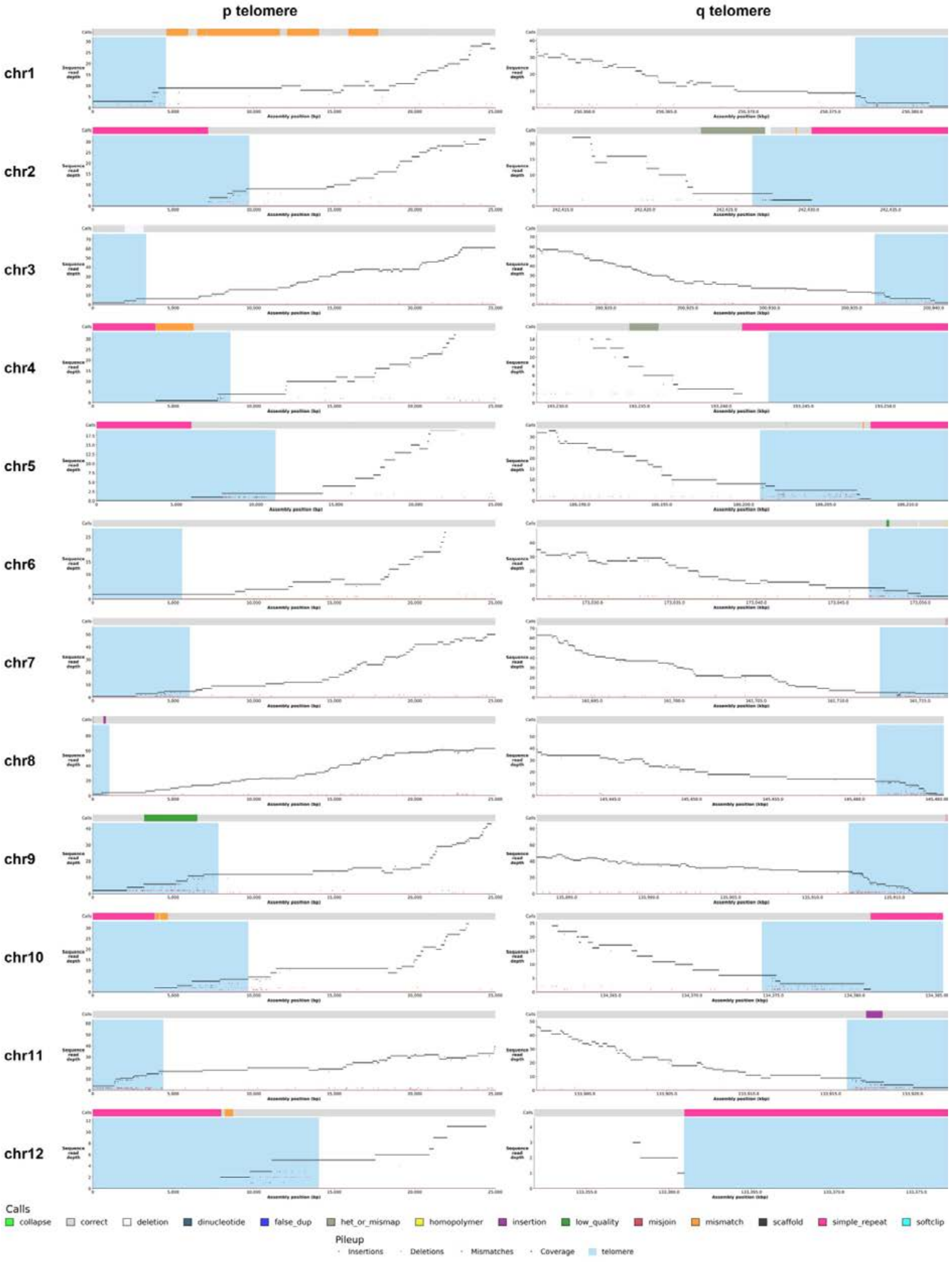


**Figure S14A**. Nucflag HiFi coverage plots for p and q telomeric regions from chromosomes 1 to 12 of haplotype 1 assembly. Telomeres are highlighted in blue.

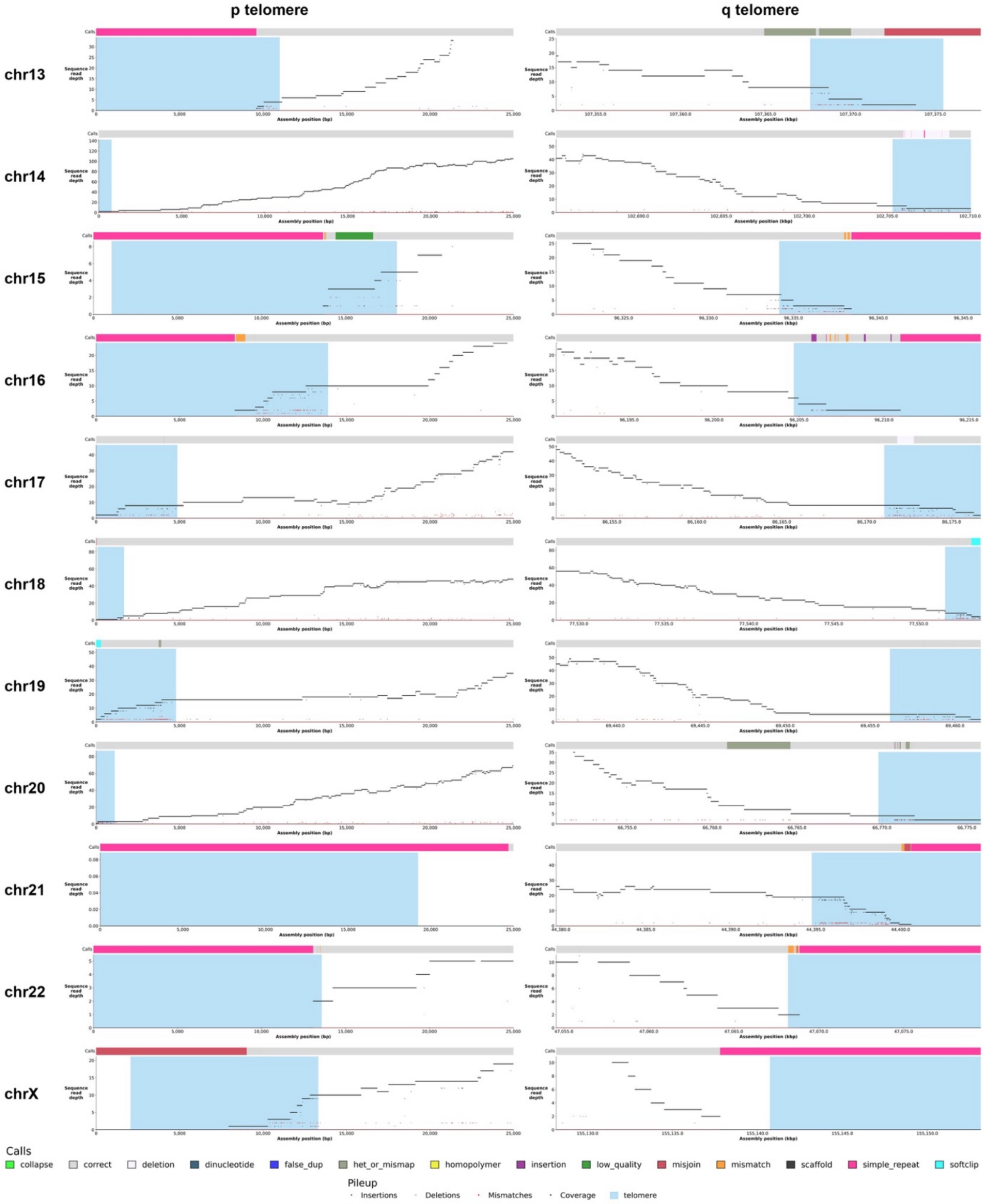


**Figure S14B**. Nucflag HiFi coverage plots for p and q telomeric regions from chromosome 13 to X.

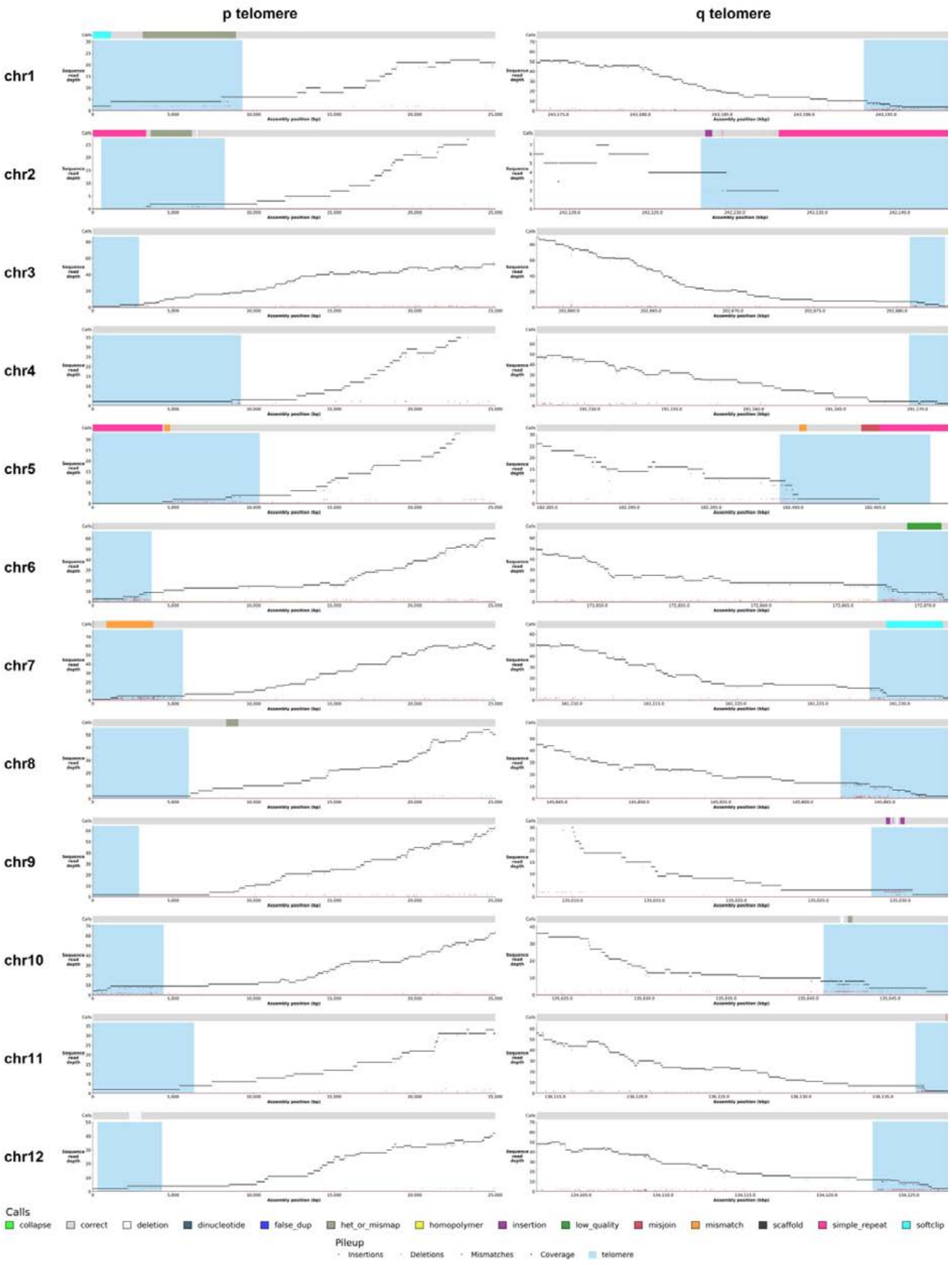


**Figure S15A**. Nucflag HiFi coverage plots for p and q telomeric regions from chromosomes 1 to 12 of haplotype 2 assembly. Telomeres are highlighted in blue.

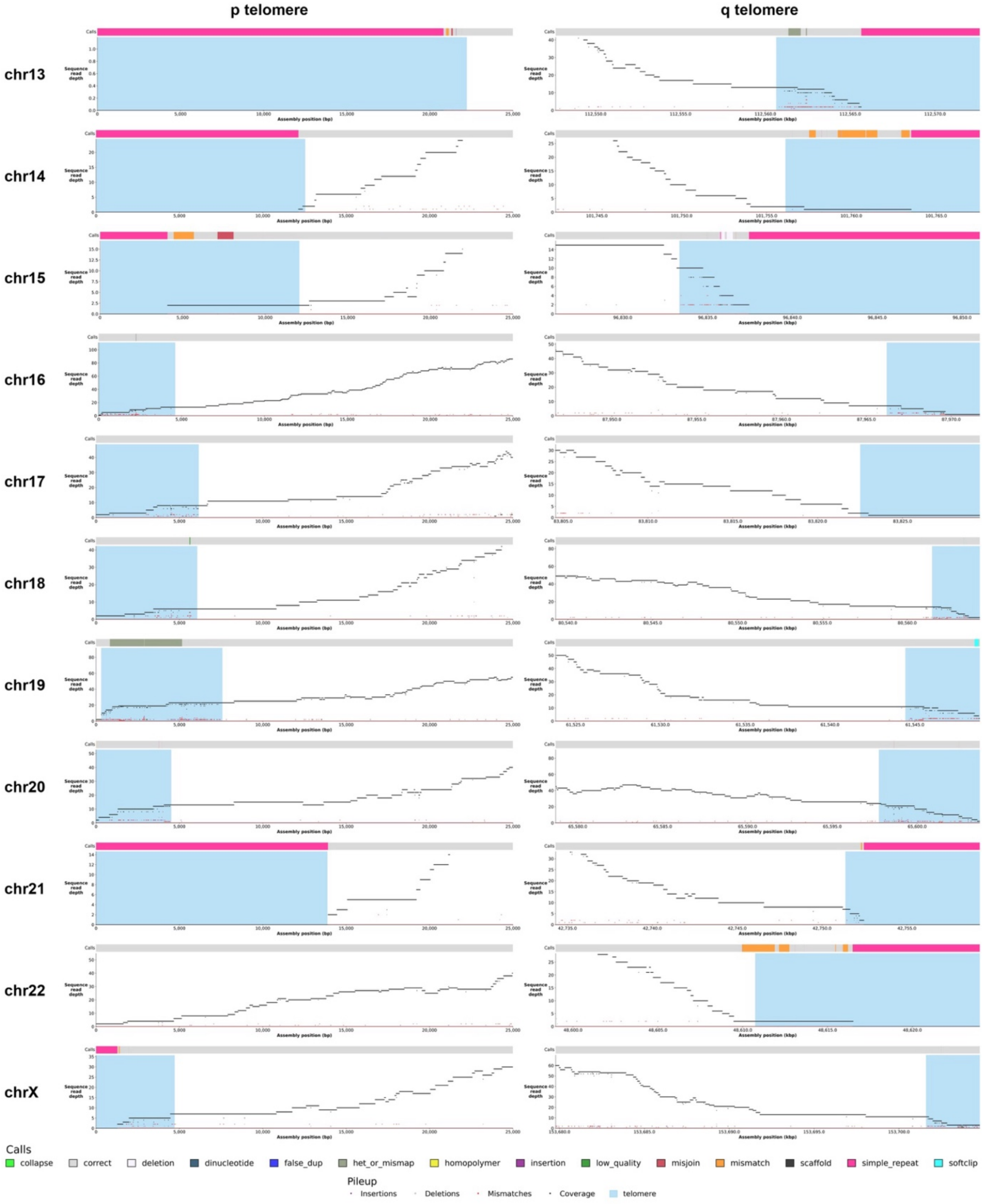


**Figure S15B**. Nucflag HiFi coverage plots for p and q telomeric regions from chromosome 13 to X.

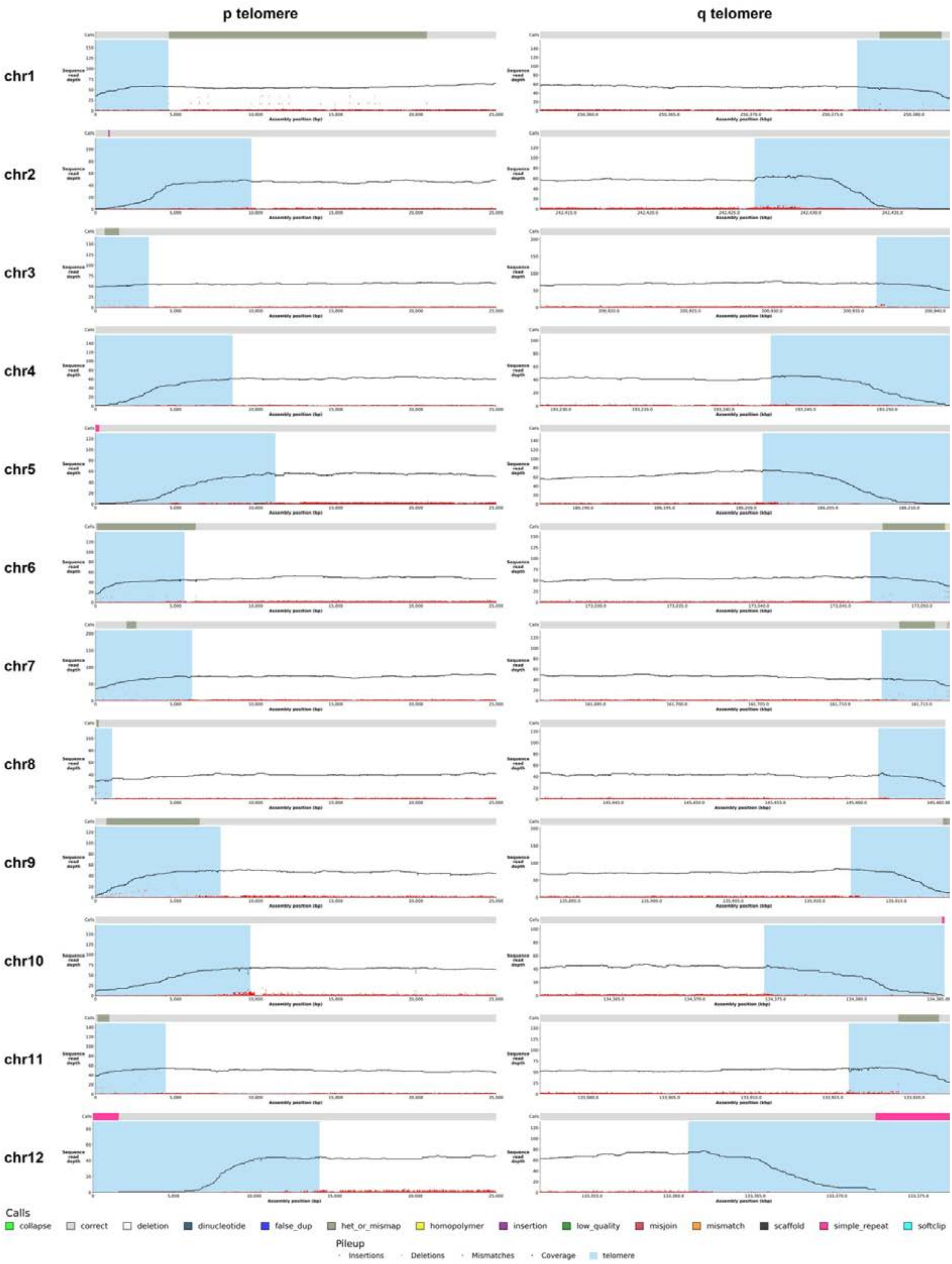


**Figure S16A**. Nucflag ONT coverage plots for p and q telomeric regions from chromosomes 1 to 12 of haplotype 1 assembly. Telomeres are highlighted in blue.

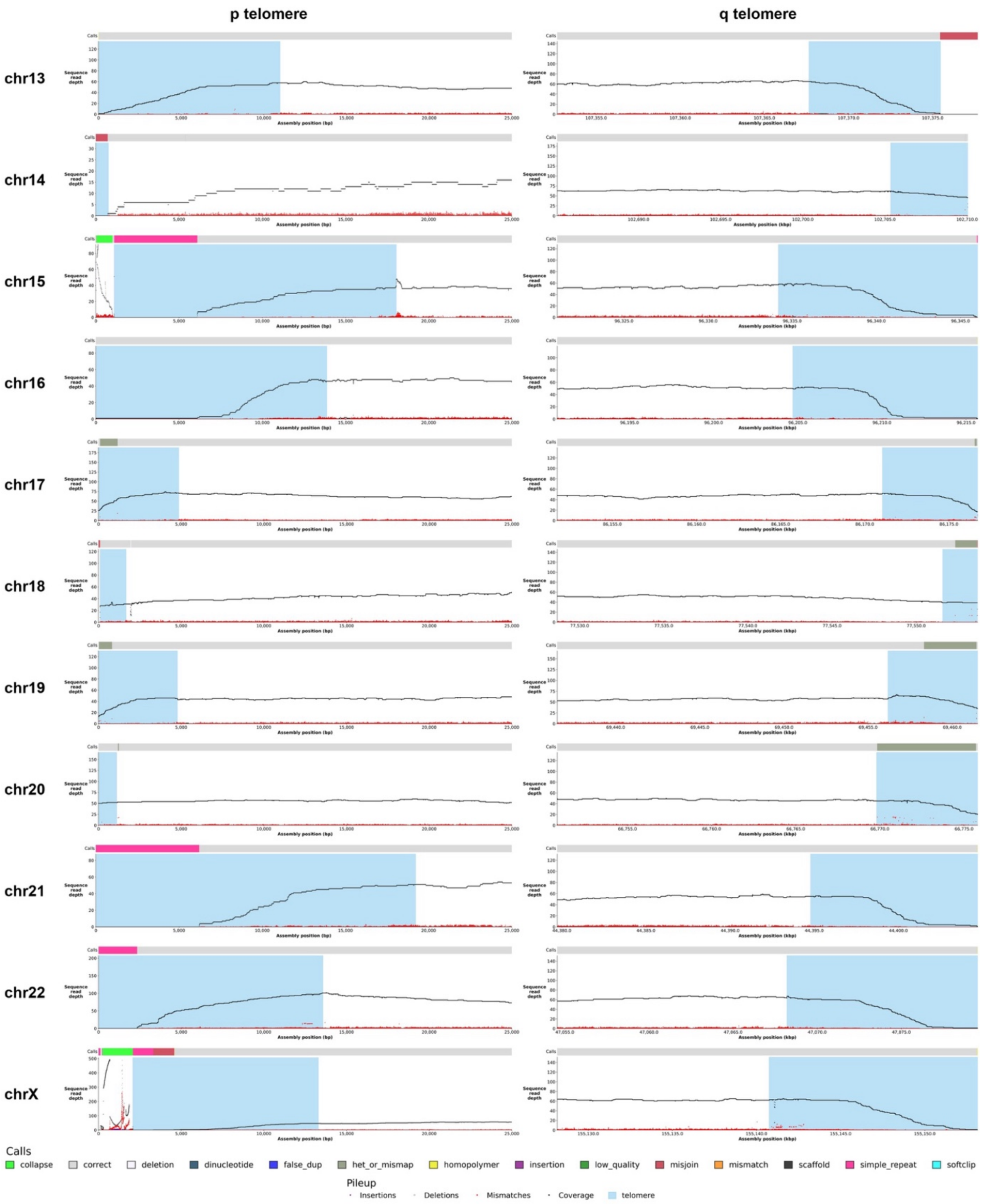


**Figure S16B**. Nucflag ONT coverage plots for p and q telomeric regions from chromosome 13 to X.

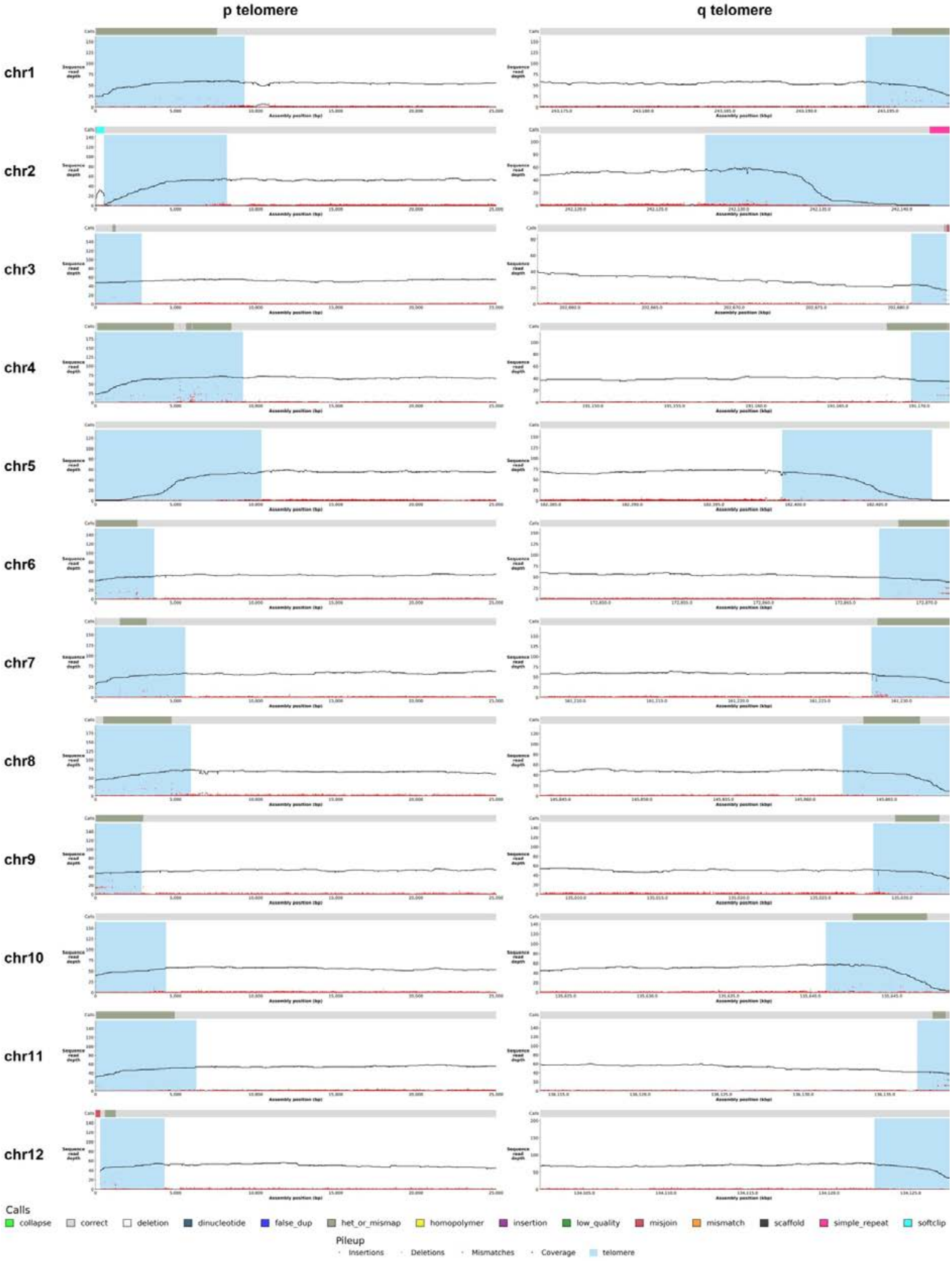


**Figure S17A**. Nucflag ONT coverage plots for p and q telomeric regions from chromosomes 1 to 12 of haplotype 2 assembly. Telomeres are highlighted in blue.

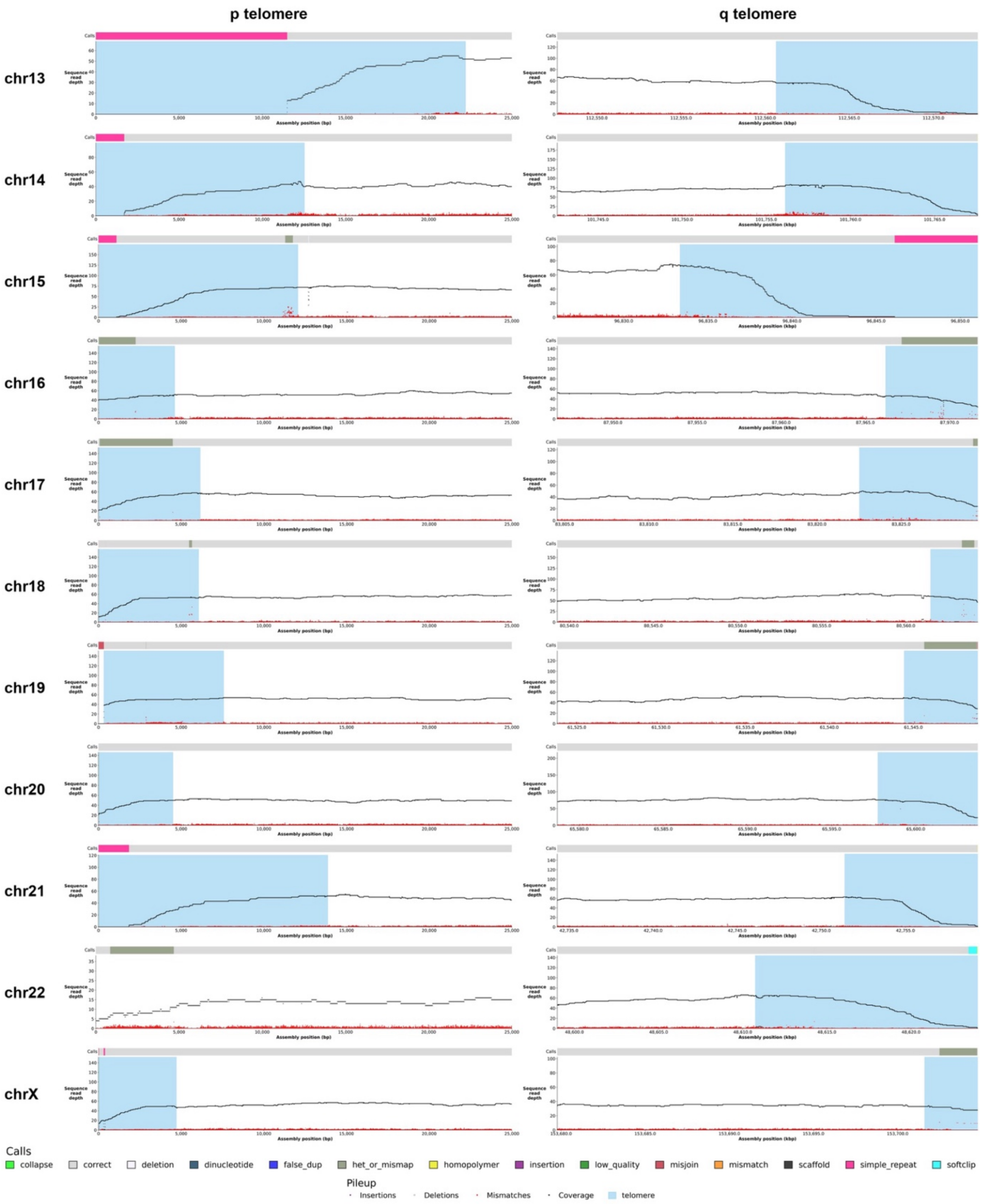


**Figure S17B**. Nucflag ONT coverage plots for p and q telomeric regions from chromosome 13 to X.

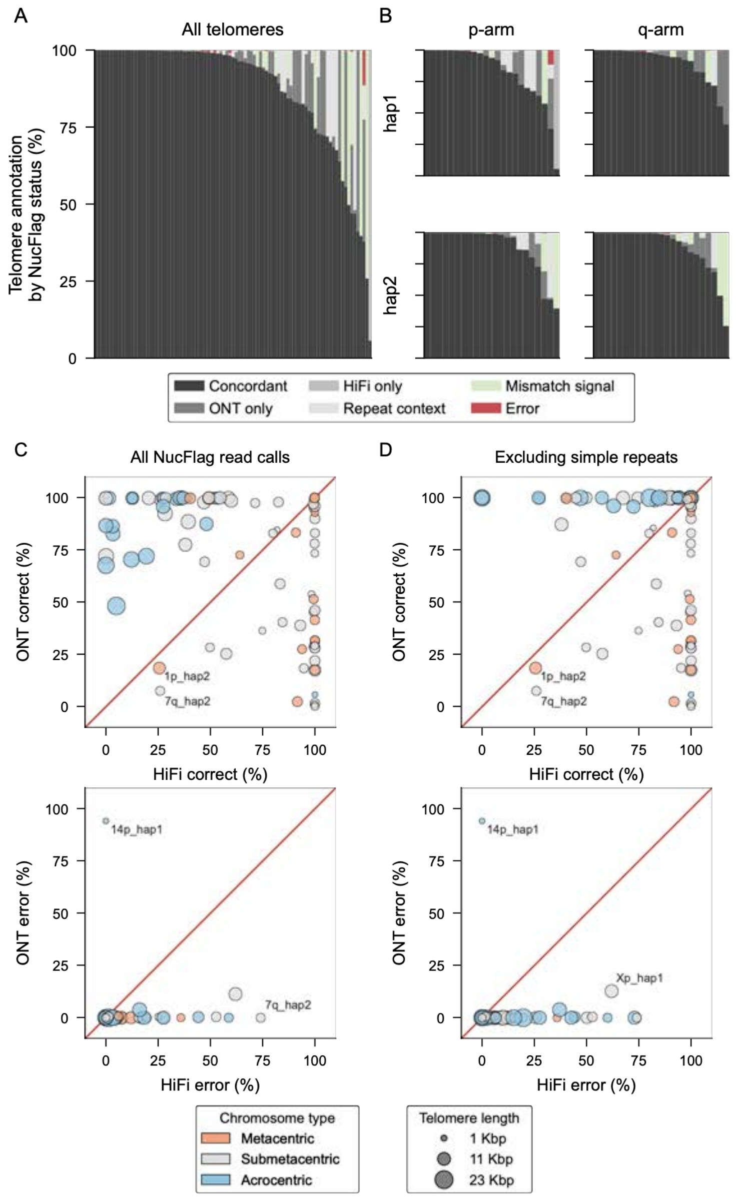


**Figure S18.** Complementary telomere-end validation in H9 by NucFlag using HiFi and ONT reads. (**A**) Percentage of each annotated telomere by bases assigned to concordant support (both technologies support the assembly), technology-specific support (ONT or HiFi only), repeat context relabels (homopolymers, dinucleotides, simple repeats, other repeats), mismatch signal (heterozygous or mis-mapped or low quality), or error (in both technologies). (**B**) Same as (**A**), stratified by chromosome arm (p or q) and haplotype (hap1 or hap2). (**C**) Percentage of each telomere labelled “correct” and “error” by ONT versus HiFi, computed either across all telomeric bases (left) or (**D**), after excluding repeat-context segments (right). In both cases, the red diagonal indicates equality between technologies, point color denotes chromosome types (acrocentric, submetacentric, metacentric), and point size scales with telomere length (Kbps).

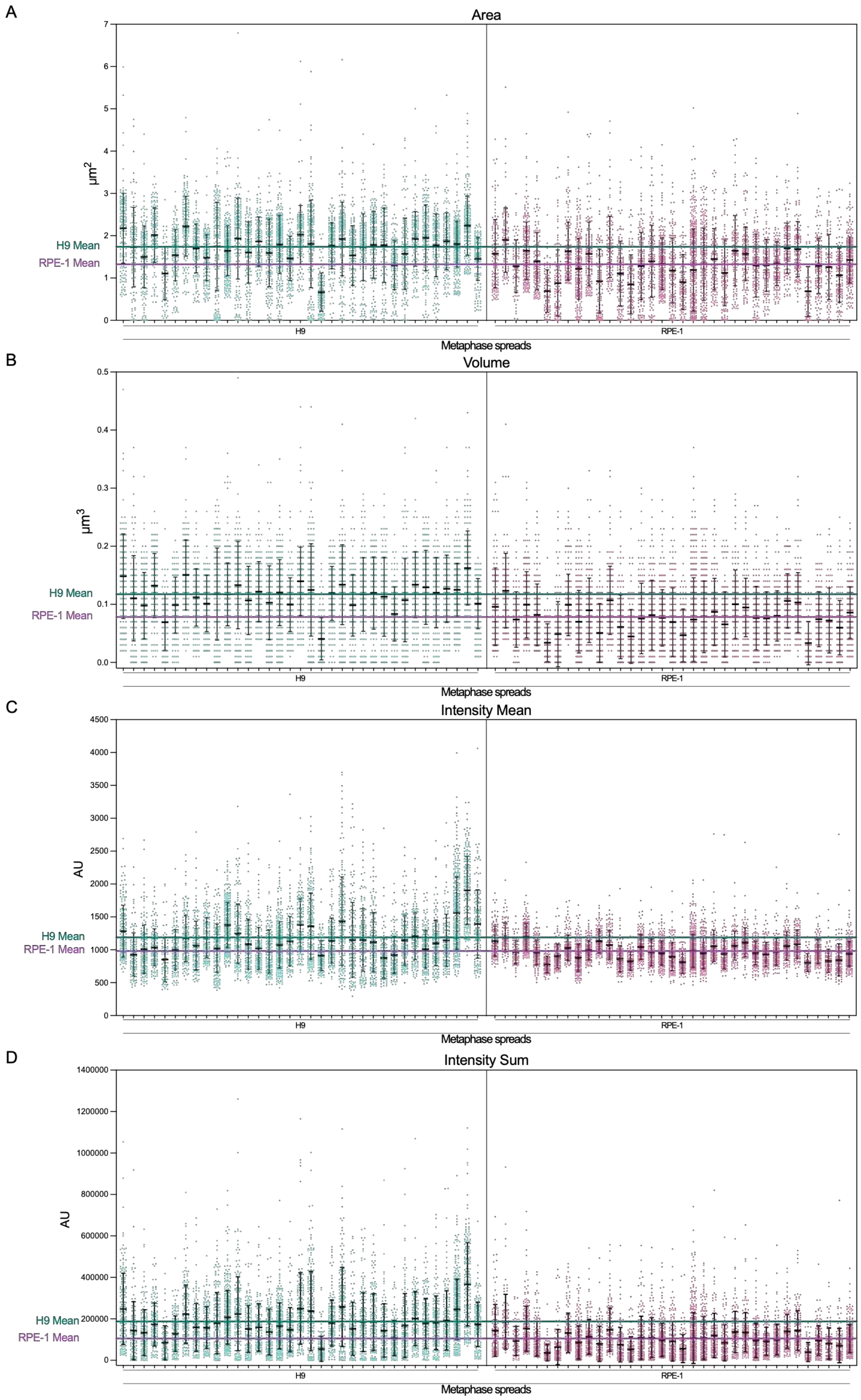


**Figure S19.** Analysis of Area (**A**), Volume (**B**), Intensity mean (**C**), and Intensity sum (**D**) of each telomere from 35 human H9 and 35 human RPE-1-hTERT metaphases stained by FISH. The mean values highlighted with the colored lines were obtained from the plots in **Figure 4C**. AU = arbitrary units.

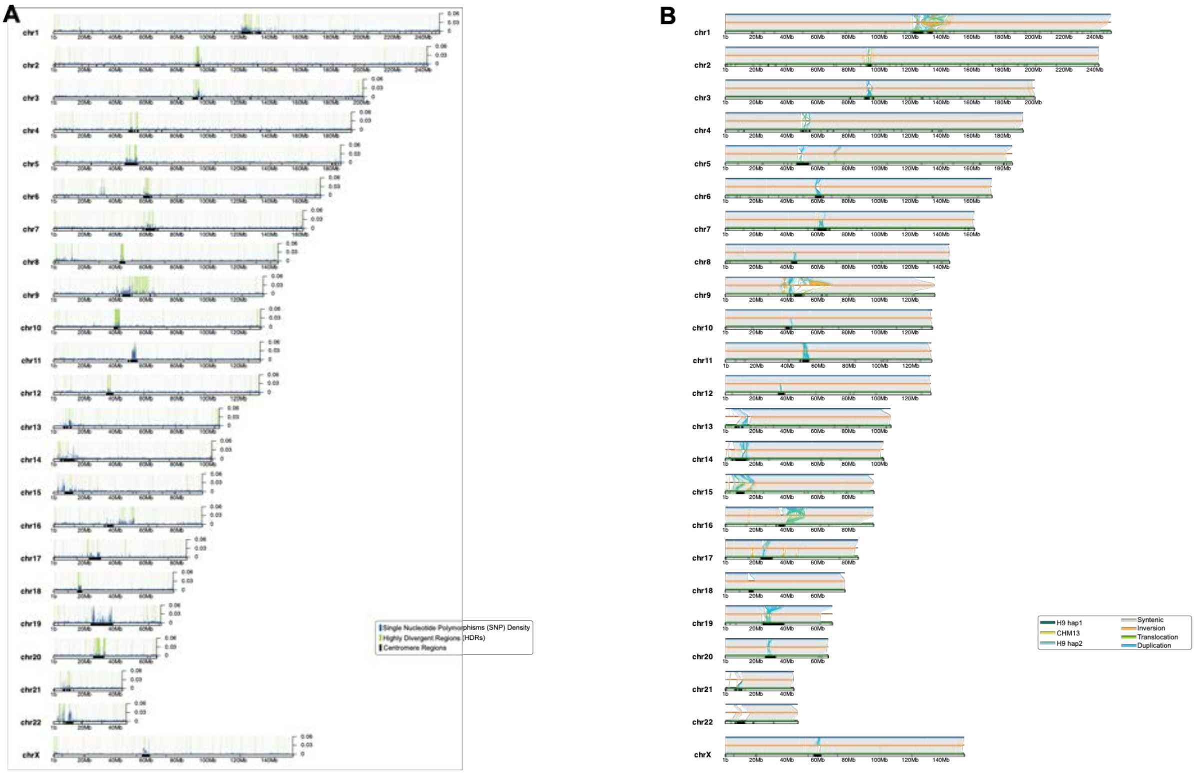


**Figure S20.** Individual chromosomes (**A**) SNVs density and (**B**) SyRI structural rearrangements as per the main **Figure 5A-B**.

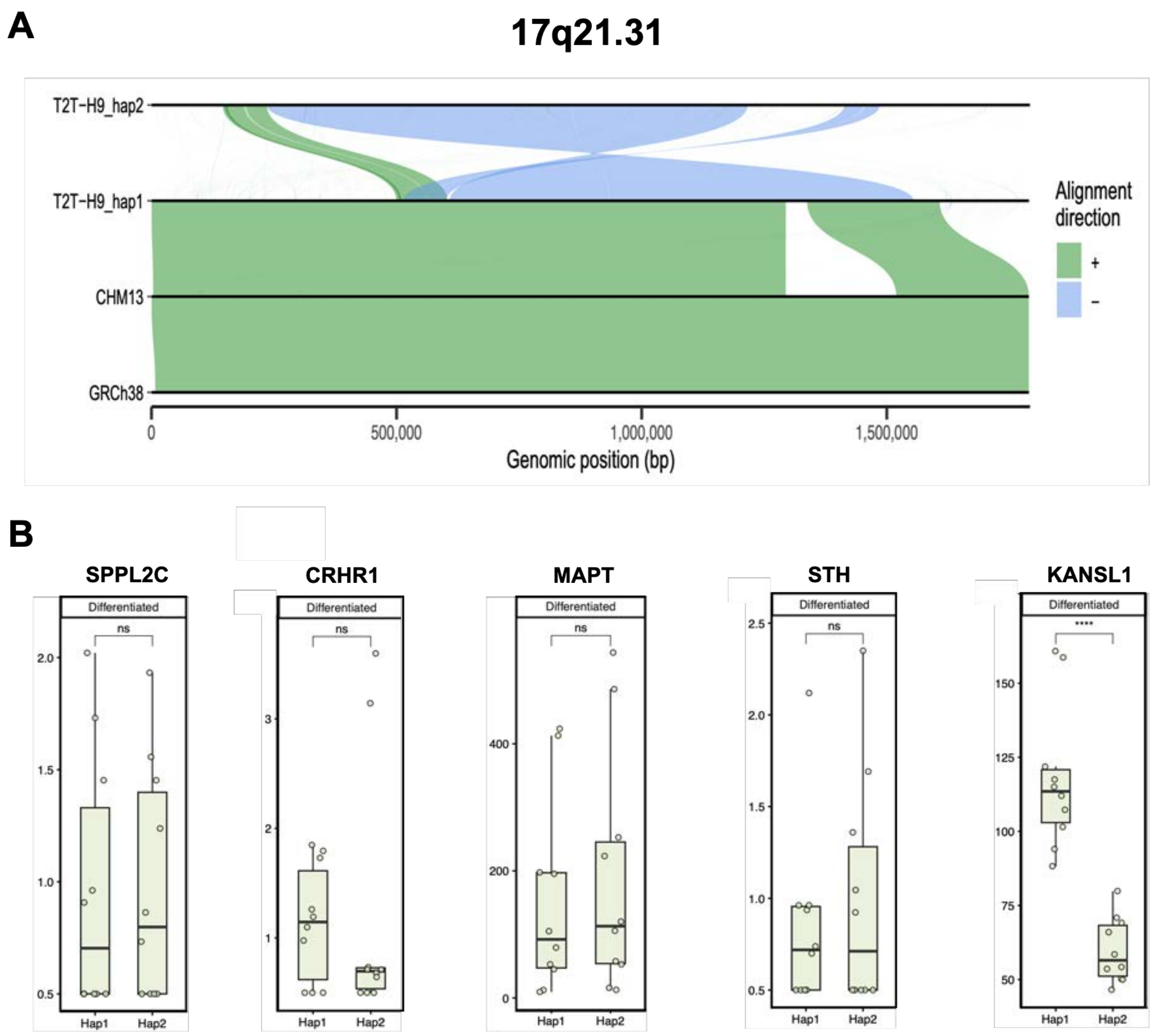


**Figure S21. (A)** SVByEye plot demonstrating the 17q21.31 inversion in H9 haplotype 2. **(B)** DESeq2 RNA-Seq analysis of five known causal genes in the 17q21.31 pathogenesis, demonstrating that KANSL1 has an increased level of expression in haplotype 1.

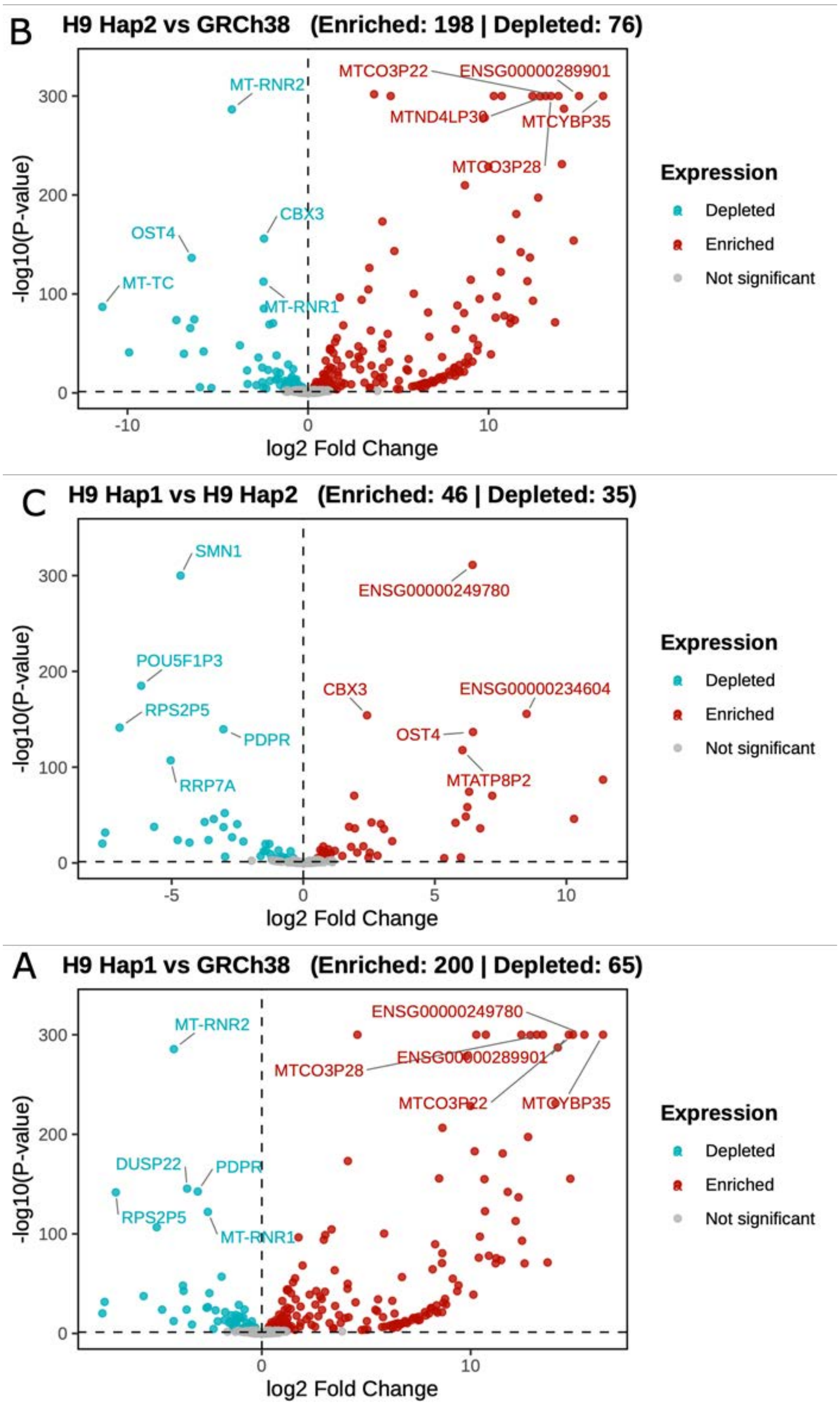


**Figure S22.** Impact of reference genome selection on transcriptomics analysis. Volcano plots displaying the log2 fold change (FC) and statistical significance [-log10(p-value)] of gene mapping differences: (**A**) H9 hap1 vs. GRCh38, (**B**) H9 hap2 vs. GRCh38, and (**C**) H9 hap1 vs. H9 hap2. In each pairwise comparison, significantly differentially mapped genes (FDR < 0.05, log2FC < 0 or FDR < 0.05, log2FC > 0) genes are shown in blue and red for each haplotype comparison as indicated in the legend. The top 5 enriched and depleted genes in each pairwise comparison are labeled.

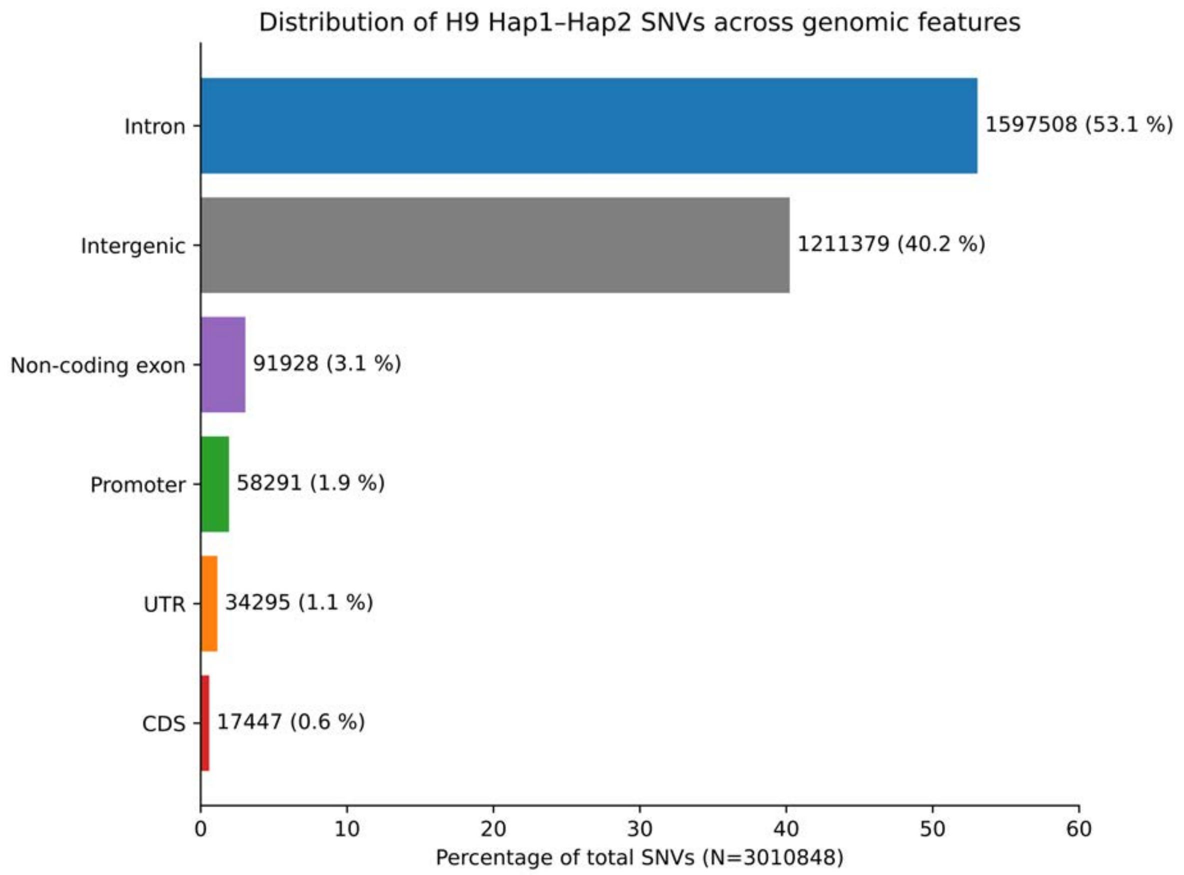


**Figure S23.** Distribution of SNVs between H9 haplotypes across genomic features. **CDS** Coding DNA Sequence, **UTR** Untranslated Region.

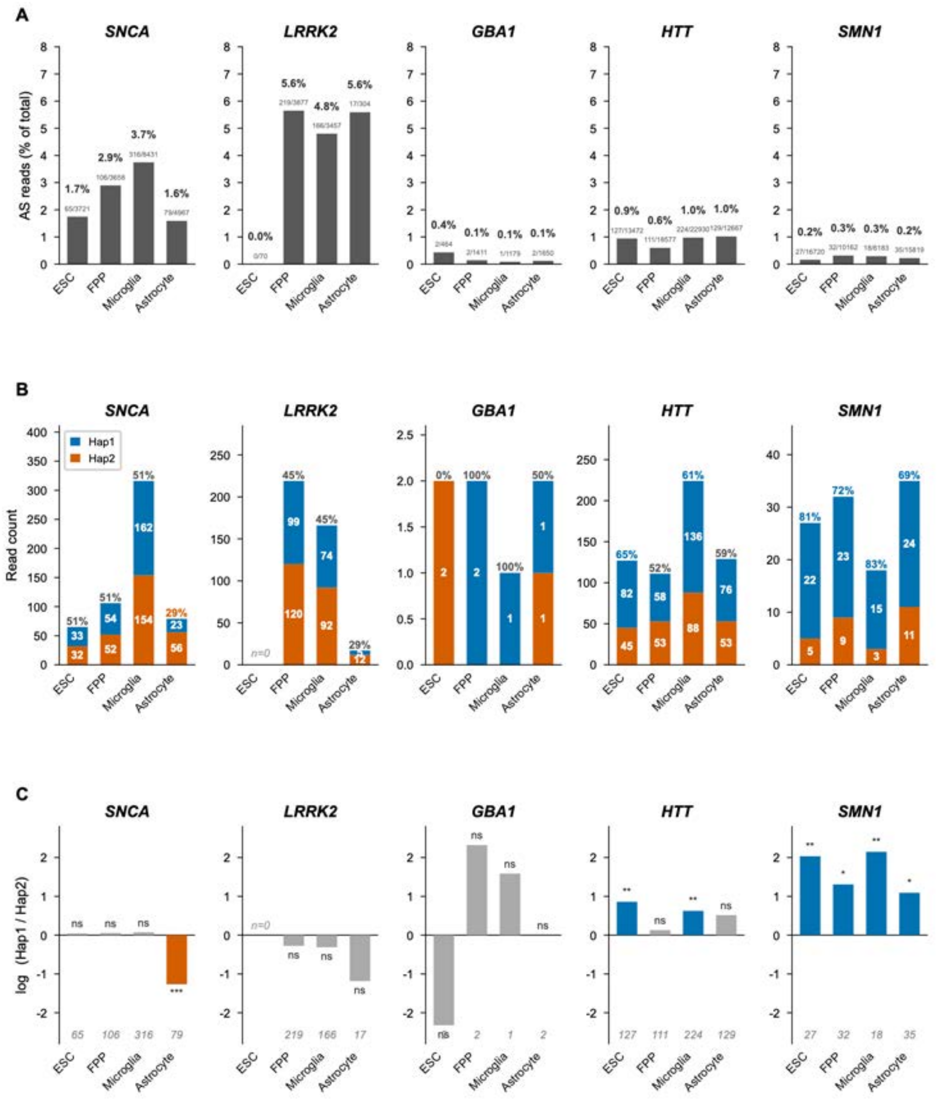


**Figure S24.** Haplotype-specific allelic expression of neurodegeneration-associated genes across H9-derived neural lineage cell types. (**A**) Allele-specific reads, and total reads per gene body were pulled from UCSC Hap1 and Hap2 ASE and diploid tracks for each cell type: pluripotent (ESC, day0), midbrain floor plate progenitors (FPP, day33), and astrocytes (day125) and the % ASE reads computed for each cell type and gene. Numbers above bars indicate the ratio of phased AS reads to total reads. (**B**) Total ASE reads (y axis) from Hap1 and Hap 2 show the balance of allelic expression, % Hap1 of total above bars, SNCA is balanced in ESC and FPP, with a cell-type-specific shift to Hap2-dominant expression in astrocytes. HTT shows Hap1-dominant expression in ESC. SMN1 shows consistent Hap1-dominant expression across all cell types. (**C**) $\log_2$(Hap1/Hap2) allelic bias. Bar color indicates direction of significant bias (blue, Hap1-dominant; orange, Hap2-dominant; gray, not significant). SNCA expression was significantly shifted to Hap2 from a 50:50 baseline only in astrocytes, whereas SMN1 showed significant SMN1 Hap dominance in all cell types. Total phased read counts are indicated at the base of each bar. Significance was determined by a two-sided binomial test against equal allelic expression ($p = 0.5$); *$p < 0.05$, **$p < 0.01$, ***$p < 0.001$, ns, not significant.

A

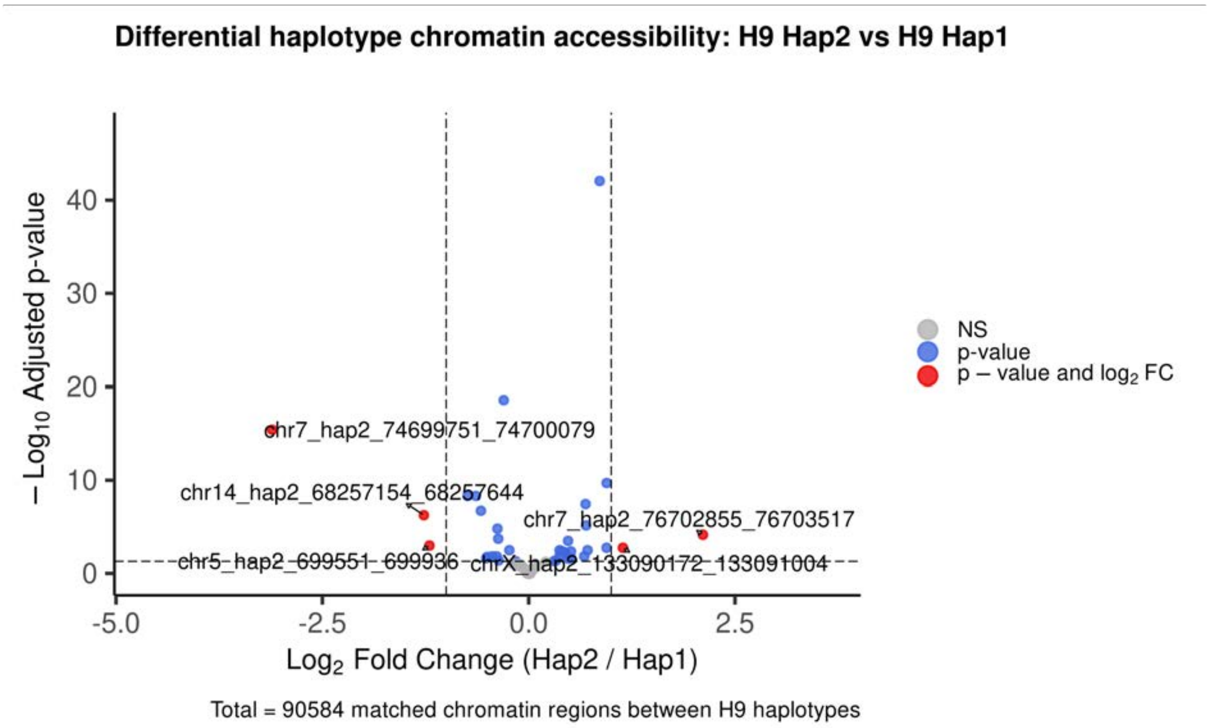


B

C

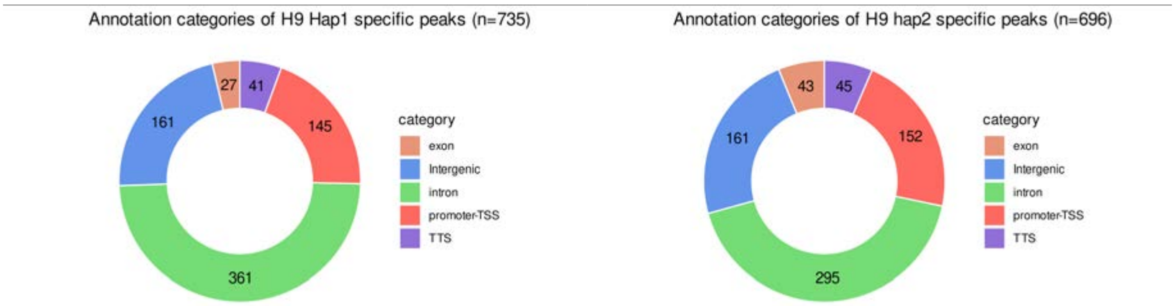


**Figure S25**. Chromatin accessibility analysis in H9 cells undergoing early neuronal differentiation. (**A**) Volcano plot displaying the log2 fold change and statistical significance [-log10(padj-value)] of differential chromatin accessibility between H9 haplotype 2 and haplotype 1. Significantly differential accessible chromatin regions (p-adj < 0.05, |log2FC| > 1) are labelled with haplotype 2 coordinates. (**B**) Donut chart illustrating the genomic distribution of H9 haplotype 1-specific ATAC-seq peaks. (**C**) Donut chart illustrating the genomic distribution of H9 haplotype 2-specific ATAC-seq peaks.
